# Integrated map of somatic mosaicism across human tissues in 25 individuals

**DOI:** 10.64898/2026.09.01.748636

**Authors:** The Somatic Mosaicism across Human Tissues Network, Fritz J. Sedlazeck, Tim H.H. Coorens, Peter J. Park, Andrew B. Stergachis

## Abstract

Although all cells in the body descend from one genome, they accumulate distinct genetic and epigenetic changes over a lifetime, producing a mosaic of somatic variation that can shape development, aging, and disease. This mosaicism is often studied in isolation, leaving unclear how these forms relate within and between individuals. Here, we present the first integrated analysis of the Somatic Mosaicism across Human Tissues (SMaHT) Network’s production resource, profiling up to 20 tissues from 25 donors using short- and long-read, duplex, single-cell, transcriptomic, and epigenomic sequencing, alongside donor-specific near-telomere-to-telomere assemblies. Somatic mutation burden cannot be captured by a single data type or metric, as tissues accumulate distinct variant classes largely independently of one another. Long-read and single-cell data resolved cell-type-specific mutational processes, traced mobile element insertions to source loci, and revealed the developmental timing and functional consequences of individual mutations. Donor-specific assemblies uncovered elevated mutation rates within centromeres and segmental duplications inaccessible to standard reference genomes, while haplotype-resolved chromatin and methylation data showed that nongenetically-deterministic epigenetic states are pervasive across tissues. Together, these findings provide an integrated, multi-scale portrait of somatic mosaicism across the human body, establishing a baseline against which its contributions to aging and disease can be measured.

## Introduction

Although the DNA in all cells within an individual can be traced back to one inherited genome, each cell does not remain genetically identical. Throughout the lifespan, individual cells accumulate changes to their genomes and epigenomes via replication and repair errors, endogenous and environmental DNA damage, chromosome missegregation, mobile element activity, and failures of epigenetic maintenance, such that the body becomes a mosaic of distinct clones^1,2^. Across a lifetime, these may amount to more than 10^15^ somatic mutations per individual, spanning somatic single-nucleotide variants (sSNVs), small insertions and deletions (sIndels), mobile element insertions (sMEIs), structural variants (sSVs), copy number variants (sCNVs), whole-chromosome changes, and DNA methylation and chromatin epimutations^1^.

Somatic mosaicism is now recognized as a fundamental feature of healthy tissue that influences development, aging, and diverse nonmalignant diseases^3,4^. Over the past decade our understanding of somatic mosaicism has rapidly advanced, but largely in parallel, with each new understanding emerging from a distinct subfield defined by its own variant class, technology, tissue, or disease cohort. Specifically, different sequencing methods across various tissue cohorts have shown that somatic mutations occur throughout development and accumulate with age at roughly linear, tissue-specific rates^5–9^, with distinct mutational signatures linking specific environmental exposures and repair processes to the variants they leave behind (*e.g.,* ultraviolet [UV] light in skin)^10^.

These approaches have shown that normal tissues are patchworks of expanding clones driven by diverse somatic alterations^11–13^, including mutations in cancer-associated genes^14,15^. DNA sequencing of blood samples has shown associations between clonal hematopoiesis and age and disease risk^16,17^. Post-zygotic variants restricted to a subset of cells can also produce limited or full presentations of typically inherited conditions. These events are likely underrecognized, as they can escape detection when clinical testing relies on a single, often unaffected, tissue^3^. Furthermore, somatic variation provides insights into early development and late clonal evolution, leveraging each cell’s variation as a unique ‘barcode’^18–22^.

However, how these many forms of somatic mosaicism relate to one another within or across individuals remains unresolved. Consequently, fundamental questions remain open: what does the full landscape of somatic mosaicism look like within a single human body; how much do these patterns vary between individuals; do different classes of somatic variation vary across tissues in concert or independently; how many of a person’s developmental mutations remain recoverable from clinically accessible tissues; which somatic mutations have functional consequences; and how much have we been missing by restricting ourselves to short reads, incomplete genome references (*i.e.,* GRCh38), or bulk assays alone?

The Somatic Mosaicism across Human Tissues (SMaHT) Network was established by the NIH Common Fund to systematically map somatic variation across the human body and to develop the technologies required to detect it^1^. Its first phase benchmarked the sequencing and analytical methods needed for this task^23^. Here, we present the first analysis of the growing SMaHT production resource, which was designed specifically to overcome our existing fragmented view of human somatic mosaicism.

We profiled up to 20 tissues from each of 25 donors, sampling the three embryonic germ layers, gonadal tissues, and clinically accessible tissues, such as blood, skin fibroblasts and buccal swab, across a broad range of adult ages (**Fig. 1a**). We applied an array of orthogonal technologies to each tissue sample, including deep short-read and long-read genome sequencing, ultrasensitive single-molecule (duplex) sequencing, transcriptome sequencing, and CpG methylation profiling. In addition, tissue samples from select donors also included single-cell genome sequencing, chromatin profiling, and near-telomere-to-telomere (T2T) donor-specific assemblies (DSAs). This multi-technology approach enabled in-depth portraits of individual donors, illustrating the range of somatic mosaicism across each body and the putative functional consequences of individual variants. Building on this, we define tissue- and cell-type-specific patterns of mosaicism, including mutational signatures, tandem-repeat expansions, mobile element insertions (MEIs), predicted functional effects, and somatic epimutations, and resolve cell-type contributions using paired single-cell and functional data. We map the extent of clonal expansion and blood-derived cell infiltration across solid tissues; quantify early developmental variants recoverable from clinically accessible samples; exploit DSAs to discover somatic variants in regions inaccessible to GRCh38, including centromeres; and map somatic epimutations of DNA methylation and chromatin. These data and analyses provide an integrated portrait of somatic mosaicism across the human body and establish the foundations of a reference baseline for investigating how somatic variation changes across tissues, aging, and disease.

**Figure 1.**
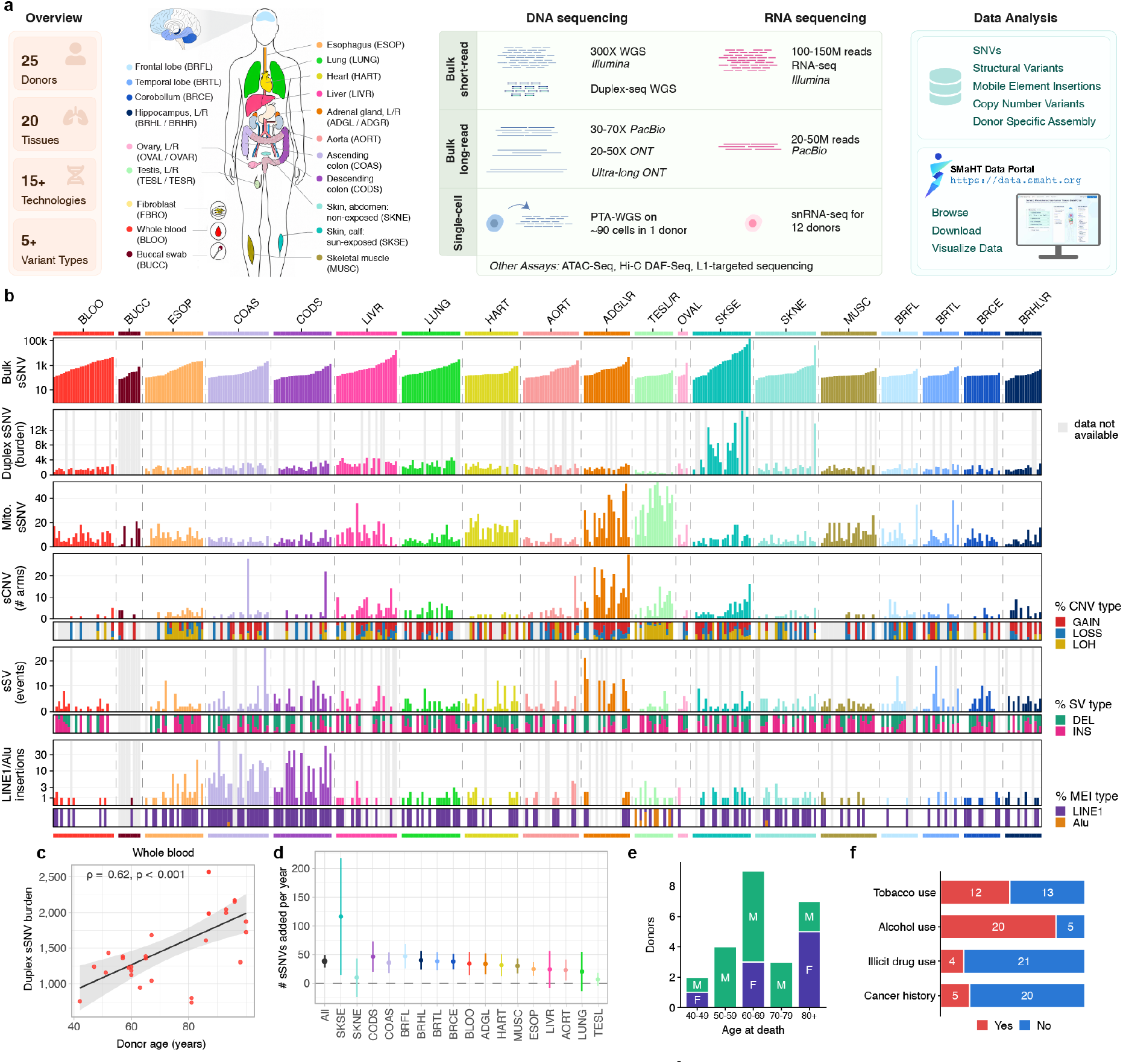
Somatic mutation burden by tissue and variant class across 25 donors. **a,** Design of the Somatic Mosaicism across Human Tissues (SMaHT) Network study. **b,** Somatic mutation burden per donor-tissue pair. Each column represents one donor-tissue pair; all 381 pairs are shown in every row, grouped by tissue and ordered by ascending bulk sSNV burden, with column order conserved across rows. From top: sSNVs from bulk Illumina WGS and PacBio HiFi where available (log scale); duplex-sequencing sSNV burden, with burdens reported per diploid genome (6.2×10^9^ bases) to correct for differences in interrogated duplex bases; mitochondrial sSNVs; number of chromosome arms overlapped by a sCNV; number of sSV events; and number of sMEI events (log scale). Lower sub-panels of the sCNV, sSV, and sMEI rows give subtype composition per pair (0-100%): gain, loss, and CN-LOH for sCNVs; deletion (DEL) and insertion (INS) for SVs; and L1 and Alu for sMEIs. Grey, data not obtained. **c,** Duplex sSNV burden versus donor age for whole blood. Each point is one donor-tissue-assay measurement pooled across four duplex assays (CODEC, CompDuplex-seq, META-VISTA-seq, NanoSeq-MBN); grey line ordinary least-squares fit with 95% confidence band. ρ and p are Spearman rank correlation coefficients and their asymptotic p-values. **d,** Burden of sSNV accumulation by tissue, given as the slope of duplex sSNV burden on donor age (ordinary least squares) with 95% confidence intervals; the leftmost grey point (All) is the cohort-wide fit across all tissues. **e,f,** Donor metadata. (e) age at death, stratified by sex. (f) donors reported as positive (red) or negative (blue) for tobacco use, alcohol use, illicit drug use and prior cancer.

## Results

### Somatic mutational landscape across 381 tissues from 25 donors

To establish an integrated map of somatic mosaicism across human tissues, we enrolled through next of kin 25 postmortem donors (median age at death of 66 years, range 42-89+, with 16 males and 9 females) (**Fig. 1a**) and collected up to 20 tissue types across each donor (381 total tissue samples, median 16 tissues per donor). From each tissue sample, we obtained multiple small tissue punches, or “cores,” and applied a range of sequencing technologies across these cores (**Fig. S1 and Table S1**). Two cores per tissue underwent short-read whole-genome sequencing (WGS) (combined median coverage 368x). Despite the challenge of obtaining high-molecular-weight (HMW) DNA from postmortem tissues, we were able to also profile the majority of samples with long-read genome sequencing using Oxford Nanopore Technologies (ONT) (n=258, median coverage 63x) and/or PacBio HiFi (n=237, median coverage 41x), with 30% of the PacBio HiFi samples profiled by Fiber-seq^24^. We additionally applied four distinct duplex DNA sequencing approaches (n=294), alongside short- and long-read transcript sequencing (n=190 and n=115, respectively). Further assays, including single-nucleus RNA-seq (snRNA-seq), single-nucleus ATAC-seq (snATAC-seq), Hi-C, and ultra-long ONT sequencing (UL-ONT), were applied to a subset of samples. For donor SMHT005, we additionally performed single-cell genome sequencing using primary template-directed amplification (PTA) on 91 cells across 7 tissues. All data (>200 terabytes), along with full metadata and processing details, will be available through the SMaHT Data Portal (http://data.smaht.org).

Integrating the paired short- and long-read data from each tissue sample revealed extensive somatic variation across every variant class examined (**Fig. 1b and Fig. S2**). Bulk WGS identified 536,127 sSNVs, with sun-exposed calf skin (SKSE) alone accounting for 56% of all calls. Excluding SKSE, median burden was 3,608 sSNVs per donor (IQR 2,304-7,120), predominantly at low variant allele fraction (median 3.4%; 12% below 2% variant allele frequency [VAF]), reflecting the added sensitivity of high-coverage WGS. After SKSE (median 1,650.0 sSNVs per sample), burden was next highest in whole blood (832.0), liver (514.0), and adrenal gland (509.0), and lowest in testis (136.5), muscle (140.0), and brain (147.0-157.0). Whereas bulk WGS preferentially captured larger, often earlier-arising clones, duplex sequencing detected a largely nonoverlapping population of sSNVs (552,270) reflecting low-frequency mutations largely invisible to bulk sequencing approaches. As expected, duplex sSNV burden increased with donor age in whole blood (ρ=0.62, p<0.001; 34.2 sSNVs/year), as well as across other tissues, rising fastest in sun-exposed skin (111.0 sSNVs/year), brain frontal lobe (31.7), and descending colon (30.4), and slowest in testis (4.4) (**Figs. 1c, 1d**).

We identified 501 unique mitochondrial DNA (mtDNA) sSNVs with a median of 72 per donor (IQR 55–85) (**Fig. 1b**). Most occurred at low heteroplasmy levels (77.9% <2% VAF). Transitions substantially outnumbered transversions (Ti/Tv = 6.32), with an excess of C>T and A>G substitutions.

This spectrum is more consistent with replication-associated deamination rather than oxidative damage as the predominant source of mtDNA sSNVs^25,26^. In contrast to nuclear sSNVs, mtDNA sSNV burden showed only a modest association with donor age (p=0.12; +0.4 sSNVs/year). Mutation burden varied markedly across tissues, with the highest values in testis (32.6 sSNVs per sample) and adrenal gland (22.1), and the lowest in hippocampus (4.0).

Beyond point mutations, we characterized somatic structural variation at multiple scales. Clonal sCNVs, owing to their large genomic footprint, can be sensitively detected from bulk sequencing at cell fractions below 1%^27,28^, and we identified 1,008 autosomal sCNVs (318 gains, 303 losses, 281 copy-neutral losses of heterozygosity (CN-LOH), and 106 unclassified events) at a median cell fraction of 1.02%. Like nuclear-encoded sSNVs, sCNV burden increased with age (ρ=0.72, p=5.7×10^-5^) and was highly tissue-specific, with the highest rates in adrenal gland (6.2 sCNVs per core) and testis (3.5 sCNVs per core, mostly CN-LOH, with 0.5% median cell fraction). In both tissues, events occurred broadly across the genome, driven by distinct processes (i.e., accumulated aneuploidy in adrenal gland and CN-LOH in testis). In contrast, other tissues showed locus-specific sCNVs, such as 9q CN-LOH, which was observed in the esophagus of 12 of 24 donors and is known to act as a second hit to *NOTCH1* mutations^14,29^.

In addition to these sCNVs, long-read sequencing further resolved 970 sSVs (median 25 unique sSVs per donor), split between deletions (57.6%) and insertions (42.4%). These sSVs (50 bp-10 kbp) occurred largely within tandem repeat regions (63.5%), and an additional 9.8% belonged to other known sSV classes^30,31^ (*e.g.,* duplications). Due to the long-read coverage heterogeneity (6x to 145x per-sample), only sSVs that occurred at ≥5% VAF in ≥1 sample are reported. With this consistent VAF floor, no correlation between sSV burden and age was observed (PearsonR 0.00, p > 0.05), potentially reflecting an early developmental origin of these sSVs, and the highest average sSV burden was observed in adrenal glands (nine donors) with 7.9 sSVs per 50x sequencing coverage.

Long-read data were similarly critical for resolving sMEIs. Using a multi-platform integration approach^32^, we identified 790 sMEIs representing 718 unique insertion sites, predominantly LINE-1, with only 13 Alu insertions. Because long reads capture both the inserted sequence and the hallmarks of target-primed reverse transcription (i.e., poly(A) tails and target-site duplications) within the same read, they enabled confident sMEI identification even from a single supporting read. Somatic retrotransposition was detected in nearly all tissues, including tissues previously not known to harbor sMEIs, such as adrenal gland, skin, aorta, muscle, ovary, and whole blood. sMEIs showed tissue-specific burdens, with colon and esophagus together accounting for 76.3% of all sMEIs, and brain frontal lobe and whole blood having the lowest burdens, patterns that mirror cancer sequencing data, in which somatic L1 activity is high in gastrointestinal cancers but low in brain tumors and hematologic malignancies^33,34^.

To provide donor-level context for these somatic findings, we performed germline screening, which identified four donors carrying heterozygous pathogenic or likely pathogenic variants in established cancer predisposition genes: one donor carried a *BRCA2* variant, which acts dominantly, and three were monoallelic carriers of the recurrent *MUTYH* allele, which typically confers risk only in a biallelic state. Additional germline variants were identified, including a heterozygous pathogenic *KCNQ1* variant associated with long QT syndrome 1 (**Table S2**). Because best practices for returning genomic findings, including consent, variant confirmation, genetic counseling, and clinical follow-up, remain far less established for deceased-donor studies than for living-donor cohorts^35^, the SMaHT Network determined in advance that results would not be returned and stated this explicitly in the next-of-kin authorization.

We additionally collected extensive phenotypic data from each donor, including age range, clinical features such as cancer history, and exposures to tobacco, alcohol, and illicit drug use, (**Figs. 1e, 1f**). Together with the germline annotations, these data provide the context required to interpret differences in somatic burden between individuals. Comparing bulk-sequence-derived sSNV burden across the same tissues from different donors allowed us to test whether some individuals are inherently more or less prone to somatic mutation. Pairwise comparisons were performed across the donor-tissue matrix, excluding outlier donor-tissue pairs with disproportionately high sSNV counts (Methods). Overall, although three donors (SMHT029, SMHT020, and SMHT040) harbored a higher sSNV burden in at least six of their tissues, no donor showed a significantly elevated sSNV burden across all of their tissues. Given the number of potential confounders (e.g., exposure history, blood infiltration, and germline variation) a full elucidation of donor-level trends will require the larger, planned SMaHT cohort.

### A single-donor portrait of somatic mosaicism across tissues and cell lineages

Having established the overall landscape of somatic mosaicism across the cohort, we next leveraged our multi-tissue sampling design to compare somatic mutation burden across many tissues from the same individual, controlling for germline background, age, and exposure history. We illustrate this with donor SMHT005, a 59-year-old male profiled across 18 tissues (12 with paired long- and short-read data).

The tissue with the highest burden differed markedly depending on which variant class was considered (**Fig. 2a**). sSNV burden was highest in sun-exposed skin by both bulk and duplex sequencing, consistent with UV exposure. sMEIs were near-exclusively L1 insertions in the colon, which accounted for 45 of 51 events, split between ascending (n=32) and descending (n=13) segments. Chromosome-arm-level sCNVs were dominated by adrenal gland, with 25 events compared with 1-7 in every other tissue. Mitochondrial sSNVs were most numerous in testis, which carried 32 of 89 distinct variants. No tissue ranked highest for more than one variant class, indicating that somatic variant burden is decoupled from variant class within a single individual.

**Figure 2.**
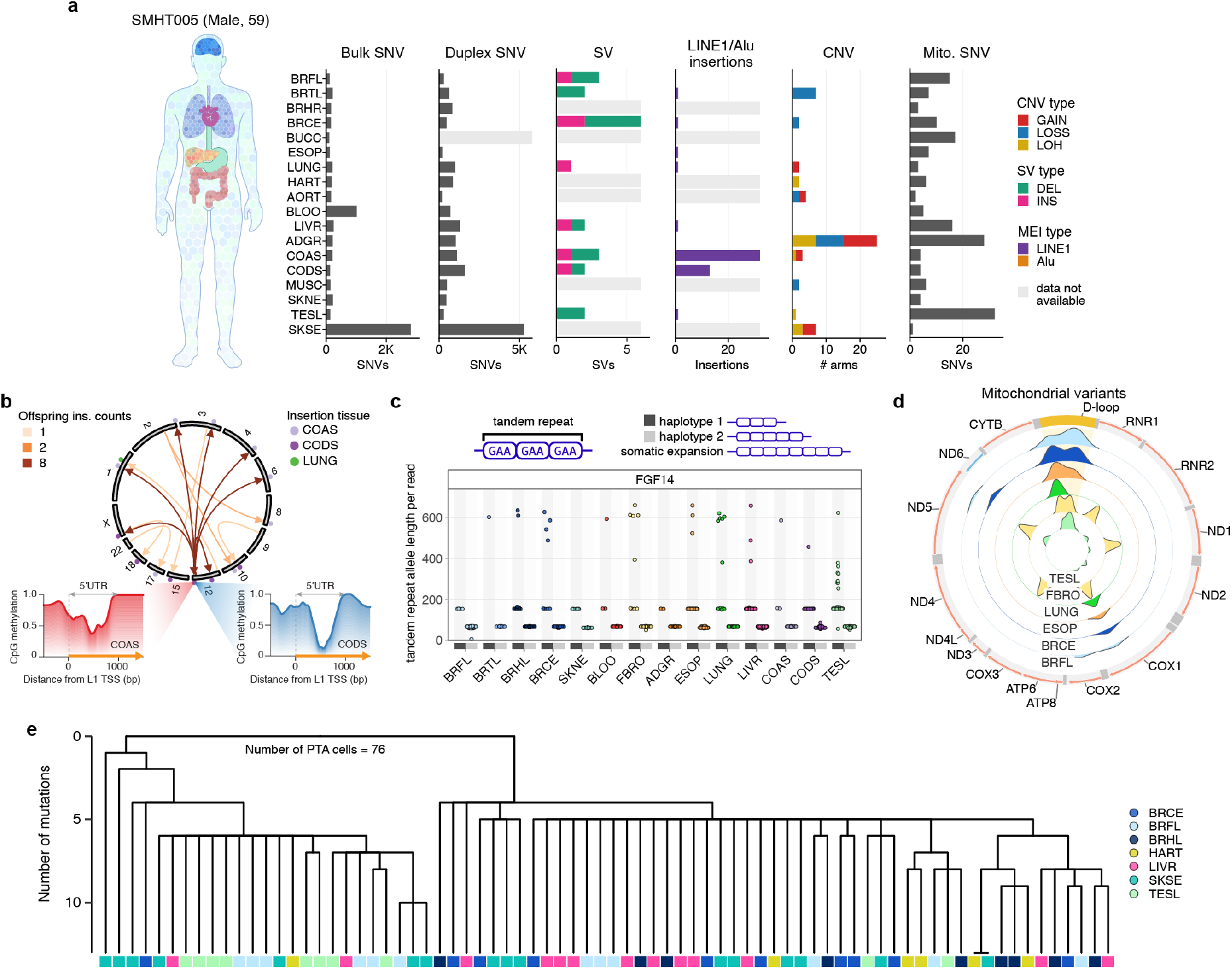
Somatic mutation burden varies across tissues within a single individual, SMHT005. **a**, Somatic variant burden across 18 tissues from donor SMHT005 (male, 59 years). Left to right: bulk sSNVs, duplex sSNVs, sSVs colored by type (DEL, INS), sMEIs colored by type (L1, Alu), sCNVs plotted as the number of affected chromosome arms and colored by type (GAIN, LOSS, LOH), and heteroplasmic mitochondrial sSNVs. Bulk and duplex sSNV counts are both genome-wide and are shown as raw, uncorrected calls. sSVs and sMEIs were called from long-read data, with sMEIs additionally called using the capture-based TEnCATS method (Methods). Light grey bars indicate tissues lacking the corresponding assay (long-read for sSVs and sMEIs, duplex sequencing for duplex sSNVs). **b**, Circos plot showing source L1 elements and their offspring somatic insertions. Arrow color intensity indicates the number of offspring generated by each source element, and colored dots denote the tissues in which the insertions were detected. The bottom panels show CpG methylation profiles across the promoter region of the most active source element in SMHT005. The 5′ UTR spans the first 1 kbp from the L1 transcription start site (TSS), and the L1 element is indicated by the yellow arrow. Blue and red lines represent long-read CpG methylation profiles derived from long-read sequencing of the descending colon (ONT) and ascending colon (PacBio HiFi), respectively. **c**, Tandem repeat instability affecting an intron in *FGF14* in donor SMHT005. The y-axis shows the haplotype-resolved TR length (bp) genotyped from long-read data, with each point representing one read. Shown is a (GAA)n TR in the first intron of *FGF14*, with stable alleles of ∼68 and 154 bp in length observed in all tissues, alongside unstable and somatic alleles in several tissues. A third allele, ∼313 bp in length, is mostly detected in testis. **d**, Heteroplasmic mtDNA sSNV distribution (VAF 0.1-95%) across six tissues from donor SMHT005. Outer tracks show gene and D-loop annotation along the mitochondrial genome; inner concentric tracks show per-tissue heteroplasmic sSNV distribution (color-coded by tissue types). **e**, Cell-lineage reconstruction from 76 PTA-amplified single cells from seven tissues of donor SMHT005 (cerebellum, frontal lobe, hippocampus, heart, liver, calf skin, and left testis). Fifty-two clonal sSNVs identified by bulk WGS of the corresponding tissues were genotyped in the PTA single-cell genomes and used for lineage reconstruction. Tips represent individual cells and are colored by tissue of origin. Terminal branches were truncated at 13 mutations to emphasize the early phylogeny.

Given this concentration of somatic L1 insertions in colon, we next sought to identify their source elements (i.e., the full-length L1 loci from which these somatic insertions originated). Long-read sequencing proved substantially more powerful for this than conventional approaches, as using source-specific sequence variants captured by long reads within L1 insertions^32^ enabled us to trace 15 somatic L1 insertions to seven full-length source elements, compared with only two insertions traced by conventional methods relying on 3′ transductions (**Fig. 2b**). Notably, a source element on chromosome 12 gave rise to eight somatic insertions without transduction across the two colon tissues. This polymorphic element is absent from the GRCh38 reference genome and was only recently reported through long-read cancer genome profiling^36^, underscoring the value of long-read, reference-independent detection approaches. Long-read CpG methylation profiling at this source element exposed colon-specific hypo-CpG methylation of their 5′ untranslated region (UTR) promoters, consistent with epigenetic derepression potentially driving their transcriptional activation and subsequent retrotransposition. Long reads were similarly indispensable for resolving somatic variation at tandem repeats (TRs), which are a major source of both inherited and somatic genome variation^37^ that can often only be accurately genotyped using reads long enough to span the full repeat tract. We assessed ∼600,000 TR loci per donor, classifying each by repeat-length variability and evidence of somatic instability (**Fig. 2c**). For example, at a (GAA)n TR within the *FGF14* locus, both haplotypes were largely stable, but haplotype 1, which harbored an expanded germline GAA repeat, showed a small number of reads with either contractions or further expansions, including some approaching the germline pathogenic range^38^. By contrast, in testis, reads from haplotype 1 formed two distinct clusters, one at the apparent germline expanded GAA repeat length, and one at roughly half that length, a discrete tissue-restricted contracted allele that potentially reflects an early developmental somatic contraction. These illustrate that TR mosaicism can result in extensive length heterogeneity accruing on a single parental haplotype and can be confined to a subset of cells or tissues.

The mitochondrial genome also showed tissue-restricted somatic heteroplasmy. Across six SMHT005 tissues assessed for mitochondrial sSNVs, mitochondrial heteroplasmies spanned a VAF of 0.1-95% and preferentially localized to the D-loop, the major mitochondrial noncoding control region. The D-loop accounted for 63.3% of heteroplasmic sSNVs, with these sSNVs consistently detected across multiple tissues, a 9.36-fold enrichment over the full mtDNA (**Fig. 2d**). Outside the D-loop, heteroplasmic sSNVs were largely tissue-specific, most strikingly in testis, which harbored the largest burden of heteroplasmic sSNVs, dispersed broadly across the mitochondrial genome. However, one heteroplasmic sSNV (*COX1* m.7056G>A) was shared by all six tissues at similarly low VAFs (1-5%), indicating this mutation was either inherited or arose early in development and persisted across lineages. Together, these findings reveal an uneven burden of mitochondrial heteroplasmy across both development and the mitochondrial genome itself.

While mutation sharing across tissues offers an indirect window into developmental timing, directly reconstructing when and how somatic lineages diverge requires tracing individual cell genomes. To do this, we performed lineage reconstruction from 76 PTA-amplified single-cell genomes across seven tissues from SMHT005 (**Fig. 2e**). This reconstruction leveraged sSNVs detected by paired bulk WGS, which captures early embryonic mutations shared by many cells within a tissue^18,39^, resulting in a phylogeny that reflects only the early portion of development (**Fig. S3**). The phylogeny resolved into two major early lineages, with cells from the seven tissues contributing to each in markedly different proportions. Quantifying these contributions using bulk sequencing across 16 tissues, we found, consistent with prior observations^18^, one lineage dominated most somatic tissues, contributing ∼91% of cells in cerebellum, with ascending colon and testis showing nearly balanced contributions from both lineages (46-52% and 40-54%, respectively) (**Fig. S4**). This balanced contribution in testis is consistent with the hypothesis that the germline is founded by two cells, one from each of the first two developmental lineages^40^. Beyond resolving lineage history itself, these tissue-specific lineage proportions help explain the origin of individual somatic variants in this donor. This makes it possible to interpret the observed VAF of any sSNV, sSV, or sMEI in SMHT005 against the expected contribution of its lineage of origin, rather than in isolation. For example, the discrete contracted *FGF14* GAA TR on haplotype 1 in testis is suggestive of having arisen in one of the two founder cells that gave rise to the testis lineage. Together, these analyses indicate that even within a single donor, somatic mosaicism is shaped by distinct processes operating at different scales and timescales, from tissue- and mutation-type-specific burden, to the developmental origin and lineage restriction of individual variants, to the earliest cell divisions of the embryo itself.

## Mutational processes underlying tissue-specific somatic variation

We next sought to explore how tissue-specific patterns of somatic mosaicism could be connected to distinct mutational processes. We found that 55% of each donor’s sSVs were within TR regions and 52% (22/42) of TR-overlapping sSVs fell within gene bodies, suggesting a possible relationship between transcription and somatic instability. Among these TRs, seven showed recurrent patterns of somatic instability across donors (**Table S3**). For example, a TR within intron 1 of *SLC10A6* (**Fig. 3a**) showed somatic instability in calf or abdominal skin in 50% (12/24) of the donors, with unstable reads averaging 125 bp longer than the presumed germline allele and reaching a mean VAF of 12.6% (range 6-22%). By comparison, instability was observed in nonskin tissues in only four donors, with a mean length deviation of 38 bp and mean VAF of 3.4%. Paired RNA-seq from these samples showed substantially higher *SLC10A6* expression in skin than in other tissues (TPM ∼18.2 vs. ∼1.04; **Fig. 3a**), suggesting that recurrent somatic TR instability at this locus coincides with tissue-elevated transcription.

**Figure 3.**
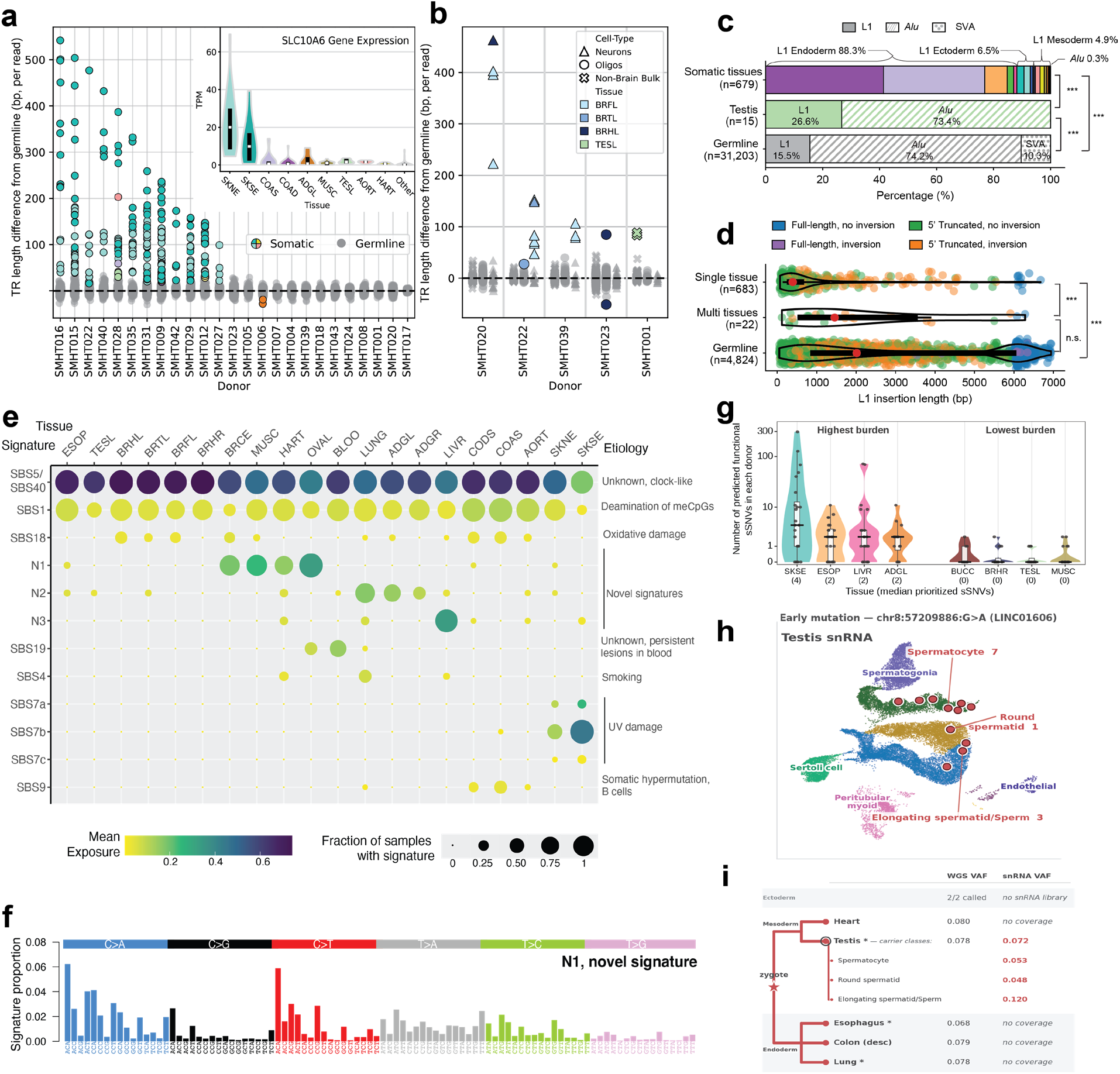
Tissue- and cell-type-specific patterns of somatic mutations. **a,** Scatter plot of TR length deviation (bp) from germline allele length per read for intron 1 of *SLC10A6*. Points represent individual reads. Reads with somatic alterations in TR length are colored according to inset. Reads with TR lengths mirroring the presumed germline length are colored in grey. Inset: RNA-seq expression (transcripts per million, TPM) by tissue across donors. **b,** Scatter plot of TR length deviation (bp) from presumed germline length in *CHST8* phased to specific cell types, showing a neuron-specific expansion. **c,** Stacked bar plots showing the relative proportions of L1, Alu, and SVA insertions among tissue-specific somatic insertions, testis-specific insertions, and non-reference germline insertionsfrom the 1000 Genomes Project (***p < 0.001, chi-squared test). Solid, hatched, and dotted patterns represent L1, Alu, and SVA insertions, respectively. Colors indicate the tissues in which tissue-specific somatic insertions were detected. **d,** Violin and scatter plots showing the length distributions of L1 insertions detected in a single tissue, multiple tissues, or the germline. Individual insertions are colored according to their structural features, including 5′ truncation and 5′ inversion, and red points indicate median insertion lengths. ***p < 0.001, Wilcoxon rank-sum test. **e,** Bubble plot showing the relative contribution of each mutational signature across tissues based on duplex sequencing data; color indicates mean contribution, and size indicates the fraction of samples with an estimated lower 95%-confidence interval contribution of at least 1%. Signatures include the COSMIC reference signatures and novel signatures N1-3. **f,** Bar plot of the 96-trinucleotide distribution of N1, the novel signature identified in muscle, heart, ovary and cerebellum. **g,** Violin plots and box plots (interquartile range and median) of the number of unique functional sSNVs predicted by ≥1 deep learning model for each donor-tissue pair (axis is logscaled), excluding tissues with fewer than four donors. **h,** UMAP of snRNA-seq data from testis cells from donor SMHT042. Red dots show cells that harbour the identified developmental sSNV in *LINC01606* (11 of 171 nuclei with reads at this position). **i,** Diagram of profiled tissues, coverage of the *LINC01606* locus, and presence of the mutation, showing that although the *LINC01606* sSNV is present across tissues by WGS, its apparent absence in other tissues profiled by snRNA-seq is explained by testis-specific expression of the gene.

We next evaluated whether any of these TRs show cell-specific somatic instability by combining repeat-spanning long reads with cell-type-associated CpG methylation patterns along the same read. This approach identified a recurrent, neuron-specific TR expansion within *CHST8* (**Fig. 3b**) in brain samples from four donors. Specifically, neuron-assigned reads showed expansions averaging ∼480 bp longer than the germline allele, while non-neuronal reads from the same tissue largely remained near germline length. This expansion was most pronounced in neocortical regions (i.e., frontal lobe, temporal lobe, and hippocampus) and comparatively minimal in cerebellum, suggesting regional specificity within the neuronal population itself, potentially reflecting the von Economo neuron-specific expression of *CHST8*^41^. SMHT001 additionally showed expansion of the same repeat in testis (**Fig. 3b**). Together, these observations show that somatic TR instability can arise predominantly within a specific cell population and can differ across anatomical regions of the same organ.

Tissue specificity also extended to sMEIs. As with TRs, read-level CpG methylation resolved the cellular origin of sMEIs, identifying twelve cell-type-specific events: five in skin localizing to dermal fibroblast or epidermal keratinocyte signatures (**Fig. S5**), three in brain localizing to neuron or oligodendrocyte signatures, two in heart localizing to cardiomyocyte or fibroblast signatures, one in aorta localizing to endothelial or smooth-muscle cell signatures, and one in liver localizing to hepatocyte or macrophage signatures. Across non-gonadal tissues, L1 elements accounted for 99.7% of sMEI insertions, consistent with prior observations in cancer^33,36^. In contrast, testis-specific insertions were predominantly Alu (73.4%), mirroring the spectrum of polymorphic germline MEIs, which are predominantly Alu elements^31^ (**Fig. 3c**). The structure of somatic L1 insertions also differed according to their tissue distribution: insertions restricted to a single tissue (i.e., likely arising later in development) were shorter and predominantly 5′ truncated (median=393bp), whereas insertions observed across multiple tissues from the same donor (i.e., likely arising early in development) were longer (median=1,456bp, p<0.001), approaching germline insertion lengths (median=2,014bp) (**Fig. 3d**). These patterns support a model whereby developmental-stage-dependent differences in retrotransposition, host surveillance, or DNA repair shape the structure of somatic L1 insertions.

We examined tissue-specific patterns of sSNVs using duplex sequencing across 294 tissue samples from 21 donors, applying CODEC (n=123)^42^, CompDuplex-seq (n=153)^43^, NanoSeq (n=160)^44,45^, or META-VISTA-seq (n=34)^43^ (Methods). Excluding two mutational signatures with a possible technical origin, we extracted ten known COSMIC reference signatures and three novel signatures (**Fig. 3e and Fig. S6b**). All tissues harboured the imprints of SBS1, caused by deamination of methylated cytosines, and SBS5/SBS40, of unknown etiology, that accumulate in a clock-like manner in both normal tissues and cancers^46^. Many tissues also bore the imprint of SBS18, which has been attributed to oxidative damage^10^.

Tissue-specific signatures included those reflecting extrinsic mutagens: SBS7a-c (UV damage) were detected in skin, with the expected elevation observed in sun-exposed sites (**Fig. S7a**); SBS4 (tobacco smoke) was detected in lung and, to a lesser extent, heart, and strongly enriched in donors with a known smoking history (**Figs. S7b, S7c**). Other tissue-specific signatures may be of endogenous origin. SBS19, of unknown etiology but linked to persistent DNA lesions^47^, was present in blood. SBS9, attributed to polymerase eta-induced somatic hypermutation, was identified in solid tissues, predominantly colon, suggesting the presence of tissue-resident B lymphocytes^48^.

We further identified three residual, novel signatures not represented in the reference database (“N1”, “N2” and “N3”, **Fig. 3f and Fig. S6b**). N1 is prevalent in heart, ovary, muscle, and cerebellum, and increases linearly with age (p=8.04×10^-8^, Pearson correlation test, **Fig. S7d**), suggesting a previously unobserved endogenous mutational process. These tissues and cell types are generally post-mitotic, polyclonal, and rarely give rise to tumors, features that may explain why these signatures were not detected in previous studies relying on normal cell clones^6^ or cancer genomes^10^. N2 was mainly identified in lung and adrenal gland samples, increased with age, and was more pronounced in smokers independently of age (**Fig. S7e,** p=3.34×10^-5^). N3 was strongly enriched in liver but not age-dependent (p=0.959, Pearson correlation test), suggesting this signature may be due to an exposure.

### Functional consequences of sSNVs

We assessed the functional consequence of the identified somatic genetic variants at two levels: first, by systematically predicting the molecular impact of sSNVs genome-wide, and second, by using single-cell profiling to trace individual mutations to their cellular and developmental origins. Across all donors, 5,922 sSNV observations (representing 4,968 unique sSNVs) overlapped annotated protein-coding sequence, and 339 sSNVs (279 unique) were predicted to be stop-gained variants, affecting 68 distinct high-pLI genes (pLI > 0.9), such as *TP53* and *NOTCH1* (**Table S4**). More broadly, based on all VEP consequence annotations, 7.9% of sSNVs (42,115/536,127) carried a potential functional consequence (highlighted in **Fig. S8**). Within this set, 25,071 sSNVs were annotated to regulatory regions, 5,340 fell in UTRs, and 965 at splice sites, among other classes. To predict which of these variants are likely to be functionally consequential, we applied four complementary models (SpliceAI, PromoterAI, AlphaGenome, and APARENT2) spanning splicing, promoter activity, tissue-specific expression, and polyadenylation, respectively (Methods). This identified 1,156 unique sSNVs (0.25% of all sSNVs) with predicted functional impact supported by at least one model: 578 predicted to alter canonical splice donor or acceptor site usage (SpliceAI delta score >0.2), 320 predicted to alter promoter activity (PromoterAI >95^th^ percentile), 240 predicted to affect tissue-specific gene expression (AlphaGenome >95^th^ percentile), and 80 predicted to affect polyadenylation site usage (APARENT2 >95^th^ percentile) (**Fig. S9 and Table S5**). Tissues with high sSNV burden, such as sun-exposed skin, esophagus and liver, had the highest number and proportion of sSNVs predicted to be functionally consequential (**Fig. 3g and Fig. S10**). However, this burden was not enriched among high-VAF sSNVs with predicted-functional sSNVs distributed similarly across the full range of observed VAFs (**Fig. S9**). This indicates that the potential for functional consequence is a pervasive feature of somatic mosaicism.

The consequence of a somatic mutation depends on the variant’s molecular impact and where a mutation resides. Thus, we used snRNA-seq and snATAC-seq data to resolve the developmental fate of individual sSNVs, distinguishing early developmental mutations broadly distributed across the body from later mutations confined to a single lineage. For example, the donor SMHT042 harboured a chr8:57,209,886 G>A sSNV within the oncogenic long noncoding RNA, *LINC01606*^49,50^, that was detected by WGS across all mesoderm-, ectoderm-, and endoderm-derived tissues sampled. Because this raised the possibility that it might extend into the germline, we performed snRNA-seq of testis tissue from SMHT042 and confirmed the variant’s presence within the germline lineage across successive post-meiotic stages of spermatogenesis: spermatocytes (VAF 0.053), round spermatids (VAF 0.048), and elongating spermatids/sperm (VAF 0.120) (**Fig. 3h**). The presence of this variant in mature sperm makes it compatible with transmission of somatic mutations to offspring.

Nevertheless, not all mutations have a developmental origin. The donor SMHT006 harboured a chr2:217,870,675 C>T sSNV within *TNS1*, which was detected by WGS only in skeletal muscle (VAF 0.057). snATAC-seq similarly identified the variant in skeletal muscle (VAF 0.056), with all 20 variant-carrying nuclei restricted to skeletal myofibers (**Fig. S11**), a pattern consistent with a later mutation confined to a single differentiated lineage. Together, these examples show that single-cell profiling can resolve not only the cell types harbouring a somatic mutation but also the likely developmental timing and lineage distribution, up to and including the potential for transmission to the germline.

### Clonal drivers and blood infiltration shape spatial heterogeneity in somatic mosaicism

Sequencing multiple cores from the same tissue further allowed us to identify tissue-specific clonal architectures within an individual (**Fig. 4a**). Somatic variants were classified as unique to a single core or shared between adjacent cores. For example, core 2 from SMHT007 abdominal skin contained 6.6-fold more sSNVs than core 1 (36,529 vs. 5,539 specific to core 2 and core 1, respectively, with only 68 shared) (**Fig. 4b and Fig. S12**). This heterogeneity was quantified across all donor-tissue pairs with two independently sequenced cores, with skin, esophagus, and liver all showing the highest sSNV heterogeneity between cores (**Fig. 4c**), suggestive of spatially confined clonal expansion. In contrast, sSNVs from separate cores of cerebellum and muscle were largely concordant, suggesting either less frequent clonal subarchitecture in these tissues or clonal subarchitecture on a spatial scale larger than the separation between sampled cores. As expected, whole blood, a liquid tissue sampled without coring, shared the largest fraction of variants between sequenced samples.

**Figure 4.**
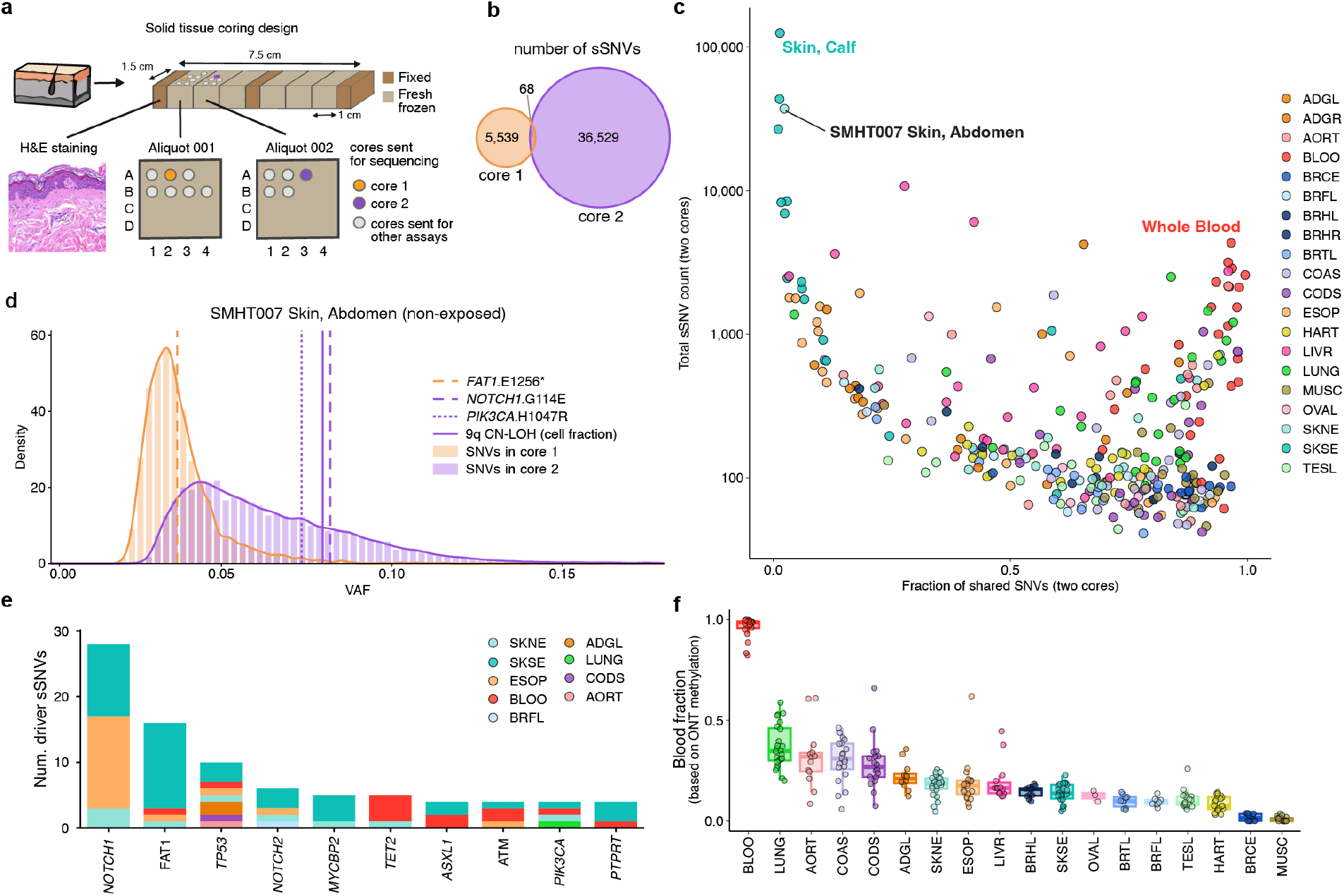
Spatially heterogeneous sSNV burden between adjacent cores of noncancerous tissue. **a,** Solid tissue sampling design, illustrated with non-sun-exposed skin (suprapubic abdomen) from donor SMHT007. Full-thickness tissue blocks (7.5×1.5 cm) were cut into sections that were alternately formalin-fixed (dark brown) or fresh-frozen (tan); formalin-fixed sections were H&E-stained for histological review (bottom left). Fresh-frozen section aliquots were sub-sampled into 3 mm spatially mapped tissue cores arrayed in a 4×4 grid (rows A-D, columns 1-4), which were distributed across assays and sequencing centers (Methods). Data from cores sequenced in panel b are core 1 (tissue aliquot 001, orange) and core 2 (tissue aliquot 002, purple); grey circles denote cores allocated to other assays. **b,** Venn diagram of sSNVs called separately in the two tissue cores from SMHT007 skin (abdomen). Areas of the circles are proportional to the number of sSNVs: 5,539 core-1-specific, 36,529 core-2-specific, and 68 shared. Illumina WGS coverage was 185x (core 1) and 138x (core 2). **c,** Total sSNVs detected at VAF ≥ 0.5% in either core (log scale) versus the fraction of those shared between the two cores, for every donor-tissue pair with two independently sequenced cores (327 donor-tissue pairs from 25 donors). Both cores were genotyped by pileup at every sSNV called in that sample, and the shared fraction is the number of sSNVs with VAF ≥ 0.5% in both cores divided by the number reaching that threshold in at least one. Each point is one donor-tissue pair, colored by tissue as shown in Fig. 1a. Whole blood, a non-solid tissue sampled without coring, is shown as a sharing control. **d,** VAF distributions of sSNVs private to each core in SMHT007 skin (abdomen): histograms with overlaid kernel density estimates, core 1 in orange and core 2 in purple. Dotted vertical lines mark VAFs of the putative driver variants: FAT1 p.E1256* (core 1, orange), NOTCH1 p.G114E, and PIK3CA p.H1047R (core 2, purple). The solid purple line marks the cell fraction of the core-2-specific chr9q CN-LOH event. **e,** Number of putative driver sSNVs per gene across all samples, for the ten most frequently mutated cancer genes. Bars are stacked and colored by tissue as shown in Fig. 1a. Variants were classified as putative drivers if predicted high-impact/likely-pathogenic, splice-altering (SpliceAI ≥0.8), or at a known mutational hotspot (cancerhotspots.org, q<0.1in canonical CHIP or IntOGen gene sets (Methods). f, Blood cell infiltration of each tissue sample, estimated by deconvolution of ONT CpG methylation data (Methods). Each point is one sample (n=294 samples from n=24 donors); tissues are ordered by decreasing median. Box plots show the median (center line), first and third quartiles (box bounds) and 1.5x the interquartile range (whiskers)

We next searched for candidate driver mutations putatively underlying these clonal expansions using three catalogues of known cancer and CHIP genes^51–53^ (Methods). The SMHT007 abdominal skin cores carried nonoverlapping candidate drivers in canonical cancer genes: core 1 carried a *FAT1* p.E1256* nonsense variant (3.7% VAF), while core 2 carried a *PIK3CA* hotspot p.H1047R (7.4% VAF) and *NOTCH1* p.G114E (8.2% VAF), with a copy neutral loss of heterozygosity (CN-LOH) of 9q (8.0% cell fraction) potentially acting as the second hit of the *NOTCH1*p.G114E variant (**Fig. 4d**). A 9q CN-LOH event was also present in core 1 at a substantially lower cell fraction (3.4%) but was on the opposite haplotype from the 9q CN-LOH event in core 2, indicating that these events arose independently (**Fig. S13**). Histological examination of adjacent blocks from this tissue showed no evidence for abnormal pathology. Extending this analysis across the full 25 donor cohort revealed multiple recurrent candidate driver variants in known cancer genes, such as *NOTCH1*, *FAT1*, and *TP53*, concentrated in the tissues where clonal expansions are well documented (*i.e.,* skin, esophagus, and blood)^14,15^ (**Fig. 4e**).

Some solid tissues showed a relatively high sSNV burden alongside a large fraction of shared sSNVs between cores, a pattern we suspected was driven by the presence of blood cells within these solid tissues (**Fig. 4c**). To directly quantify the presence of blood cells, we used CpG methylation signals from ONT data to deconvolve the tissue of origin of each read, allowing us to estimate the fraction of blood-derived reads across tissues. By this metric, lung, abdominal aorta, and colon were among the most blood cell-infiltrated solid tissues, whereas heart, brain, and muscle contained the least (**Fig. 4f**). These findings underscore that, unless explicitly accounted for, blood cell infiltration can inflate apparent somatic variant burden in solid tissues and thereby produce a misleading picture of true tissue-intrinsic mosaicism.

### Single-tissue sampling incompletely captures early developmental somatic mutations

As clinical genetic diagnostic testing relies almost exclusively on a single DNA source (i.e., blood or a buccal swab), we sought to evaluate how well single-tissue sequencing captures early developmental sSNVs, which play a significant role in rare disease^54^. We defined early developmental sSNVs as those present in at least two tissues, excluding mutations likely arising from blood-derived clonal expansion or infiltration of blood cells into solid tissues, as mutations arising from these processes do not reflect true early developmental origin (**Figs. S14**, **S15, S16**) (Methods). This yielded 2,718 early developmental sSNVs across 16 donors (∼170 per donor) (**Fig. 5a and Table S6**), including 15 early developmental sSNVs with predicted functional impact (**Fig. 3g**). Consistent with an early embryonic origin, these mutations showed no evidence of age-related increase and were dominated by SBS1 (52%) and SBS5 (42%) (**Fig. S17**), which are ubiquitous across normal tissues.

**Figure 5.**
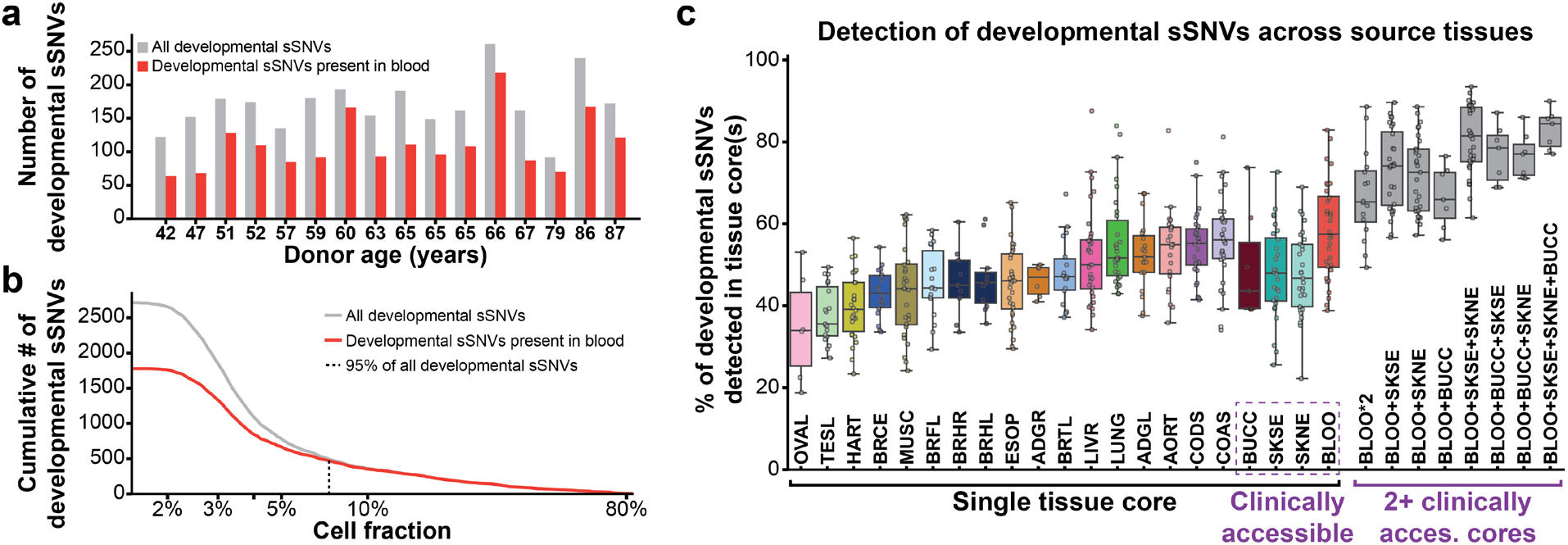
Blood-based sequencing misses one quarter of early developmental sSNVs. **a,** Number of early developmental sSNVs per donor, shown in total (grey) and as detected in either of two blood samples at 150x coverage (red). Donors harboured an average of 170 developmental sSNVs, of which 112 were detected in at least one of two blood samples per donor. **b,** Cumulative count of total and blood-detected sSNVs by cell fraction (defined as 2x variant allele fraction). Dashed line denotes cell fraction at which 95% of developmental sSNVs were detected in blood. **c,** Fraction of developmental sSNVs present in a single tissue core/sample, grouped by tissue. Tissues sorted by median percentage from OVAL to COAS. The detection rate was increased by sampling two tissues compared to one (***p < 0.01): The rightmost 12 columns show the fraction of mutations detected in buccal swab (BUCC), sun-exposed skin (SKSE), non-sun-exposed skin (SKNE), blood (BLOO), blood with double coverage (BLOO*2), blood combined with one (BLOO+SKSE/SKNE/BUCC), two (BLOO+SKSE+SKNE; BLOO+SKSE+BUCC, BLOO+SKNE+BUCC) or three clinically accessible tissues (BLOO+SKSE+SKNE+BUCC).

Notably, on average 33% (14%-55%) of each donor’s early developmental sSNVs, including seven sSNVs with predicted functional impact, were undetectable in blood at current sequencing depth (**Fig. 5a**), underscoring that blood alone is an incomplete proxy even for developmental sSNVs. As expected, sSNVs with higher tissue-wide VAFs, likely reflecting an origin during one of the earliest cell divisions, were more frequently detected in blood: 95% of developmental sSNVs with a VAF greater than 3.6% were also present in blood, compared with a substantially lower detection rate among lower-VAF variants (**Fig. 5b**). Finally, we examined the fraction of early developmental sSNVs detected by sampling blood compared with two or more tissue cores (**Fig. 5c**). The median fraction of detected sSNVs in a single blood sample was 57.5%, which increased significantly (p<0.01) with the addition of clinically accessible tissues, to 73.4% with addition of a single skin biopsy, or to 66% with the addition of a buccal sample (**Fig. 5c**). Notably, the addition of a skin or buccal sample at ∼150x coverage increased the fraction of detected developmental sSNVs more than the addition of a second blood sample (“BLOO*2”). The incomplete detection of early developmental sSNVs in a single tissue is likely due to unbalanced distribution of early cell divisions to different embryonic tissues and the stochastic nature of tissue sampling. Together, these findings demonstrate that single-tissue sequencing, the current standard in clinical genetic diagnostics, systematically underestimates the true burden of early developmental somatic mosaicism and indicates that sampling even one additional clinically accessible tissue, as well as sequencing at deeper (∼300x) coverage could potentially improve diagnostic yield for rare diseases caused by somatic mosaicism.

### Somatic mosaicism across near-T2T genomes

We next examined somatic mutations within genomic loci absent or incompletely represented by GRCh38, including segmental duplications (SDs) and centromeres, which comprise 12% of the human genome^55^, and are central to many human diseases^56^. These loci harbour increased rates of *de novo* mutations^57,58^ and cancer somatic mutations^59^, but resolving their rate of somatic mutation in tissues has been hindered by short-read mapping artifacts^60^ and misassemblies or complete absence within the GRCh38 reference^55^. Donor specific assemblies (DSAs) have the potential to overcome many of these artifacts by providing an accurate diploid germline reference for read mapping^57,59^. For 11 of the 25 donors, we generated a diploid DSA using PacBio HiFi, UL-ONT, and Hi-C data from donor-derived dermal fibroblast cultures (Methods) (**Fig. 6a and Table S7**)^61^, yielding highly contiguous assemblies (median contig N50 135.1 Mbp, median QV 57.7), on par with current gold-standard assemblies^62^ (**Fig. S18**). We built assembly graphs linking each DSA to GRCh38 and CHM13 to compare loci across haplotypes and reference assemblies, revealing ∼396 Mbp of DSA sequence absent from GRCh38 (371-418 Mbp, total 4,370 Mbp), including ∼41 Mbp of non-repeat sequence (39-43 Mbp, total 451 Mbp) (**Fig. 6b**). In addition, DSAs allowed us to identify regions within GRCh38 misassembled relative to each donor’s genome (**Fig. 6b and Fig. S19**), errors that can generate artifacts mimicking somatic mutations. Together, these findings indicate that DSA-based approaches expand the scope of human somatic variant catalogs over GRCh38-based approaches.

**Figure 6.**
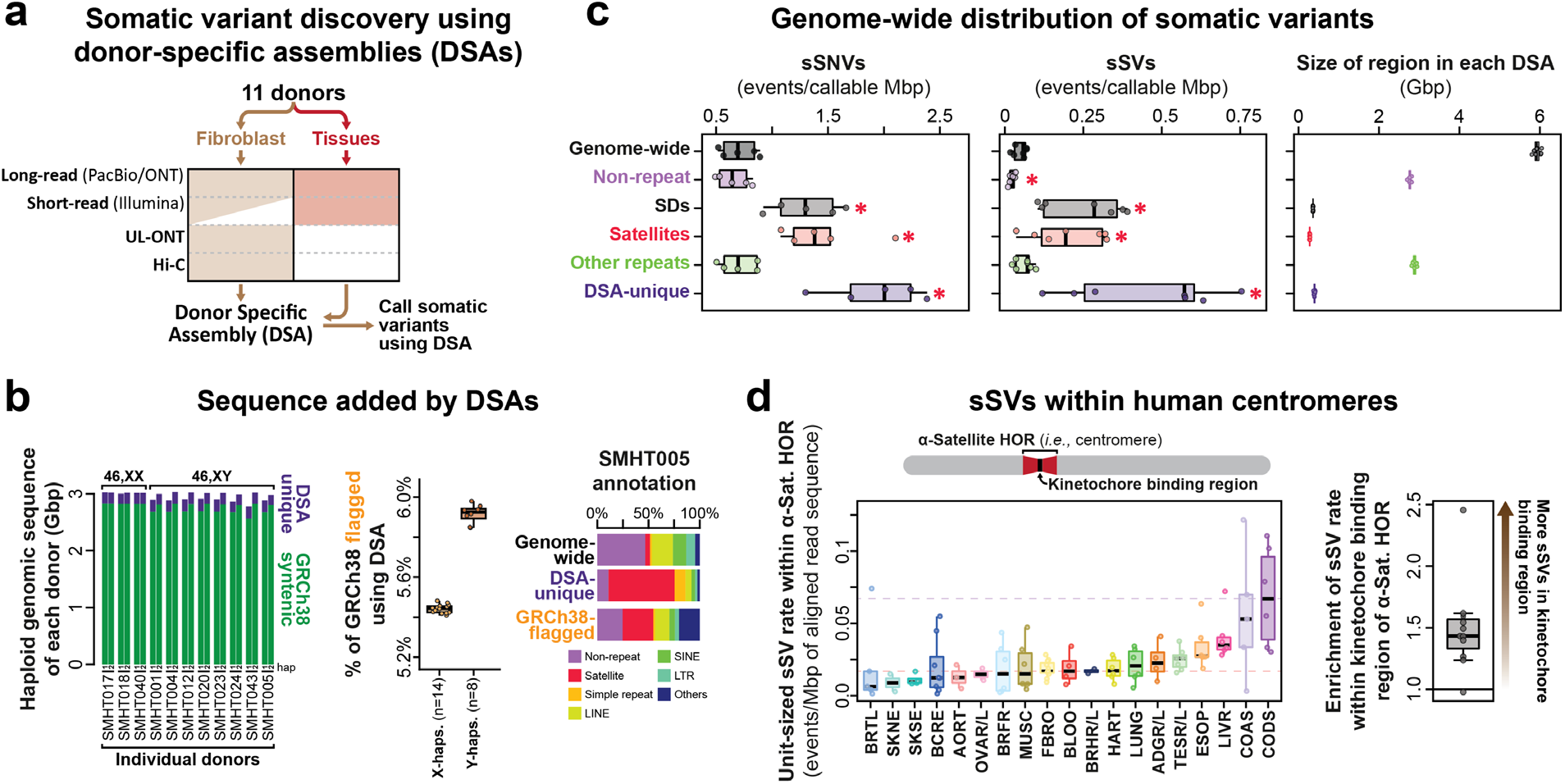
Somatic mosaicism across near-T2T genomes. **a,** Schematic for construction of DSAs for 11 of the 25 donors. The partially filled-in box indicates that only some donors utilized short-read data from fibroblasts. **b,** (Left) DSA-unique sequences, in addition to sequences shared with GRCh38, by haplotype. (Middle) Percentage of GRCh38 flagged as improperly assembled relative to a DSA, split by Y- and X-containing DSA haplotypes. (Right) Annotation of genomic regions in the SMHT005 DSA, including the burden of GRCh38 flagged as inaccurate using the DSA. **c,** Box plots and swarm plots showing the burden of sSNVs (left) and sSVs (middle) across various genomic loci, and the number of gigabase pairs (Gbp) in each genomic bin (right). Each dot represents one donor. *P < 0.05, paired two-sided t test with Benjamini-Hochberg correction. For box plots, the center lines define the median, box limits represent the upper and lower quartiles, and the whiskers extend to themaximum and minimum points within 1.5× the interquartile range; any points beyond are outliers. **d,** (Top) Schematic of a centromere, showing the focal kinetochore-binding regions within the centromeric α-satellite HOR array. (Left) Box plots and swarm plots showing the burden of sSVs across donors, separated by tissue. Dashed lines indicate median burden for colon descending (CODS) and blood (BLOO) samples. (Right) Box plots and swarm plots showing the relative burden of sSVs within the focal kinetochore-binding regions versus the rest of the α-satellite HOR, per donor-tissue pair. Each dot represents a donor.

To characterize tissue somatic variation, we mapped short and long reads from each donor to their DSA and called somatic variants using approaches optimized for sSNV or sSV detection (Methods)^23,59,63^. DSA regions absent from GRCh38 had an elevated burden of sSNVs and sSVs, with SDs and satellite repeats significantly elevated relative to non-repeat genomic loci (**Fig. 6c and Fig. S20, S21**). These results indicate that GRCh38-based catalogs systematically underrepresent true somatic variation. Next, we explored somatic variation within human centromeres, which are composed of α-satellite higher-order repeats (HORs) that markedly diverge in their sequence and structure between individuals^64^, necessitating a DSA-based approach for accurately detecting somatic variation. Across all donors and tissues, sSVs within α-satellite HORs displayed a pattern consistent with arising from break-induced replication (BIR) mediated-repair (**Fig. 6d**), a proposed mechanism for centromeric sSVs^65–68^ that has recently been described in cancers^59,69^. These sSVs were predominantly unit-sized (171 bp multiples), with insertions largely representing perfect duplications of the adjacent HOR array (**Fig. S22**). Furthermore, these sSVs were significantly enriched within the kinetochore-binding domains, which are marked by hypo-CpG methylation^70^ (**Fig. 6d**). Tissues containing dividing epithelial tissues (i.e., esophagus, ascending and descending colon) had significantly higher rates of these sSVs compared to other tissues, consistent with replication-driven sSVs. However, despite its high rate of cell division, blood appeared to have a burden of centromeric sSVs similar to post-mitotic tissues (i.e., heart, brain, etc.), suggesting blood-specific differences in the accumulation or repair of DNA damage within α-satellite HORs. Altogether, these findings expand the landscape of somatic variation across donor tissues and establish a blueprint for T2T studies of tissue mosaicism.

### Stochastic epigenetic states modulate the function of the somatic genome

In addition to somatic genetic variants, mitotically stable stochastic epigenetic states, often termed somatic epimutations, modulate the function of the somatic genome, manifesting as autosomal random monoallelic expression (aRME) or chromatin accessibility (aRMCA). These states can silence or activate regulatory elements, mimic the effects of pathogenic germline or somatic genetic variants, and play a central role in oncogenesis and rare disease^59,71–76^. While stochastic CpG methylation changes are known to increase with age^77,78^, their scope across the human body remains incompletely resolved. Haplotype-specific transcript or epigenetic differences unexplained by imprinting or underlying heterozygous genetic variants can directly pinpoint genes and regulatory elements subject to stochastic epigenetic states (i.e., aRME or aRMCA)^59,79,80^. To determine whether we could observe stochastic epigenetic states within these 25 donors, we first quantified CpG methylation entropy, which captures the degree of disorder in CpG methylation at a given site^81^ and increases with age in humans^82,83^, tracking tissue-specific stem cell division rates^84^. Using 297 long-read ONT datasets across 18 tissues from the 25 donors, we calculated 5mCpG and 5hmCpG methylation entropy genome-wide and at individual cCREs (**Fig. 7a**) (Methods). Donor age was significantly associated with elevated genome-wide entropy (0.00086 units/year; donor-clustered P=1.5×10^-11^; **Fig. 7a**), independent of age-related shifts in cell-type distribution (**Fig. S23**), though the strength of this association varied by tissue (largest SKSE, r=0.76, lowest BRHL, r=−0.51) (**Fig. S24**), potentially reflecting tissue-specific differences in methylation maintenance. These findings validate our ability to quantify stochastic epigenetic processes within these donors and reveal the progressive accumulation of epigenetic features consistent with somatic epimutations over the human lifespan.

**Figure 7.**
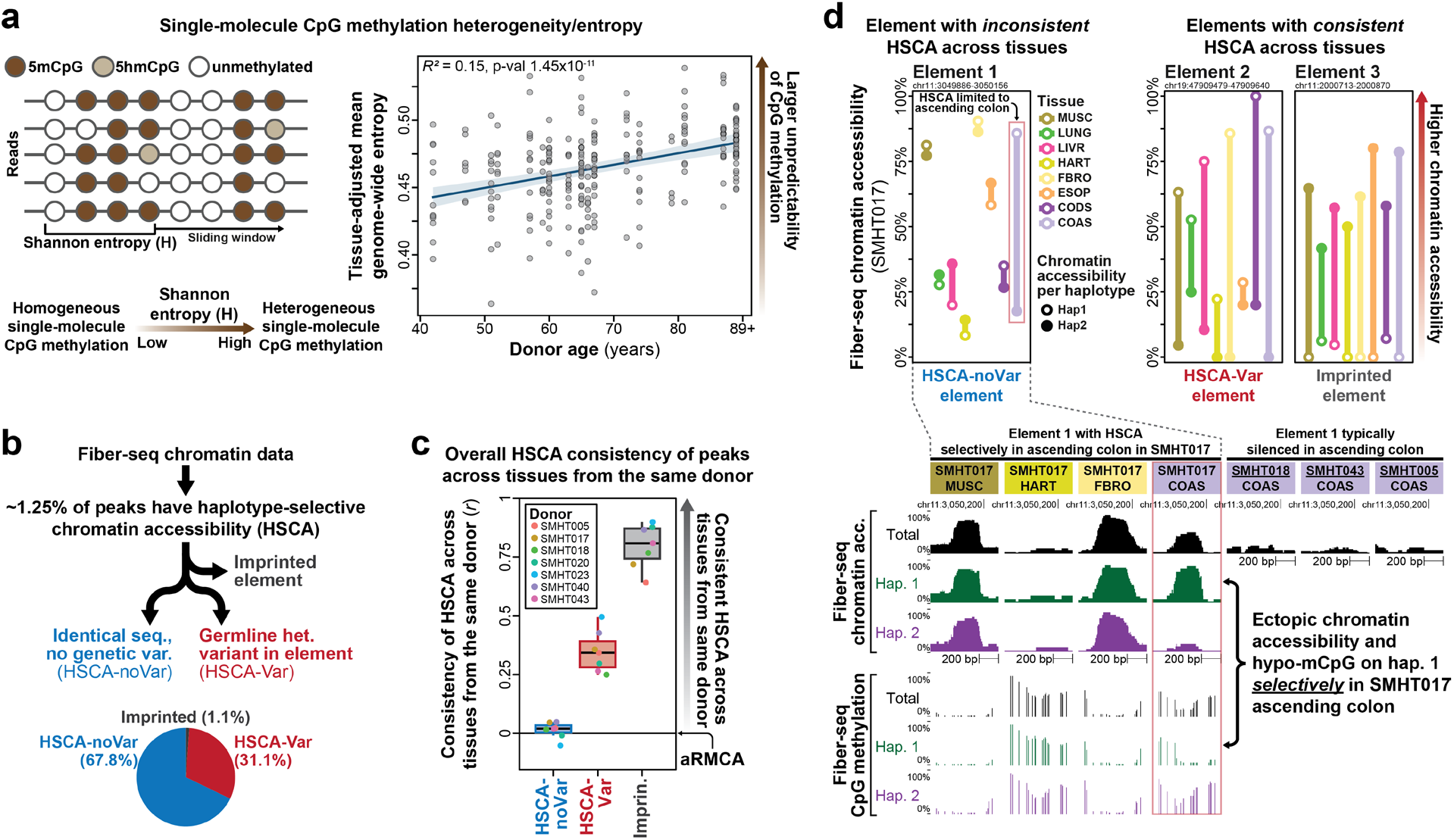
Stochastic epigenetic states modulate the function of the somatic genome. **a,** (Left) Schematic of CpG methylation entropy measurements from single-molecule ONT sequencing data. Rows represent individual reads, with circles denoting 5mC, 5hmC, or unmethylated CpGs; Shannon entropy (H) was calculated from read-level methylation patterns within sliding windows containing four CpGs spanning ≤50 bp, and averaged genome-wide per sample. (Right) Genome-wide methylation entropy within individual tissues from each donor, as a function of donor age. Points show donor-tissue mean entropy values, adjusted to a common, observation-weighted tissue baseline. Line shows the pooled age association estimated by ordinary least-squares regression with tissue-level fixed effects; shaded region shows the pointwise 95% confidence interval, calculated using donor-clustered covariance estimates to account for repeated tissues from the same donor. R2 is reported after tissue-type adjustment. **b,** (top) Flow chart showing the identification of Fiber-seq elements with autosomal haplotype-selective chromatin accessibility (HSCA). (Bottom) Pie chart showing, across all tissues and donors, the overlap of HSCA elements with known imprinted elements or with heterozygous germline genetic variants that may be driving HSCA. **c,** Box plot and swarm plot showing the consistency of HSCA signal at each Fiber-seq element across different tissues from the same donor. Each dot represents one donor. The center line indicates the median and the box encompasses the interquartile range (middle 50% of the data). HSCA-noVar elements show a pattern consistent with autosomal random monoallelic chromatin accessibility (aRMCA). **d,** (Top) Haplotype-resolved Fiber-seq chromatin accessibility of three elements in SMHT017, showing tissue-specific patterns of HSCA. The HSCA-noVar element (left) shows HSCA limited to ascending colon, whereas the HSCA-Var and imprinted elements (right) show consistent HSCA across tissues from the same donor. (Bottom) Fiber-seq chromatin accessibility and CpG methylation tracks for the HSCA-noVar element above, shown across multiple tissues from SMHT017 and in ascending colon from three additional donors. “Total” is the averaged signal across both haplotypes.

We next leveraged 55 high-coverage Fiber-seq datasets across seven donors to localize cCREs with features consistent with aRMCA, using Fiber-seq’s long-read chromatin stenciling to resolve haplotype-selective chromatin accessibility (HSCA) patterns genome-wide^24,79,85^. Overall, ∼1.25% of Fiber-seq elements across all samples showed HSCA (≥25% difference in accessible fibers between haplotypes) (**Fig. 7b**), which we classified into three categories based on each element’s underlying sequence: (1) known imprinted elements (1.1%); (2) HSCA elements harbouring a heterozygous germline variant plausibly driving HSCA (“HSCA-Var,” 31.1%); and (3) HSCA elements with identical sequence between haplotypes and not overlapping a known imprinted element (“HSCA-noVar,” 67.8%). We previously showed that imprinted and HSCA-Var elements show consistent HSCA across tissues, whereas HSCA-noVar elements frequently do not^79^, a defining signature of aRMCA, since genetic or imprinting-driven HSCA should recur across all tissues within a donor, while aRMCA should not. Consistent with this, unlike HSCA-Var and imprinted elements, HSCA-noVar elements showed largely inconsistent HSCA across tissues from the same donor (**Fig. 7c and Fig. S25**). For example, an intronic *CARS1* cCRE typically silenced in ascending colon tissue showed ectopic, haplotype 1-selective chromatin accessibility restricted to ascending colon tissue in SMHT017, directly mirrored by haplotype 1-selective hypo-CpG methylation at the same element (**Fig. 7d**), convergent evidence of aRMCA. Notably, this element, which lacks evidence of imprinting in other tissues from the same donor or other donors (**Fig. 7d**), is located ∼1 Mbp downstream of the *H19*/*IGF2*-imprinting control region (ICR) (**Fig. S26**), but it remains unclear whether HSCA at this element reflects somatic disruption of the *H19*/*IGF2*-ICR imprinting domain selectively within SMHT017’s ascending colon. Together, these findings establish that stochastic epigenetic states manifesting as nongenetically-deterministic chromatin accessibility and CpG methylation changes are pervasive across human tissues.

## Discussion

By integrating multiple classes of genetic and epigenetic variation across up to 20 tissues from 25 postmortem donors, we show that somatic mosaicism cannot be described by a single mutation burden (**Figs. 1b, 2a**): different tissues preferentially accumulated different classes of mutations, and even within the same individual, the tissue with the greatest burden depended on whether sSNVs, sSVs, sMEIs, sCNVs, TRs, or mitochondrial sSNVs were considered. For example, in donor SMHT005 alone, sun-exposed skin dominated for sSNVs, colon for sMEIs, adrenal gland for sCNVs, and testis for mitochondrial sSNVs. The human body therefore represents a mosaic not only of clones but of distinct mutational processes operating at different scales and in different cellular contexts.

Resolving this mutational landscape required technologies with complementary sensitivities^23^. Deep bulk sequencing captures early developmental sSNVs and expanded clones, revealing distinct tissue-specific copy number mechanisms: aneuploidy in adrenal gland^14,29^ versus CN-LOH in testis (**Fig. 1b**). Duplex sequencing extends detection to variants on individual molecules and uncovered tissue-restricted mutational signatures, including a previously undescribed signature in heart, ovary, muscle, and cerebellum (**Figs. 3e, 3f**). Long reads resolve sSVs, TR instability, and sMEIs while retaining haplotype and epigenetic information, revealing striking tissue- and cell-type-specificity, including a concentration of somatic L1 insertions in colon contrasted with Alu-enriched insertions in testis (**Fig. 3d**). Despite the strong cis preference of the L1 retrotransposition machinery^86^, this preferential Alu mobilization in testis may reflect greater Alu transcript abundance, structural features that facilitate access to L1 ORF2p, or stronger suppression of L1 mobilization specifically within testis^87,88^. Single-cell and single-nucleus measurements localize variants to specific developmental lineages and cell types, including tracing an early embryonic mutation directly into the germline (**Fig. 3h**). DSAs resolve variants within loci absent from GRCh38, revealing an elevated burden of sSVs consistent with replication-associated instability within the kinetochore-binding region, establishing them as a focal point of somatic structural instability despite their essential role in chromosome segregation (**Fig. 6d**). No single approach was sufficient on its own, and together, these technologies highlight a continuum from rare events in individual cells to tissue-restricted clones to early developmental mutations distributed across the genome and the body.

Integrating this mutational landscape with RNA, chromatin, and CpG methylation measurements and advanced variant effect predictors further revealed the potential mechanisms driving these mutations, their cellular origins, and their molecular consequences. Using paired RNA expression and CpG methylation data, we identified TRs with cell-type-selective somatic instability, including instances directly associated with tissue-specific expression of the encompassed gene (**Fig. 3b,c**). Using paired CpG methylation and Fiber-seq chromatin data, we identified somatic epimutational processes throughout the body that produce nongenetically-deterministic alterations in genome function (**Fig. 7**). Collectively, these findings demonstrate that a purely DNA-centric view of somatic variation, which is the basis of virtually all existing somatic variant catalogs, is fundamentally incomplete.

This integrated view also impacts how somatic variants should be clinically interpreted. Histologically normal tissues had clones harbouring mutations in canonical cancer-associated genes, including *NOTCH1*, *FAT1*, and *PIK3CA*, consistent with prior reports that positively selected clones driven by these mutations are common in aging skin and esophagus^14,15^ (**Figs. 4d, 4e**). Yet such variants are not synonymous with disease, as their consequences depend on the cell type in which they occur, the size and spatial distribution of the clone, and their functional context. Indeed, our spatial sampling shows that adjacent pieces of the same tissue can contain markedly different somatic landscapes, highlighting that a tissue sampling location alone can strongly determine which mutations are observed (**Figs. 4b, 4c**). This is particularly consequential for the clinical practice of detecting early developmental somatic variants that cause rare disease, which are commonly assessed using a single accessible tissue (blood, buccal swabs, or skin) that may not be phenotypically impacted by the variant in question. Our study showed that deep short-read sequencing of blood at ∼150x coverage detected only 57.5% of an individual’s early developmental somatic variants (**Fig. 5**), consistent with the high false-negative rate observed clinically and raising the prospect that current single-sample clinical workflows underappreciate the contribution of early developmental mosaicism to rare disease. In contrast, sequencing two or more accessible tissues from the same individual increased detection to 66-73% (**Fig. 5**), suggesting that single-tissue testing is an inherently limited diagnostic strategy and that synchronous multi-tissue testing offers a tractable path to improving diagnostic yield. For clinical scenarios in which the prior suspicion for a Mendelian genetic condition is high but routine single-tissue testing is non-diagnostic, we recommend testing an additional clinically accessible tissue rather than doubling coverage of blood DNA (p<0.01, **Fig. 5c**).

Beyond this conceptual gap, our results also demonstrate that current somatic variant catalogs, including the one presented in this study, remain technically incomplete in several respects. Long reads are essential for resolving sSVs, TR instability, and sMEIs (**Figs. 2b, 2c, 3**), and we anticipate that as cost decreases and throughput increases, deeper long-read coverage will further enable detection of variants with low-VAF. Single-cell sequencing plays a central role in identifying late-arising, low-VAF variants restricted to small clones, and we anticipate broader application as the technology matures. Single-cell and spatial profiling will also be essential to resolve which specific cell types mediate the mutational processes described here, as many of the tissues profiled comprise a heterogeneous mixture of cell types, as in adrenal gland where functionally and developmentally distinct cell populations coexist. DSAs revealed that regions poorly captured by GRCh38 harbour some of the highest rates of somatic variation (**Fig. 6b,c**), yet these assemblies still fall short of complete T2T representations, leaving repetitive regions such as ribosomal DNA arrays unresolved. Application of duplex sequencing to post-mitotic, rarely-tumorigenic tissues largely absent from prior somatic variant catalogs (e.g., heart, ovary, muscle, and cerebellum) revealed a previously undescribed mutational signature (**Fig. 3f**), underscoring how much of the somatic variant landscape remains uncharacterized outside of commonly studied tissues. Finally, while paired functional data (RNA expression, chromatin accessibility, CpG methylation) let us directly link somatic variants to their cellular and molecular consequences, such data were not generated for every sample, and often required separate tissue cores. More comprehensive pairing of functional assays with genetic profiling will be needed to fully resolve the functional impact of somatic mosaicism throughout the body. Together, these limitations indicate that the landscape described here should be considered a lower, not upper, bound on the true extent of human somatic mosaicism.

Overall, the initial SMaHT resource of 25 donors establishes a framework for studying somatic mosaicism across human tissues. Expanding to 150 donors, the aim of the SMaHT project, will further enable us to distinguish recurrent, tissue-specific mutational processes from rare events, and determine how mosaicism varies with age, environmental/lifestyle exposures, and germline genetic background. In parallel, the network’s ELSI team is engaging donor families and community stakeholders to assess the risks, challenges, and opportunities of relaying germline and other genomic findings to next of kin, informing future genomic studies involving deceased donors. Continued technology development, more refined donor-specific genome assemblies, and deeper integration of single-cell, spatial, and functional measurements should extend detection to lower-frequency and currently inaccessible variants, and, just as importantly, resolve when each variant arose, where it is distributed, which cells carry it, and whether it alters cellular function. Establishing this baseline across normal human tissues provides the foundation for distinguishing the pervasive mosaicism of development and aging from the subset of events that contribute to disease.

## Data availability

The production datasets described in this study will be made available through the database of Genotypes and Phenotypes (dbGaP) (https://dbgap.ncbi.nlm.nih.gov/) under the study accession number phs004194. Subsets of data and metadata (e.g., somatic variant call sets, transcript levels; age, sex, Hardy scale) are deemed open-access data as per SMaHT Data Use Policy (https://smaht.org/data-use-policy/); raw sequencing data require approval by the investigator’s institutional representative and dbGaP access committee. More information is available at the SMaHT Data Portal at https://data.smaht.org/.

## Acknowledgement

The SMaHT project would not be possible without the generosity of donor families who have provided such precious gifts to support this important work. We are grateful to them and their loved ones’ legacies. We also express gratitude to our community stakeholders for their ongoing input to the study.

## Author List

### OC: Organizational Center - Ting Wang - (U24NS132103)

Ting Wang^1^, Heather A. Lawson^1^, Lucinda Antonacci-Fulton^1^, Casey Andrews^1^, Sarah Cody^1^, Shihua Dong^1^, Milinn Kremitzki^1^, Daofeng Li^1^, Tina A. Lindsay^1^, Shane Liu^1^, Benpeng Miao^1^ 1. Washington University in St. Louis

### TPC: Tissue Procurement Center Thomas Bell - (U24MH133204)

Thomas Bell^1^, Thomas Blanchard^2^, John Clarke^3^, Valerie Estela-Pro^1^, Melissa Faith^4,5^, Abdi Geleta^3^, Michelle Gilbert^6^, Melissa Grimm^7^, Azra Hasan^1^, Emilie Hattrell^1^, Raquel Hernandez^8^, Eric Ho^2^, Robert Johnson^2^, Joseph Kreeb^6^, Alexandra LeFevre^2^, Kathryn Leonard^1^, Phoebe McDermott^8^, Matthew McGillicuddy^7^, Mary Pfeiffer^7^, Russel Roberts^6^, Kristina Sanders^2^, Emmitt Savannah^6^, Isabel Sleeman^1^, Larry Suplee^3^, Nicole Tropello^3^, Patrick Van Hoose^1^, Melissa VonDran^1^

1. National Disease Research Interchange (NDRI)
2. University of Maryland School of Medicine
3. Gift of Life Donor Program
4. Johns Hopkins University School of Medicine
5. Johns Hopkins All Children’s Research Institute
6. LifeGift
7. ConnectLife
8. Johns Hopkins All Children’s Hospital

### DAC: Data Analysis Center (UM1DA058230)

Peter Park^1^, Michail Andreopoulos^1^, Mingyun Bae^1,2^, Michele Berselli^1^, Christophe Boetto^1^, Ann Caplin^1^, Hye-Jung Elizabeth Chun^1^, Shannon Ehmsen^1^, William Feng^1^, Cesar Ferreyra-Mansilla^1^, Rohini Gadde^2,4^, Nils Gehlenborg^1^, Dominik Glodzik^1^, Yujie Guo^1^, Yoo-Jin Ha^1^, Caitlin Harrigan^1^, Neng Huang^1,3^, Hu Jin^1^, Junsoo Kim^1,2^, Jayoung Ku^1,2^, Eunjung Alice Lee^1,2^, Sandra Lee^1^, Heng Li^1,3^, Po-Ru Loh^1,4,5^, Lovelace Luquette^1^, Maximillian Marin^1,3^, Julia Markowski^1^, Dominika Maziec^1^, Utku Öztürk^1^, Han Qu^1^, Suhas Rao^1^, William Ronchetti^1^, Andrew Schroeder^1^, Corinne Sexton^1^, David Tang^1^, Alexander Veit^1^, Vinayak Viswanadham^1^, Seunghyun Wang^1,2^, Kyung Ah Woo^1^, Charlotte Zhang^1^, Yuwei Zhang^1^

1. Harvard Medical School
2. Boston Children’s Hospital
3. Dana-Farber Cancer Institute
4. Broad Institute of MIT and Harvard
5. Brigham and Women’s Hospital

### Genome Characterization Centers (GCCs)

#### GCC-Broad: Broad Institute of MIT and Harvard – Kristin G. Ardlie (UM1DA058235)

Kristin G. Ardlie^1^, Niall Lennon^1^, Pradeep Natarajan^1^, Tim H.H. Coorens^1^, Viktor Adalsteinsson^1^, Lisa Anderson^1^, Mona Arabzadeh^1^, Natalia Brzozowska^1^, Carrie Cibulskis^1^, Fabio Cunial^1^, Laura Domènech^1^, Yushu Huang^1^, Nicola G. Kriefall^1^, Kate Lawrence^2^, Yixuan (Kathy) Liu^1^, Stephen B. Montgomery^2^, Tetsushi Nakao^1^, Azeet Narayan^1^, Stella Ning^1^, Jack Stohlman^1^, Hang Su^1^, Md Mesbah Uddin^1^, Yilin Xie^2^, Zhi Yu^1^

1. Broad Institute of MIT and Harvard
2. Stanford University

### GCC-BCM: Baylor College of Medicine – Richard A. Gibbs (UM1DA058229)

Richard A. Gibbs^1^, Harsha Doddapaneni^1^, Chenghang Zong^1,2^, Elizabeth G. Atkinson^1^, Sravya Venkata Bhamidipati^1^, Jesse Ryan Farek^1^, Marie-Claude Gingras^1^, Christopher M. Grochowski^1^, Walker Hale^1^, Divya Kalra^1^, Ziad Khan^1^, Kavya Chowdary Kottapalli^1^, Heer Hemant Mehta^1^, Donna M. Muzny^1^, Muchun Niu^1,2^, Luis F. Paulin^1^, Jeffrey Rogers^1^, Evette Scott^1^, Fritz J. Sedlazeck^1,2^, Erik Stricker^1^, Kim Walker^1^, Tao P. Wu^1,2^, Ismail Yaman^1^, Yang Zhang^1,2^

1. Human Genome Sequencing Center, Baylor College of Medicine
2. Department of Molecular and Human Genetics, Baylor College of Medicine.

### GCC-NYGC: New York Genome Center – Nicolas Robine (UM1DA058236)

Samuel Aparicio^1^, Timothy R. Chu^1^, Uday Evani^1^, Heather Geiger^1^, Soren Germer^1^, Tausif Hasan^1^, Manisha Kher^1^, Rajeeva Musunuri^1^, Giuseppe Narzisi^1^, Nicolas Robine^1^, Alexi Runnels^1^, Erica Wolin^1^ 1. New York Genome Center

### GCC-UW-SCRI: University of Washington and Seattle Children’s Research Institute – James T. Bennett (UM1DA058220)

James T. Bennett^1,2^, Andrew B. Stergachis^2^, Chia-Lin Wei^2^, Evan E Eichler^2,3^, Natalie YT Au^1^, Marcelo Ayllon^2^, Stephanie Bohaczuk^2^, Yuanye Chi^2^, Colleen P Davis^2^, Danilo Dubocanin^2^, Christian D Frazar^2^, Vea C Freeman^1^, Kendra Hoekzema^2^, Meng-Fan Huang^2^, Caitlin Jacques^2^, Dana Jensen^1^, J T Kolar^2^, Nidhi Koundinya^2^, Youngjun Kwon^2^, Amy Leonardson^1^, Jiadong Lin^2^, Taralynn Mack^2^, Yizi Mao^2^, Sean R. McGee^2^, Anna Minkina^2^, Katherine M. Munson^2^, Shane Neph^2^, Patrick M Nielsen^2^, Michelle D Noyes^1^, Chris Oliveira^2^, Jeffrey Ou^2^, Jane Ranchalis^2^, Luyao Ren^2,3^, Matthew Richardson^2^, Erica Ryke^2^, Adriana E Sedeno-Cortes^2^, Tristan Shaffer^2^, Coey Sit^2^, Joshua D Smith^2^, Min-Hwan Sohn^2^, Melanie Sorensen^2^, Mannat Srivastava^1^, Andrew B. Stergachis^2^, Kaitlyn Sun^2^, Lila Sutherlin^1^, Elliott G Swanson^2^, Mitchell R Vollger^2,4^, Jeffrey Weiss^2^, Isaac Wong^2^, Christina Zakarian^2^

1. Seattle Children’s Research Institute
2. University of Washington
3. Howard Hughes Medical Institute
4. University of Utah

### GCC-WashU-VAI: Washington University in St. Louis and Van Andel Institute – Ting Wang (UM1DA058219)

Ting Wang^1^, Robert Fulton^1^, Hui Shen^2^, Derek Albracht^1^, Eddie Belter^1^, Emma Casey^1^, Paul Cliften^1^, Matthew Cordes^1^, Shihua Dong^1^, Zheng Dong^1^, Qichen Fu^1^, John Garza^1^, Josh Jang^2^, Juan Jiang^1^, Sheng Chih Jin^1^, Nahyun Kong^1^, Zefan Li^1^, Daofeng Li^1^, Tina Lindsay^1^, Shane Liu^1^, Tianjie Liu^1^, Juan Macias^1^, Christopher Markovic^1^, Elvisa Mehinovic^1^, Benpeng Miao^1^, Theron Palmer^2^, Daniel Rohrer^2^, Andrew Ruttenberg^1^, Ayush Semwal^2^, Jiawei Shen^1^, Toni Sinnwell^1^, Zitian Tang^1^, Chad Tomlinson^1^, Yung-Chun Wang^1^, Xiaoyun Xing^1^, Zixi Yu^1^, Wenjin Zhang^1^, Xiaoyu Zhuo^1^

1. Washington University in St. Louis
2. Van Andel Research Institute

### Tool and Technology Development (TTD)

#### TTD-Abyzov: Mayo Clinic – Alexej Abyzov (UG3NS132128)

Alexej Abyzov^1^, Taejeong Bae^1,2^, June Hyug Choi^3^, Yujin A. Choi^3^, Hyungbin Chun^3^, Mrunal Dehankar^1,4^, Yeongjun Jang^1^, Seok-Won Jeoung^3^, Mee Sook Jun^3^, Su Rim Kim^3^, Nam Seop Lim^3^, Nanda Maya Mali^3^, Ji Won Oh^3^, Milovan Suvakov^1^, Yifan Wang^1^

1. Mayo Clinic
2. Korea University
3. Yonsei University College of Medicine
4. University of Minnesota

### TTD-Burns: Dana Farber Cancer Institute – Kathleen H. Burns (UG3NS132127)

Justin Becker^1,2^, Bradley Bernstein^1,2^, Aidan Burn^1^, Kathleen Burns^1,2^, Rony Chanoch^1,2^, Wenchih Cheng^1^, Daniel Fridman^1^, Rohini Gadde^,2,3^, Olympia Hatzilambrou^1^, Katie Irish^3^, Jennifer Karlow^1,2^, Cheuk-Ting Law^1,2^, Eunjung Alice Lee^2,4^, Arnaz Maryam Tariq^1^, Carlos Mendez Dorantes^1^, Adam Voshall^2,4^, Boxun Zhao^2,4^

1. Dana-Farber Cancer Institute
2. Harvard Medical School
3. Broad Institute of MIT and Harvard
4. Boston Children’s Hospital

### TTD-Chen: Broad Institute of MIT and Harvard – Fei Chen (UG3NS132135)

Fei Chen^1^, Jason Buenrostro^1^, Claudia Chu^1^, Tim H. H. Coorens^1,2^, Niklas L. Engel^2^, Gad Getz^1^, Benno Orr^1^, Andrew Russell^1^

1. Broad Institute of MIT and Harvard
2. EMBL-EBI

### TTD-Choudhury: Boston Children’s Hospital – Sangita Choudhury (UG3NS132144)

Zheming An^1,2^, Maniteja Arava^1^, Ming Hui Chen^1,2^, Sangita Choudhury^1,2^, Guanlan Dong^1,2^, Nazia Hilal^1,2^, Taejoo Hwang^1,2^, Se-Young Jo^1,2^, Eunjung Alice Lee^1,2^, Monica Devi Manam^1^, Shulin Mao^1,2^, Diane Shao^1,2^, Christopher Walsh^1,2,3^

1. Boston Children’s Hospital
2. Harvard Medical School
3. Howard Hughes Medical Institute

### TTD-Evrony: New York University – Gilad D. Evrony (UG3NS132024)

Gilad Evrony^1^, Mare Grońska-Pęski^1^, Nisrine Jabara^1^, Jonathan Shoag^2,3^, Amoolya Srinivasa^1^, Jimin Tan^1^, Aristotelis Tsirigos^1^

1. NYU Grossman School of Medicine
2. University Hospitals Cleveland Medical Center
3. Case Western Reserve University School of Medicine

### University of Massachusetts Chan Medical School – Thomas G. Fazzio (UG3NS132136)

Trishita Basak^1^, Thomas G. Fazzio^1^, Manuel Garber^1^, Katrina Newcomer^1^, Sandhiya Ravi^1^ 1. UMass Chan Medical School

### TTD-Landau: Weill Cornell Medicine – Dan A. Landau (UG3NS132139)

Andrew D’Avino^1^, Elliot Eton^1^, Saravan Ganesan^1^, Dan Landau^1^, Yiyun Lin^1^, Levan Mekerishvili^1^, Joe Pelt^1^, Catherine Potenski^1^, Tamara Prieto^1^, Jake Qui^1^, Ivan Raimondi^1^, Rahul Satija^2^, Dennis Yuan^1^

1. Cornell
2. New York Genome Center

### TTD-Marth: University of Utah – Gabor T. Marth (UG3NS132134)

Brad Demarest^1^, Stephanie Gardiner^1^, Stephanie Georges^1^, Gabor Marth^1^, Yi Qiao^1^, Hunter Underhill^1^

1. University of Utah

### TTD-Mills: University of Michigan – Ryan E. Mills (UG3NS132084)

Brandt Bessell^1^, Alan P. Boyle^1^, Ingrid M. Flaspohler^1^, Steve J. Losh^1^, Michael J. McConnell^2^, Ryan E. Mills^1^, Camille Mumm^1^, Jessica A. Switzenberg^1^, Jinhao Wang^1^, Weichen Zhou^1^

1. University of Michigan
2. Rare Mosaic Scientific Consulting, LLC

### TTD-Sedlazeck: Baylor College of Medicine – Fritz J. Sedlazeck (UG3NS132105)

Siyuan Cheng^1^, Adam English^1^, Yilei Fu^1^, Fritz Sedlazeck^1^, Tao Wu^1^, Xinchang Zheng^1^

1. Baylor College of Medicine

### TTD-Urban: Stanford University – Alexander E. Urban (UG3NS132146)

Jan Korbel^1^, Abhiram Natu^2^, Dmitrii Olisov^1^, Reenal Pattni^3^, Pingping Qu^3^, Livia Tomasini^2^, Alexander Urban^3^, Flora Vaccarino^2^, Xiaowei Zhu^4^

1. EMBL-Heidelberg
2. Yale University
3. Stanford University
4. City University of Hong Kong

### TTD-Walsh: Boston Children’s Hospital – Christopher A. Walsh (UG3NS132138)

Irene Antony^1,2^, Alaa Arraf^1,2^, Emre Caglayan^1,2^, Brian Chhouk^1,2,3^, Hayley Cline^1,2^, Liz Enyenihi^1,2^, Benjamin Finander^1,2^, Robert S. Hill^1,2^, Andrea Kriz^1,2^, Alisa Mo^1,2, 4^, Daniel Snellings^1,2^, Ashton Tillett^1,2^, Christopher A. Walsh^1,2,3^, Sijing Zhao^1,2^, Zhou Zinan^1,2^

1. Boston Children’s Hospital
2. Harvard Medical School
3. Howard Hughes Medical Institute
4. Nationwide Children’s Hospital

### TTD-Zong: Baylor College of Medicine – Chenghang Zong (UG3NS132132)

Muchun Niu^1,2^, Yichi Niu^1,2^, Yang Zhang^1,2^, Chenghang Zong^1,2^

1. Human Genome Sequencing Center, Baylor College of Medicine
2. Department of Molecular and Human Genetics, Baylor College of Medicine.

### Associate Members

Rui Chen^1^, Rui Luo^1^

1. UC Irvine

Dmitry Gordenin^1^, Safia Sauty^1^, Yun-Chung Hsiao^1^, Klimczak Leszek^1^

1. National Institute of Environmental Health Sciences

## Contributions

**Writing group leads:** Andrew B. Stergachis, Peter J. Park, Fritz J. Sedlazeck, Tim H.H. Coorens

**Writing group contributors:** Alexej Abyzov, Mona Arabzadeh, Kristin G. Ardlie, Mingyun Bae, James T. Bennett, Rui Chen, Elizabeth Chun, Harsha Doddapaneni, Adam English, Yilei Fu, Richard A. Gibbs, Christopher M. Grochowski, Hu Jin, Sheng Chih Jin, Nicola G. Kriefall, Eunjung Alice Lee, Tina A. Lindsay, Stephen B. Montgomery, Muchun Niu, Jeffrey Ou, Luis F. Paulin, Luo Rui, Corinne Sexton, Min-Hwan Sohn, Flora M. Vaccarino, Vinay Viswanadham, Seunghyun Wang, Weichen Zhou

**Figure 1** is contributed by: Corinne Sexton, Elizabeth Chun, Hu Jin, Nahyun Kong, David Tang, Adam English, Seunghyun Wang, Vinay Viswanadham.

**Figure 2** is contributed by: Corinne Sexton, Hu Jin, Mingyun Bae, Luis F. Paulin, Hang Su, Lovelace J Luquette, Niklas L. Engel, Yeongjun Jang, Weichen Zhou

**Figure 3** is contributed by: Adam English, Luis F. Paulin, Yilei Fu, Seunghyun Wang, Tim H. H. Coorens, Yilin Xie, Luo Rui, Rui Chen, Harsha Doddapaneni, Fritz J. Sedlazeck

**Figure 4** is contributed by: Hu Jin, Corinne Sexton, Michail Andreopoulos, Stephanie Gardiner, Caitlin F. Harrigan, Yilei Fu, David Tang, Elizabeth Chun

**Figure 5** is contributed by: Yeongjun Jang, Yifan Wang

**Figure 6** is contributed by: Min-Hwan Sohn, Tara Mack, Michelle Noyes, Anna Minkina, Juan Macias-Velasco, Issac Wong, Katherine M. Munson, Youngjun Kwon, Danilo Dubocanin, Nidhi Koundinya, Andrew B. Stergachis

**Figure 7** is contributed by: Anna Minkina, Yilei Fu, Mitchell R. Vollger, Shane Neph, Yuanye Chi, Fritz J. Sedlazeck, Andrew B. Stergachis

## Author Contributions

### OC - Ting Wang - (U24NS132103)

T.W. co-led the OC as PI with H.A.L. and L.F. as MPIs. Logistics and Organization were performed by H.A.L., L.A.F., T.A.L., S.C., C.A., and M.K..

### TPC - Thomas Bell (U24MH133204)

T.J.B. supervised the project at TPC as PI. Execution & data collection were performed by T.B., J.C, V.E.P., M.F., A.G., M.G., M.G., A.H., E.H., R.H., E.H., R.J., J.K., A.L., K.L., P.M., M.M., M.P., R.R., K.S., I.S., L.S., N.T., P.V., and M.V.

### DAC - Peter J. Park (UM1DA058230)

P.J.P. supervised the project at the DAC with H-J.E.C., E.A.L., H.L., P-R.L., and N.G. Specific contributions at each group within the DAC are as follows:

#### Harvard Medical School

M.B., S.E., W.C.F., C.F-M., U.O., W.R., A.S., and A.D.V. contributed to data curation and processing under the supervision of H-J.E.C. and P.J.P.

C.E.S. and H.J. performed the somatic variant analysis, with additional contributions from M.A., C.B., A.H.C., D.F., D.G., Y.G., C.F.H., Y-J.J.H., L.J.L., D.M., S.S.L., J.M., H.Q., S.S.R., V.V.V., K.A.W., C.Z., and Y.Z.

H-J.E.C., C.F.H., H.J., L.J.L., C.E.S., V.V.V., and Y.Z. generated figures.

P.J.P., C.E.S., and H.J. wrote the manuscript related to somatic variants, with additional writing contributions from H-J.E.C., C.F.H., L.J.L., and V.V.V.

#### Boston Children’s Hospital

R.G. contributed to data processing under the supervision of E.A.L.

S.W. and M.B. performed the mobile-element-mediated insertion (MEI) analyses, with additional contributions from J.K., J.K., and R.G. under the supervision of E.A.L.

M.B. and S.W. generated figures and wrote the manuscript related to sMEI analyses.

#### Broad Institute

D.T. contributed to copy number variation (CNV) analyses under the supervision of P-R.L.

D.T. generated figures and wrote the manuscript related to sCNV analyses.

#### Dana Farber Cancer Institute

M.G.M. and N.H. contributed to transcriptome analyses under the supervision of H.L.

### GCC-BCM - Richard A. Gibbs (UM1DA058229)

R.A.G. co-led the GCC as PI and EC Co-Chair with H.D. and C.Z. serving as MPIs.

Execution and data collection were performed by S.V.B., M.C.G., K.G., K.C.K., H.H.M., D.M.M., Y.N., M.N., and I.Y.

Analysis and software development were carried out by E.G.A., R.C., A.C.E., J.R.F., C.M.G., W.H., D.K., Z.K., R.L., M.N., L.F.P., J.R., E.S., F.J.S., E.S., K.W., and Y.Z.

### GCC-Broad - Kristin Ardlie (UM1DA058235)

K.G.A. co-led the GCC as PI and EC Co-Chair, with N.L and P.N. serving as MPIs. Execution, data generation, and curation were performed by L.A., C.C., A.N., S.N., J.S., V.A., F.C., and

M.A. Analysis and software development were carried out by L.D., A.N., N.B., H.S., N.G.K., S.N., M.A., K.L., Y.X., T.N., M.M.U, Y.L., Y.H. Data and analysis efforts were supervised by K.G.A., Z.Y., S.B.M., and T.H.H.C.

### GCC-NYGC - Nicolas Robine (UM1DA058236)

N.R. co-led the GCC as PI with S.G. and A.R., with support from S.A.. Data processing and analyses were performed by U.E., H.G., R.M., G.N. and T.C.. Data coordination and sharing were performed by T.H. and M.K. Project management was supervised by A.R., E.W. and N.R.. Experimental method section was contributed by A.R..

### GCC-UW-SCRI - James T. Bennett (UM1DA058220)

J.T.B. co-led the GCC as PI and EC Co-Chair with A.B.S., C.W. and E.E.E. serving as MPIs, with support from J.O., C.D.F., C.P.D., J.M.W., and K.M.M.

Execution and data collection were performed by N.Y.T.A., M.A., J.T.B., C.D.F., V.C.F., K.H., M.H., C.N.J., D.M.J., Y.M., K.M.M., P.M.N., J.O., J.R., E.R., C.S., M.Sorenson., A.B.S., B.C.S., K.S., L.S., and E.G.S.

Analysis and software development were carried out by J.T.B., S.C.B., Y.C., D.D., E.E.E., C.D. F., K.H., J.T.K., N.K., Y.K., A.L., J.L., T.M., S.R.M., A.M., K.M.M., S.N., M.D.N., C.O., L.R.,

M.V., A.E.S., T.S., J.D.S., M.Sohn., M.Sorenson., A.B.S., B.C.S., E.G.S., M.R.V., C.W., J.M.W., I.W., and C.Z.

### GCC-WashU-VAI - Ting Wang (UM1DA058219)

T.W. co-led the GCC as PI with R.F., H.S. serving as MPIs.

Execution and data collection were performed by T.W., R.F., H.S., T.A.L., and T.P.

Analysis and software development were carried out by S.C.J., N.K., Z.T., A.R., J.F.M., D.L., B.M., E.B., C.T., J.E.G., W.Z., Z.X., Q.F., E.M., H.P., X.Z., Z.L., S.D., E.C., Y.C, Y-C.W., and Z.Y.

### TTD-Abyzov - Alexej Abyzov (UG3NS132128)

Y.J. and Y.W. contributed to the analysis of developmental SNVs under the supervision of T.H.H.C, J.T.B., and A.A. Y.J. contributed to building cell lineage tree. A.A. led single cell analysis efforts via PTA.

### TTD-Urban-Flora M. Vaccarino (UG3NS132146)

Execution, data generation, and curation for PTA experiment were performed by A.N. F.M.V. supervised project and co-led single cell analysis via PTA.

### TTD-Burns - Kathleen Burns (UG3NS132127)

B.Z. developed eHAT-seq experimental protocols and analysis pipelines, and generated and analyzed eHAT-seq data. R.G. developed eHAT-seq analysis pipelines and analyzed the data.

A.M.T. contributed to eHAT-seq library construction and validation.

A.B. developed ONT-L1-seq experimental protocols and A.V. developed ONT-L1-seq analytical pipelines; additional contributions are from J.A.K, C.-T. L., C.M.-D., W.-C.C., O.H., J.S.B., D.F., and R.C. Research was supervised by B.E.B., K.H.B., and E.A.L.

### TTD-Choudhury - Sangita Choudhury (UG3NS132144)

N.H. performed experiments, data generation, or running calculations with additional help from

M.A. and Z.A. S.Y.J., G.D., S.M., T.H. contributions with development, benchmarking, application of analytical pipelines and data QC. Under the supervision of D.D.S., C.A.W., E.A.L., S.C..

### TTD-Evrony - Gilad Evrony (UG3NS132024)

N.J., A.S., J.T., and M.G.-P. contributed to methods development under the supervision of A.T., J.S., and G.D.E.

### TTD-Fazzio - Thomas Fazzio (UG3NS132136)

Execution & Data Collection: T.B.; Analysis: K.N. and S.R.; Supervision: T.G.F. and M.G.

### TTD-Landau - Dan Landau (UG3NS132139)

I.R., D.Y., J.P., T.P., Y.L., J.Q., A.D., L.M., E.E., S.G. and C.P. contributed to the network’s single-cell group, under the supervision of D.A.L. and advice from R.S.

### TTD-Marth - Gabor Marth (UG3NS132134)

S.G., S.J.G, H.U. and G.M. contributed to the generation of the short-variant calls at the DAC.

### TTD-Mills - Ryan Mills (UG3NS132084)

R.E.M. and A.P.B. supervised the project at the University of Michigan, with W.Z. providing additional project oversight. J.A.W. performed data generation; W.Z., S.J.L, and B.B. performed somatic variant analysis, with additional contributions from C.M., J.W., and I.M.F.

### TTD-Sedlazeck - Fritz Sedlazeck (UG3NS132105)

L.F.P., A.C.E., Y.F., F.J.S. contributed to manuscript writing. A.C.E., L.F.P., Y.F. contributed to SV data generation, analysis, and software development under the supervision of F.J.S. Y.F. contributed to methylation data generation, analysis, and software development under the supervision of F.J.S.

### TTD-Zong - Chenghang Zong (UG3NS132132)

C.Z. supervised the project at BCM-TTD as PI. Execution, data collection, analysis, and software development were carried out by M.N., Y.N., and Y.Z.

## Funding

This research is supported by the NIH Common Fund, through the Office of Strategic Coordination/Office of the NIH Director under awards U24 MH133204, U24 NS132103, UG3 NS132024, UG3 NS132061, UG3 NS132084, UG3 NS132105, UG3 NS132127, UG3 NS132128, UG3 NS132132, UG3 NS132134, UG3 NS132135, UG3 NS132136, UG3 NS132138, UG3 NS132139, UG3 NS132144, UG3 NS132146, UM1 DA058219, UM1 DA058220, UM1 DA058229, UM1 DA058230, UM1 DA058235, and UM1 DA058236.

*GCC-UW-SCRI – James T. Bennett (UM1DA058220):* M.R.V. and S.C.B. were supported by a T32 training grant from the US National Institutes of Health (NIH) (2T32GM007454-46). M.R.V. was also supported by a Pathway to Independence award from the National Institute of General Medical Sciences (R00GM155552). T.M. was supported by a T32 training grant from NIH (T32HG000035). D.D. was supported by a T32 training grant from the NIH (T32GM141828).

A.B.S. holds a Career Award for Medical Scientists from the Burroughs Wellcome Fund and is a Pew Biomedical Scholar. This work was supported, in part, by NIH grants 1DP5OD029630 and 1U01HG013744 to A.B.S; by the US National Human Genome Research Institute (NHGRI) of the NIH under Award Number R01HG002385 to E.E.E; and by the US National Heart, Lung, and Blood Institute for the NIH under Award Number R01HL130996 to J.T.B. E.E.E. is an investigator of the Howard Hughes Medical Institute.

*TTD-Sedlazeck - Fritz Sedlazeck (UG3NS132105):* Y.F. is supported by SMaHT Pilot U24 NS132103-03.

*TDD-Burns - Kathleen Burns (UG3NS132127):* J.A.K. is supported by an American Cancer Society Postdoctoral Fellow Award (PF-22–123–01-DMC). C.-T.L. has been supported by an American Cancer Society Postdoctoral Fellowship Award (PF-23-1149403) and an NIH Pathway to Independence Award (K99GM157510). C.M.-D. is a Fellow of the Jane Coffin Childs Fund for Medical Research and a recipient of the Charles A. King Trust Postdoctoral Research Fellowship Award.

*TTD-Mills - Ryan Mills (UG3NS132084):* B.B. was supported by a T32 NIH training grant (T32HG000040). I.M.F. was supported by a training grant (T32) from the NIH (T32GM141746).

*TTD-Choudhury - Sangita Choudhury (UG3NS132144):* S.C. was supported by Cell Discovery Network, The Manton Foundation, and The Warren Alpert Foundation.

## Conflicts of interest

N.G. (GCC-Broad) is a co-founder and equity owner of Datavisyn. J.T.B. (GCC-UW-SCRI) is a consultant for Mosaica Medicines. N.L. (GCC-Broad) is an advisor to FYR Diagnostics, Everygene Inc., and has received speaking honoraria from Illumina. C.A.W. (TTD-Choudhury) has consulted for Bioskryb Genomics (cash, equity), Mosaica Therapeutics (equity), Maze Therapeutics (equity), CAMP4 Therapeutics (cash), none of which are relevant to this work. S.B.M (GCC-Broad) is an advisor to BridgeBio, MyOme and PhiTech, none of which are relevant to this work. C.A.W. (TTD-Choudhury) and P.J.P. (DAC) have filed preliminary patent applications for methods to identify mosaic mutations, which are not relevant to this work. F.J.S. (BCM-GCC) receives research support from Illumina, PacBio, and ONT. D.K. (GCC-BCM) immediate family member is employed at ONT. R.A.G. (GCC-BCM) and BCM have equity in Codified Genomics, Inc.. B.E.B. (TTD-Burns) has financial interests in Arsenal Biosciences, Sesame Therapeutics, HiFiBio, nChroma Bio, and Cell Signaling Technologies. A.B.S. (GCC-UW-SCRI) has a patent related to the Fiber-seq method for which he has received royalty payments. E.E.E. (GCC-UW-SCRI) is a scientific advisory board (SAB) member of Variant Bio, Inc. K.M.M. (GCC-UW-SCRI) has received travel and lodging expenses from PacBio to speak at the company-sponsored PRISM event in 2026. P.J.P. (DAC) is a member of the SAB for Bioskryb Genomics, Inc. (cash, equity). M.V. and T.B. (TPC) have a patent (PCT/US2026/025955) related to this work. J.W.O. (TTD-Abyzov) is the founder and CEO of Absolute DNA, with no direct relation to this study, and the interests are managed by University-Industry Foundation in Yonsei University Health System in accordance with their conflict-of-interest policies. All other authors declare no conflicts of interest.

## Declaration of generative AI and AI-assisted technologies

Portions of the text were refined using generative AI and AI-assisted technologies to improve clarity and flow of the English. All revised passages were subsequently reviewed to ensure that the original meaning and intent were preserved.

## Main Methods

### Methods Section 1 | Donors and tissue collection

#### Eligibility criteria (age range, health status, exclusion of known malignancy)

Tissues for the SMaHT study were recovered from postmortem donors by the Tissue Procurement Center’s (TPC) network of three organ procurement organizations (OPOs). SMaHT-specific standard operating procedures (SOPs) for donor screening, authorization for tissue donation, biospecimen procurement and preservation were initiated at each partner OPO. Briefly, donors were considered eligible for inclusion if they were over the age of 18 years, had no known genetic disorders, infectious disease, or active cancer, and tissues could be collected within 24 hours of cardiac cessation^1^.

#### Informed consent and IRB approval at all TPCs

Ethical approval for the sample collection and authorization form for SMaHT was obtained by National Disease Research Interchange (NDRI) through the University of Pennsylvania (IRB#5 FWA00004028) under NDRI’s Tissue Procurement Program protocol (#704541). Authorization for tissue donation from deceased donors was provided by the donor next-of-kin at TPC-partnering OPOs.

#### Fresh-frozen biospecimen handling and cryopreservation

Frozen samples of up to 19 distinct tissue sites were recovered from each eligible donor^1^. All tissues, except the brain, were sampled and preserved on site at the OPO. Blood was sampled into 1cc aliquots and frozen on dry ice. Tissues were sampled into 1×1.5×1cm or 1×1.5cmxfull-thickness aliquots, placed into cryosettes, and preserved using a thermal tray on a bed of dry ice. Whole brain was processed by University of Maryland Brain and Tissue Bank (UMBTB). Four regions of brain (cerebellum, frontal lobe, temporal lobe, and hippocampus) were sampled into 1×1.5×1cm aliquots and flash frozen on dry ice. All samples were stored in a −80°C freezer. The TPC distributed 3.0 mm tissue cores from solid tissue aliquots for downstream analysis. Tissue core extraction was performed using the CXT 353 Bench-Top Frozen Sample Aliquotter (Basque Engineering, West Newbury, MA). Guided by a laser positioning system and a predefined digital extraction map, 3.0 mm tissue cores were extracted from the tissue aliquot and distributed using a labelling schema that maintained spatial orientation within the originating tissue aliquot.

### Methods Section 2 | Sample preparation and sequencing

#### Overview

All assays were performed on material distributed by the TPC from the same 25 donors and, wherever possible, from adjacent 3.0 mm cores of the same tissue aliquot, so that data types can be compared directly within a tissue site and within an individual. Extraction, library preparation and sequencing protocols, including instruments, kits and per-center target coverage, are outlined in the Supplementary Methods. All data are available through the SMaHT data portal (data.smaht.org).

#### Nucleic acids and nuclei extractions

DNA and RNA were extracted from blood, tissue, fibroblast, and buccal samples across different GCCs using validated workflows. Nuclei for UL-ONT and Fiber-seq were isolated by tissue homogenization and filtration, followed by enzymatic processing and recovery of HMW DNA. Nuclei for single-nucleus assays were released by mechanical dissociation and enriched on magnetic beads.

#### Short-read whole-genome sequencing (Illumina)

Bulk short-read WGS was generated for every sample (n = 381, median 368×) on Illumina NovaSeq X Plus and NovaSeq 6000 instruments.

Each sample was sequenced independently at two GCCs, which received physically distinct cores and each targeted ∼300×. Benchmarking established that robust detection of variants below 2% VAF requires more than 300×, with diminishing returns beyond, and that pooling replicate ∼300× datasets extends sensitivity below 1% VAF. The two-center design therefore measured center-to-center variation, confirmed the absence of substantial batch effects, and captured genuine core-to-core clonal variation within a tissue.

#### Long-read whole-genome sequencing (PacBio HiFi and ONT)

Long-read WGS comprised ONT (n = 258, median 63×; PromethION, R10.4.1 flow cells) and PacBio HiFi (n = 237, median 41×; Revio), assigned across GCCs by instrument availability at target depths of 12× to 50× (Supplementary Methods). Long reads were used to validate short-read sSNV calls, were essential for sSV and sMEI detection, and provided whole-genome DNA methylation and chromatin data, in the setting of Fiber-seq. Unlike cell lines, many tissues did not yield DNA of sufficient molecular weight despite considerable effort, and platform success was difficult to predict, so long-read coverage varied widely across tissues and donors. UL-ONT (SQK-ULK114) and Hi-C libraries were generated from cultured donor fibroblasts to support DSAs. Fiber-seq libraries, in which accessible chromatin is marked before sequencing using the non-specific adenine methyltransferase Hia5^24^, were prepared from nuclei of a subset of tissues and sequenced on PacBio Revio to profile chromatin accessibility.

#### Duplex-seq

Four duplex methods were used to detect mutations below the limit of bulk WGS: CODEC, NanoSeq, CompDuplex-seq and META-VISTA-seq. Each derives consensus information from both strands of a DNA molecule, greatly reducing sequencing error^43^. Libraries were prepared using method-specific adapter, barcoding, or transposase strategies and sequenced on Illumina platforms.

#### PTA single-cell DNA sequencing

Single cells or nuclei were isolated from tissue cores of donor SMHT005, amplified by primary template-directed amplification (PTA)^89^ and sequenced to 15×-40× on Illumina, yielding 91 cells from seven tissues. Reads were aligned with the Illumina pipeline described below, followed by somatic mutation calling and single-cell quality control (QC) with SCAN2^90^.

#### Bulk and single-nucleus RNA-seq

Bulk total RNA-seq with ribosomal and globin depletion was generated for 190 samples, and PacBio Kinnex full-length isoform sequencing for 115. For 12 of the 25 donors, single-nucleus RNA-seq and ATAC-seq libraries were generated on the 10x Genomics platform.

#### TEnCATS sequencing

Transposable Element nanopore Cas9-Targeted Sequencing (TEnCATS) targeting L1Hs, AluYa5 and AluYb8 was performed on HMW DNA from donor SMHT005 tissue cores as previously described^91^, with three PromethION flow cells sequenced per tissue.

### Methods Section 3 | Genome alignment, reference assemblies, and QC

#### Reads alignment to the GRCh38 genome

All sequencing data were aligned to GRCh38 (GCA_000001405.15) with ALT contigs and the hs38d1 decoy sequence excluded, using the pipelines established for the SMaHT benchmarking study^23^. An ALT/decoy-free build was used throughout because ALT contigs lower mapping quality and reduce variant-calling sensitivity. Illumina WGS reads were cleared of two-color poly-G artifacts with fastp and aligned with BWA-MEM as implemented in Sentieon, then duplicate-marked, indel-realigned, and base-quality-recalibrated. PacBio HiFi WGS and Fiber-seq reads were aligned with pbmm2 and ONT reads with minimap2^92^, retaining the MM and ML methylation tags in both cases. RNA-seq reads were aligned with the Sentieon implementation of STAR^93^ against a GENCODE v47 index^94^ in two-pass mode, following a pipeline adapted from GTEx and TOPMed, and quantified with both RSEM and RNA-SeQC. Kinnex reads were aligned with pbmm2 under the Iso-Seq preset, collapsed into isoform models with IsoSeq collapse and classified against the reference annotation with Pigeon. Tool versions and alignment parameters are given in the Supplementary Methods.

#### DSA Generation

DSAs were generated from fibroblast DNA for eleven donors using 60× PacBio HiFi, 30× Hi-C and 60× UL-ONT ultra-long data, of which 20× came from reads longer than 100 kbp. Phased assemblies were produced with Verkko v2.2.1^61^, then filtered and quality-controlled with NCBI FCS, Merqury, and Compleasm. The resulting assemblies had a median QV of 57.7, a median contig N50 of 135.1 Mbp, and an average of 24.3 T2T contigs and scaffolds.

#### Quality control

Per-sample QC used the metrics and thresholds validated in the benchmarking study^23^. Alignment statistics and coverage were computed with Samtools, Picard, and mosdepth^95^; sample identity was assessed with Somalier^96^, cross-individual contamination with VerifyBamID2^97^, and human and microbial sequence content with Kraken2^98^. Each dataset was assigned PASSED, FLAGGED, or FAILED status; FAILED datasets were not released, and confirmed sample swaps or mislabels were retracted. The full list of QC metrics and thresholds is given in the Supplementary Methods.

### Methods Section 4 | Somatic variant detection

#### Bulk-based sSNVs

In order to detect sSNVs in SMaHT bulk data, we leveraged the unique qualities of this dataset; namely, we incorporated short- and long-read data types and variant callers, validated calls across multiple tissues in a single donor, and included core-specific information (Supplementary Methods: “Types of variant evidence”). Briefly, we called sSNVs using four variant callers: Strelka2^99^, TNHaplotyper2^100^, RUFUS^101^, and longcallD^102^. These four softwares were run on each core’s data separately and also on the pooled cores’ data as well. Calls then underwent extensive filtering, including removal of germline calls (as determined by DNAscope Hybrid, see Supplementary Methods), population database filtering (gnomAD pop AF < 0.001), problematic genome region filtering (centromere, SDs, simple repeats/satellites), and a panel of normals filtering from the Brain Somatic Mosaicism Network based on 1000 Genomes Project (1KGP) data^103^. Beyond these filters, calls were required to have a minimum alternate read support based on the depth at that variant site. This was determined by a Poisson test based on an assumed sequencing error rate of 0.001. Passing variants then underwent a germline binomial test and if long-read data were available, phasing to determine true somatic calls. After filtering, each variant was marked with PASS if it was 1) called by more than one caller, 2) present in both PacBio and Illumina short-read sequencing, or was found in 3) multiple tissues or 4) cores in the same donor. Calls that survive filtering but not PASS are denoted as

LowEvidence in the VCF FILTER field. The full pipeline with caller parameters as well as downstream filters is available at github.com/smaht-dac/calling-pipelines.

#### Duplex-based sSNVs

CompDuplex-seq and NanoSeq libraries were processed with the unified CompDuplex–NanoSeq pipeline (https://github.com/zonglab/CompDuplex), which performs adapter trimming, alignment to GRCh38, consensus molecule generation, duplex validation and variant filtering on base quality, mapping quality, and duplex support. META-VISTA-seq data were processed with META-VISTA-seq pipeline v2.0.0, which assigns and trims Tn5 duplex barcodes, aligns reads to GRCh38 no-alt with bwa-mem and minimap2, and calls sSNVs and indels from barcode-resolved pileups requiring double-stranded support, cross-aligner concordance, and absence in donor-matched bulk WGS, followed by post-hoc filters on Tn5 insertion-boundary proximity, common gnomAD v4.1 sites, and unmerged-alignment genotype. CODEC data were processed from BCL files to variant calls using CODECsuite best-practice workflows (https://github.com/broadinstitute/CODECsuite), implemented as a WDL pipeline in Terra (https://dockstore.org/my-workflows/github.com/broadinstitute/TAG-public/SingleSampleCO-DEC). Full tool versions, thresholds, and filtering criteria are given in the Supplementary Methods.

#### Mitochondrial sSNVs

Mitochondrial variants were called in donors using a combined short- and long-read workflow with cross-evidence validation (Supplementary Methods: “Mitochondrial variant calling”). Briefly, we called variants with four callers: Mutect2 in mitochondrial mode v1.0^104^ and Mutserve v2.0.3^105^ on Illumina CRAMs, and MitoScope v0.3.0^106^ and Himito v1.1.2^107^ on PacBio and ONT BAMs, run per tissue, donor, and technology on merged alignments. Calls were merged within each sample and standardized by renaming contigs to chrM, left-aligning and normalizing indels, atomizing and splitting multiallelic records, retaining only PASS records from the short-read callers, and removing the blacklisted site chrM:3107. Variants supported by ≥2 technologies, or by ≥2 tissues from the same donor together with ≥2 callers, were assigned HighConf; support from tissues alone or callers alone was assigned LowConf, and remaining calls LikelyArtifact. Heteroplasmy frequencies were averaged per technology, and each variant was classified as homoplasmic or heteroplasmic at a 0.95 VAF threshold and as germline or somatic according to whether it was present in all tissues or only a subset within a donor.

#### Somatic mobile element insertions (sMEIs)

sMEIs were detected using long-read whole-genome sequencing (WGS) data generated on Oxford Nanopore Technologies (ONT) and PacBio platforms. All available long-read WGS datasets from 25 SMaHT donor tissue samples were included in the analysis. Candidate sMEIs were identified using two long-read-based MEI detection methods, PALMER (v2.3.3)^108^ and longcallD (v0.0.8)^102^ (Supplementary Methods, “sMEI callset generation”). When both PacBio and ONT datasets were available for a given tissue sample, each dataset was analyzed independently using both methods. Candidate sMEIs were subsequently filtered to remove germline variants and false-positive calls. To remove putative germline variants, we applied two complementary filters: (1) haplotype phasing and (2) polymorphic MEI site filtering. Using our custom haplotype phasing pipeline^32^, candidates classified as germline were excluded (Supplementary Methods, “sMEI haplotype phasing”). We further excluded germline calls located within 100 bp of polymorphic MEI sites reported in previous population-based studies^109–115^. To filter out false positive calls, we applied three additional filterers: (1) mobile element young subfamily, (2) target-primed reverse transcriptase (TPRT) signatures, and (3) primary alignment status. Candidate insertion sequences were first assigned to mobile element subfamilies using BLAST^116^. Only candidates assigned to evolutionarily young, retrotransposition-competent subfamilies (LINE-1: L1Hs and L1PA2; Alu: AluY, AluYa5, and AluYb8; SVA: SVA_E and SVA_F) were retained for downstream analyses. Then, we annotated TPRT signatures using our custom pipeline and only considered the candidates with at least one TPRT signature (Supplementary Methods, “sMEI target primed reverse transcription (TPRT) and structural feature annotation”), target site duplication (TSD; 2–30 bp), or polyA/T tail (≥10 bp)^32^. Finally, candidates supported exclusively by supplementary or secondary alignments were removed. After all filtering steps, the remaining candidates were manually inspected to confirm TPRT signatures and to verify the correct alignment of both 5′ and 3′ flanking sequences to the insertion breakpoints using BLAT^117^.

#### Somatic structural variants (sSVs)

To discover sSVs, aligned BAMs were processed with three sSV discovery tools: DELLY^118^, Sniffles^119^ and Severus^120^. Per-donor VCFs were merged with BCFtools and genotyped with kanpig^121^, discarding sSVs with fewer than two supporting reads across all samples. Redundant representations were collapsed with Truvari^122^, keeping the representative with the highest alternate read support, and the collapsed set was genotyped again with kanpig. Six variant-level filters removed sSVs in low-confidence regions, sSVs with low read support, and sSVs whose representation prevented somatic and germline alleles from being distinguished. Neighboring sSV counts, sSV class, and gene overlaps were annotated^31,123^.

SVs with a donor-level VAF below 0.20 and at least one sample at ≥10× coverage were taken as somatic candidates. Alignments over candidate loci were refined with LongCallD (doi: 10.64898/2026.03.20.713111), rediscovered, and passed back through the merging, genotyping, and annotation procedure. Candidates were then excluded if no tissue reached ≥10× coverage, if present in a 1KGP panel of normals, if the median mapping quality of supporting reads was <60, if read counts showed a cross-platform batch effect, or if they failed a haplotype-phasing test. For the eight donors with a DSA, two further filters removed candidates matching assembly-derived germline SVs.

Tandem repeats allele length were extracted based on the adotto v2.1 TR catalog, following the same strategy for both PacBio and ONT. Per locus, allele sizes from all samples were clustered without supervision, and loci were classified by cluster number and size. Allele sizes were then redistributed to samples for QC against platform bias and single-read clusters. Highly unstable loci were annotated for gene overlap and inspected manually. Full versions, parameters and filter definitions are in the Supplementary Methods.

##### Somatic copy number variants (sCNVs)

We detected sCNVs using the pipeline described in^28^, which carefully models phased allelic imbalance and normalized read depth in bulk short-read sequencing. In this pipeline, phased allelic imbalance is used to robustly detect events, and normalized read depth is used to classify events as losses, gains, or CN-LOH based on decreased, increased, or unchanged coverage, respectively. Detecting sCNVs from allelic imbalance requires long-range phase accuracy (ideally haplotype blocks longer than tens of megabases) for maximal sensitivity^27^. Hence, we generated phased haplotypes using a joint approach that combined statistical phasing^124^ from large reference panels (https://imputation.researchallofus.org/) with read-backed phasing^125^ from long reads, which produced haplotypes accurate to on average >30 Mbp. The remaining phase switches were accommodated using a hidden Markov model, which identified genomic segments exhibiting consistent, allelic imbalance patterns characteristic of sCNVs. Candidate sCNVs were classified as gains, losses, and CN-LOH probabilistically with priors informed by tissue, chromosome, and event span.

### Methods Section 5 | Data integration and downstream analyses

#### Duplex mutational burden and aging-rate estimation

sSNV burden was quantified by duplex sequencing on four error-corrected platforms (CODEC, CompDuplex-seq, NanoSeq, and META-VISTA-seq) and pooled across assays. For each donor-tissue sample with a positive burden estimate and known donor age, burden in whole blood was regressed on age by ordinary least squares and the association assessed by Spearman correlation. Tissue-specific rates were obtained by regressing burden on age separately for each tissue with at least three sampled donor-tissues, collapsing bilateral sites, and reporting as slopes with 95% confidence intervals alongside a pooled cohort-wide model. Ovary was excluded for insufficient sampling. Only slopes are reported, because reliable intercepts require young donors, which this cohort largely lacks (**Fig. 1f**).

#### Cell-lineage reconstruction from PTA data

Single-cell Illumina CRAMs were assessed with SCAN2^90^ on four metrics: median absolute pairwise difference, GC bias, allele balance and depth distribution; cells failing two or more were excluded, retaining 56 of 91. To sample the early embryonic lineage as broadly as possible, lineage reconstruction instead used every cell with coverage at 52 clonal sSNV loci identified from bulk WGS of the corresponding tissues, irrespective of this filter. Reference and mutant read counts at each locus formed a site × cell allele-count matrix, supplied to Sequoia^126^ with its depth, germline, and beta-binomial filters disabled so that all bulk-derived clonal mutations were retained. Fifteen cells with insufficient coverage across these loci were excluded, leaving 76 for tree building (**Fig. 2e**).

#### Functional annotation of somatic mutations (*Fig. 3g*)

Functional effects of sSNVs were predicted with four tools: PromoterAI^127^ for promoter-proximal effects on expression, SpliceAI^128^ for splice-altering effects, APARENT2^129^ for polyadenylation effects, and AlphaGenome^130^ for tissue-specific expression effects. For the first three, pre-computed scores were downloaded from the respective repositories and intersected with our call set. PromoterAI covers ±500 bp of the TSS of all protein-coding transcripts in GENCODE v39, SpliceAI covers sSNVs within genes up to 50 bp from a splice site, and APARENT2 covers ±100 bp of polyA-DB(R. Wang et al. 2018) cleavage sites. AlphaGenome was run from a local installation on the RNA-seq tracks, matching each tissue to the corresponding GTEx tissue and scoring every gene within ±1 Mbp of the variant.

An sSNV was called predicted functional if it met the threshold for at least one tool: |PromoterAI| > 0.2028 or |APARENT2| > 0.5053, each the 95th percentile of all possible sSNVs in the respective window; a SpliceAI delta score of 0.2 or above, the authors’ recommended high-recall threshold; or an AlphaGenome absolute log2 fold change above 0.2, approximately the 95th percentile for common variants in each RNA-seq track.

#### Mutational signature extraction and decomposition, including identification of novel tissue-associated signatures

Prior to signature extraction, trinucleotide mutation counts from the respective duplex sequencing methods (CODEC, CompDuplex-seq, META-VISTA-seq and NanoSeq) were corrected by using the observed heterozygous germline calls per sample to estimate the trinucleotide channel-specific sensitivity, which serves as an amalgamation for trinucleotide context bias, biases in bioinformatics-calling approaches and other platform-specific effects (see Supplementary Methods). Signatures were then extracted using the Hierarchical Dirichlet Process (HDP) method (https://github.com/nicolaroberts/hdp) without priors, with 10 independent chains, a burn-in of 15,000 iterations, and taking samples every 200 iterations, for a total of 200 samples per chain. This resulted in the extraction of 11 raw HDP signatures, which were then deconvolved into COSMIC reference mutational signatures using an expectation maximization framework. The likely reference signatures present (SBS1, SBS4, SBS5, SBS7a-d, SBS9, SBS12, SBS16, SBS18, SBS19, SBS40a, SBS85, SBS92 and SBS101) were then incorporated into a second HDP run as priors. This resulted in the extraction of six non-reference signatures, and ten known signatures (SBS1, SBS4, SBS5, SBS7a-c, SBS18, SBS19 and SBS40a). One non-reference signature was a residual linear combination of SBS1, SBS5 and SBS40a, rather than a likely true novel signature. These signatures are frequently challenging to deconvolve because of their extremely common co-occurrence^6^. This signature, along with its associated exposure, was decomposed into its constituent signatures (SBS1 [0.12], SBS5[0.54], SBS40a[0.34]), with an overall very high reconstruction cosine similarity (0.97). The signature exposure for SBS5 and SBS40 is reported in a merged manner, as there is frequent cross-attribution between these two clock-like signatures^6,131^. Two other non-reference signatures were strongly associated with specific duplex platforms and may likely reflect a technical origin. This leaves three remaining novel signatures (N1, N2 and N3) of a likely biological origin.

#### QC and cell-class annotation for snRNA-seq and snATAC-seq data

Nuclei were filtered on UMI count, gene count, and mitochondrial fraction for snRNA-seq, and on TSS enrichment and unique fragment count for snATAC-seq. Ambient transcripts were subtracted with SoupX, and doublets were removed with DoubletFinder for snRNA-seq and ArchR for snATAC-seq. Cell classes were annotated per tissue. For snRNA-seq we used reference-free marker-based annotation: highly variable genes were scaled and reduced by PCA, batch-integrated across donors with Harmony, and clustered unsupervised at a resolution set by a bootstrap-stability sweep, after which each cluster was scored against a curated marker panel using its ranked differentially expressed genes. For snATAC-seq, labels were transferred from the tissue-matched snRNA-seq reference by unconstrained Seurat integration through ArchR, following iterative LSI on the tile matrix and Harmony batch correction.

#### Blood fraction estimation from ONT methylation

Blood cell infiltration was estimated from ONT-modified base CRAM files by cell-type deconvolution against a DNA methylation atlas^132^. Read-level methylation calls were generated with wgbstools (v0.3.0-7-g6f24bed) bam2pat in nanopore mode against hg38, and cell-type proportions were estimated per sample with UXM deconv (commit 8d0bb45) against the U25 four-marker hg38 atlas. The blood infiltration fraction was taken as the summed contribution of the atlas cell types Blood-B, Blood-Granul, Blood-Mono+Macro, Blood-NK and Blood-T. Commands and repository links are given in the Supplementary Methods.

#### Identification of driver sSNVs

Candidate driver genes comprised the union of known cancer driver genes^51^ and genes recurrently mutated in clonal haematopoiesis^52,53^ (**Fig. 4e**). Within this gene panel, PASS/LowEvidence sSNVs identified from bulk sequencing were filtered with VEP annotations. The per-donor-tissue call sets were classified as putative driver hits if they met any of three criteria: (i) predicted high-impact consequence or a likely-pathogenic AlphaMissense classification^133^; (ii) a SpliceAI splice-altering score ≥0.8^128^; or (iii) a match to a recurrent, statistically significant mutational hotspot residue (cancerhotspots.org, q<0.1)^134^. Variants also detected in a donor’s whole blood were attributed to blood contamination/haematopoietic clones and excluded from all other tissues of that donor.

#### Derivation of early developmental mutations

Developmental sSNVs arise early and are therefore shared across tissues and samples within a donor (Fig. 5). For each donor and each GCC dataset, candidates were defined as sSNVs present in at least two tissues with at least two supporting reads per tissue, restricted to easy genomic regions (∼74% of the genome) and requiring mapping and base qualities of at least 20. Blood and lung were excluded as discovery tissues to separate tissue-shared developmental mutations from blood-derived sSNVs distributed across tissues (Fig. 4f). Final candidates required consensus between the two GCCs and support from at least one long read in any sample from that donor, which provides further evidence of reproducible presence and authenticity.

Nine of the twenty-five donors showed evidence of clonal hematopoiesis and blood infiltration and were excluded, leaving sixteen. Cell fraction of developmental sSNVs was calculated by concatenating all tissues available for a donor. Percentages of developmental sSNVs across tissues and tissue combinations were compared by paired one-sided t test.

### Methods Section 6 | Statistical analysis

#### Figure 1

For Figure 1e and 1f, duplex burden-age associations were assessed by Spearman correlation (blood) and by linear regression of burden on age, fit separately per tissue and pooled across the cohort, with slopes reported with 95% confidence intervals.

#### Figure 2

Not necessary.

#### Figure 3

For Figure 3c, the significance of difference in the relative proportions of L1, Alu, and SVA insertions between following groups was tested by pairwise chi-squared tests; tissue-specific somatic insertions (Total n=679; L1=677, Alu=2, SVA=0), testis-specific insertions (Total n=15; L1=4, Alu=11, SVA=0), and non-reference germline insertions (Total n=31203; L1=4824, Alu=23154, SVA=3225).

For Figure 3d, the L1 insertion lengths were compared between following groups using pairwise two-sided Wilcoxon rank-sum tests; single-tissue (n=684), multi-tissue (n=22), and non-reference germline (n=4824) events.

#### Figure 4

Not necessary.

#### Figure 5

For Figure 5c, a one-sided t test was performed to compare percentages of detected developmental sSNVs by group.

#### Figure 6

For Figure 6c, p-values were calculated using paired two-sided t tests and adjusted for multiple testing using Benjamini-Hochberg. sSNV p-values: non-repeat 0.0041, SDs 0.0033, satellites 0.0041, other repeats 0.92, DSA-unique 0.0033. sSV p-values: non-repeat 0.0047, SDs 0.0047, satellites 0.0091, other repeats 0.015, DSA-unique 0.0047.

#### Figure 7

For Figure 7c (box and whiskers plot), the center line indicates the median and the box encompasses the interquartile range (middle 50% of the data). For Figure 7b, defining a Fiberseq peak at a consensus peak region requires haploid coverage ≥10, with at least four reads containing a FIRE element, and at least 25% chromatin actuation (reads with FIRE elements/total reads). An HSCA peak in a tissue is defined as one where each haplotype has coverage ≥10, with a peak called in at least one haplotype (≥4 FIRE elements, ≥25% chromatin actuation), a chromatin actuation difference of ≥25% between haplotypes, and a Fisher’s exact nominal p-value of ≤0.005 (comparing the raw fraction of reads with FIRE elements in each haplotype).

