## Supplementary Materials for "Integrated map of somatic mosaicism across human tissues in 25 individuals"

### Supplementary Figures

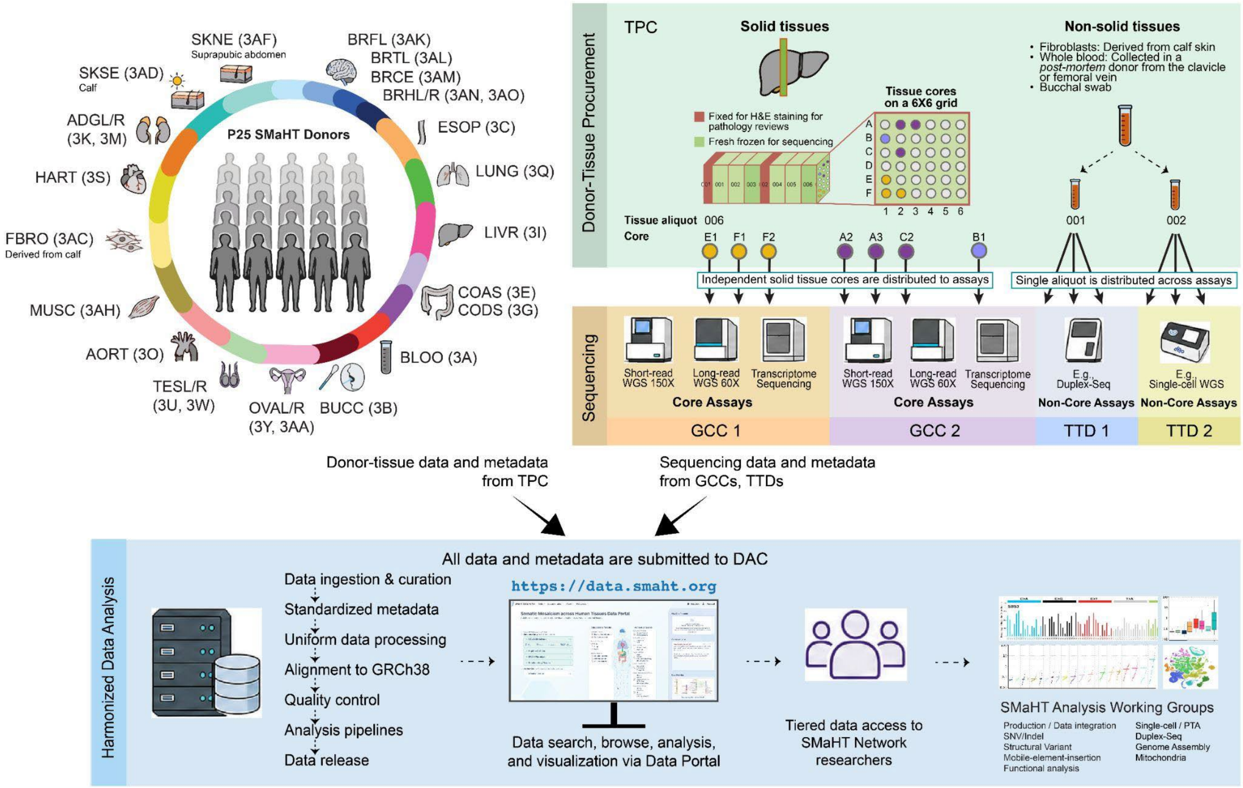


**Figure S1 | Experimental design of the SMaHT production study, related to Figure 1a.** The production phase of the SMaHT Network is carried out as the following: Donor-tissue samples were procured at the Tissue Procurement Center (TPC) and prepared for H&E staining for histopathology review and snap-frozen for sequencing. Independent tissue cores (for solid tissues) or aliquots (for non-solid tissues) were distributed to Genome Characterization Centers (GCCs) and/or Technology & Tools Development groups (TTDs) for sequencing. Metadata and sequencing data are submitted to the Data Analysis Center (DAC), where the data are uniformly processed and analyzed. The data are made available for browsing, searching, visualization, and tier-access downloads via the SMaHT Data Portal (https://data.smaht.org). Multiple analysis working groups were formed to collaborate on various analysis types, including sSNV/sIndel (nuclear and mitochondrial), sSVs, sMEIs, genome assembly, and RNA-seq/functional analyses.

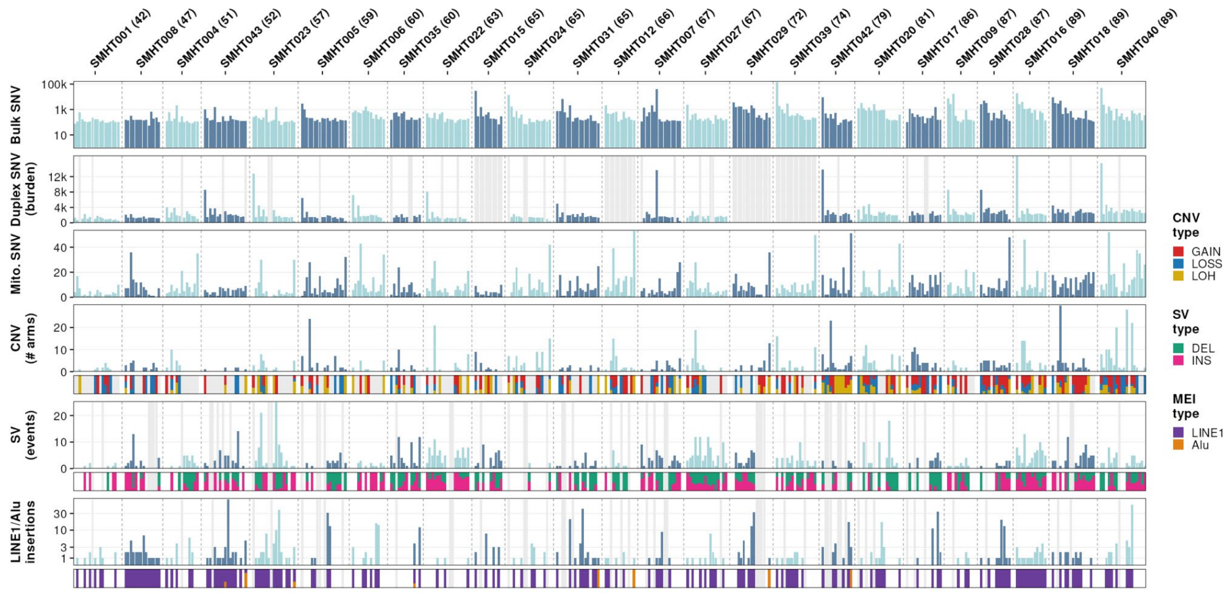


**Figure S2 | Donor-sorted mutational landscape, related to Figure 1b.**
Genome-wide variant burden across six assay classes for 25 donors. Donors are grouped into blocks ordered by ascending age (years); within each block, tissues are ordered by descending median bulk sSNV burden. Bars alternate between two colors by donor block. Top to bottom: Bulk sSNV (PASS calls, log scale); Duplex sSNV burden (CODEC/CompDuplex-seq/NanoSeq-MBN/VISTA-seq); mitochondrial sSNVs; sCNVs (number of chromosome arms affected), with GAIN/LOSS/LOH composition below; sSVs (number of events, VAF ≥5%), with DEL/INS composition below; LINE1/Alu insertions, with element-type composition below. Grey bars indicate data is not available.

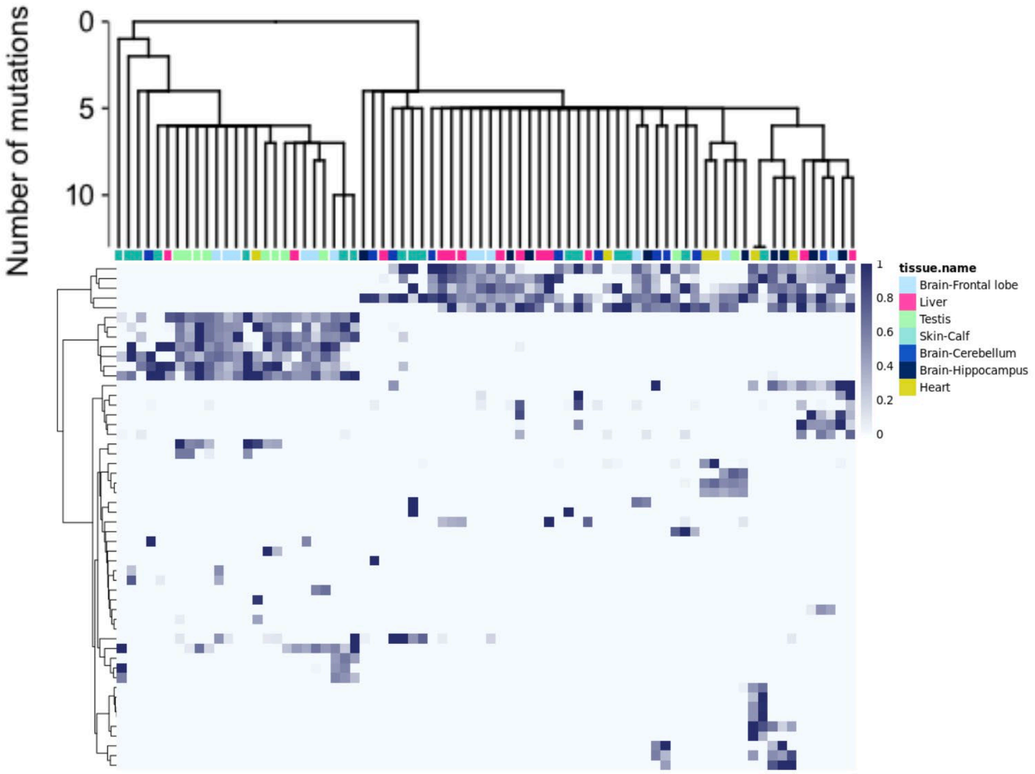


**Figure S3 | Single-cell lineage tree and VAF heatmap for SMHT005, related to Figure 2e.** Cell-lineage reconstruction from 76 PTA-amplified single cells from seven tissues of donor SMHT005 (cerebellum, frontal lobe, hippocampus, heart, liver, calf skin, and left testis). Fifty-two clonal sSNVs identified by bulk WGS of the corresponding tissues were genotyped in the PTA single-cell genomes and used for lineage reconstruction. Tips represent individual cells and are colored by tissue of origin. Terminal branches were truncated at 13 mutations to emphasize the early phylogeny. The accompanying heatmap shows the VAF for each sSNV (rows) across individual single cells (columns), ordered as in the phylogenetic tree.

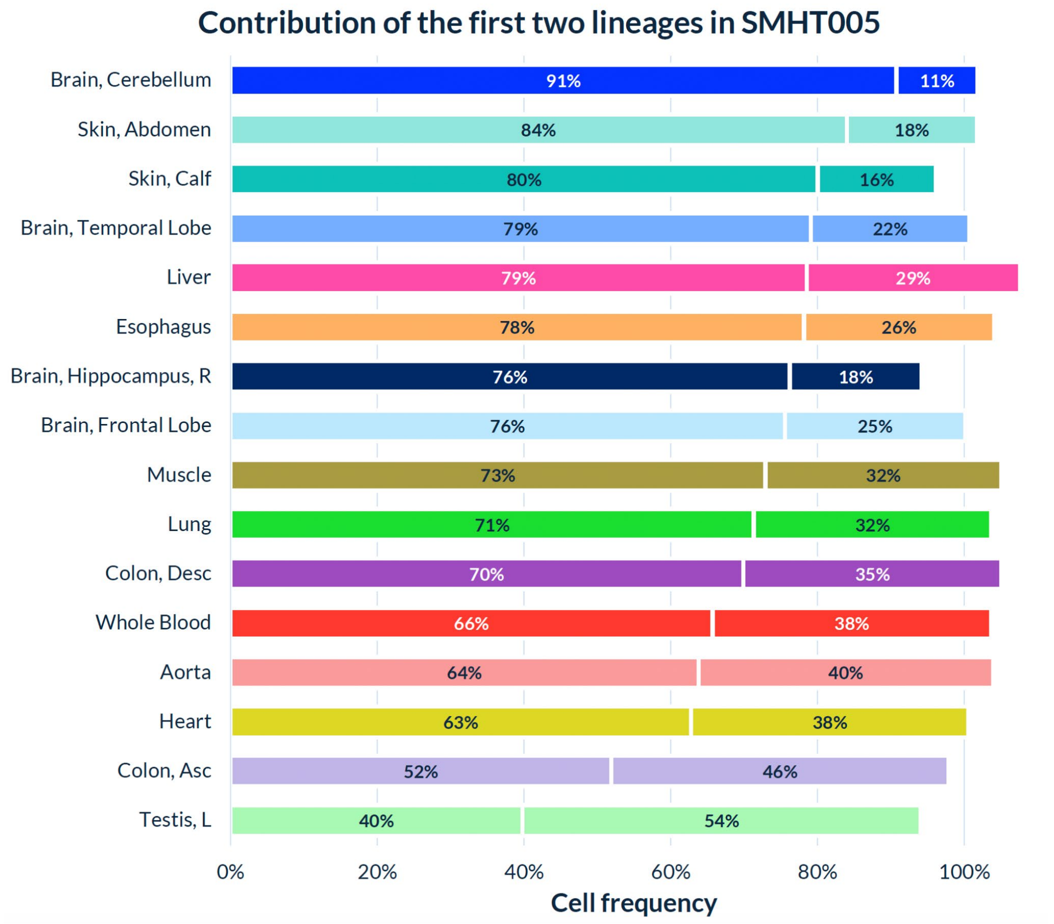


**Figure S4 | Contribution of first two lineages to SMHT005, related to Figure 2e.**
Proportion of the first two embryonic lineages (%) estimated across 16 bulk-sequenced tissues from donor SMHT005. Horizontal bars represent the major (left) and minor (right) lineage fractions per tissue, arranged from bottom to top by ascending dominance of the major lineage. Proportions range from nearly equal contributions in the left testis (54% vs 40%) to marked asymmetry in the cerebellum (91% vs 11%).

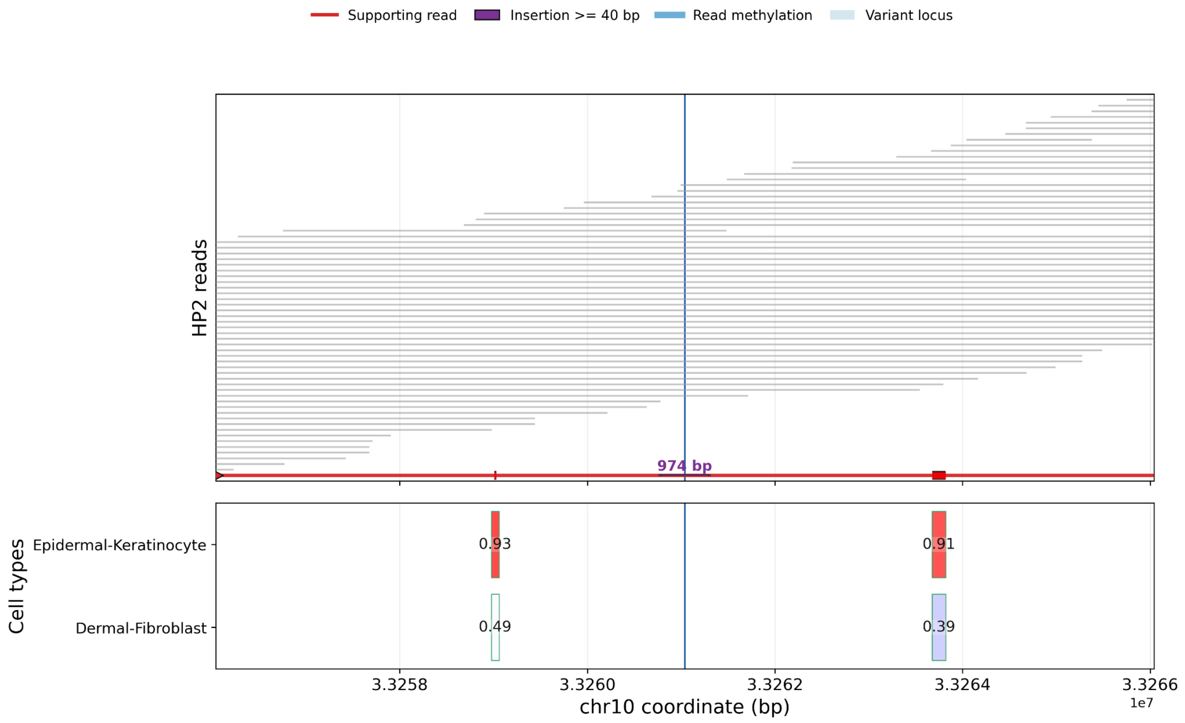


**Figure S5 | Assignment of a sMEI in skin tissue to epidermal keratinocytes, related to Figure 3c.**
SniffCell results showing that a 974 bp insertion in SMHT020 abdominal skin is carried by one ONT read. Its methylation pattern across two ctDMRs within the displayed window supports an epidermal-keratinocyte rather than dermal-fibroblast origin. Gray lines represent other reads from the same haplotype, and the blue vertical line marks the insertion breakpoint.

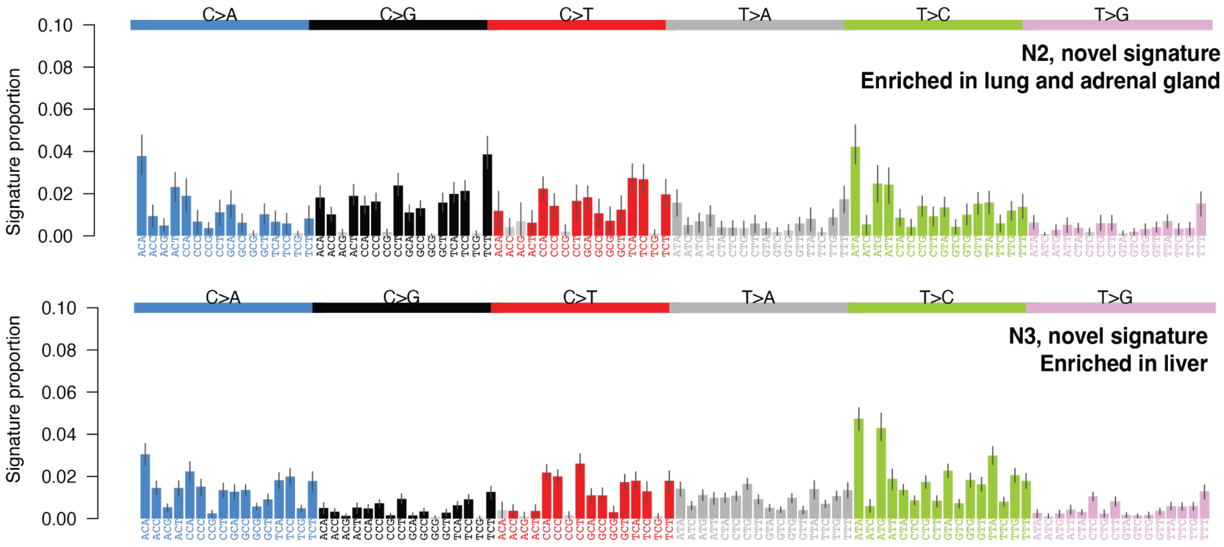


**Figure S6 | Novel signatures from duplex data, related to Figure 3e.**
Spectra of two novel mutational signatures not shown in Fig. 3e, N2 (mainly found in lung and adrenal gland) and N3 (mainly found in liver).

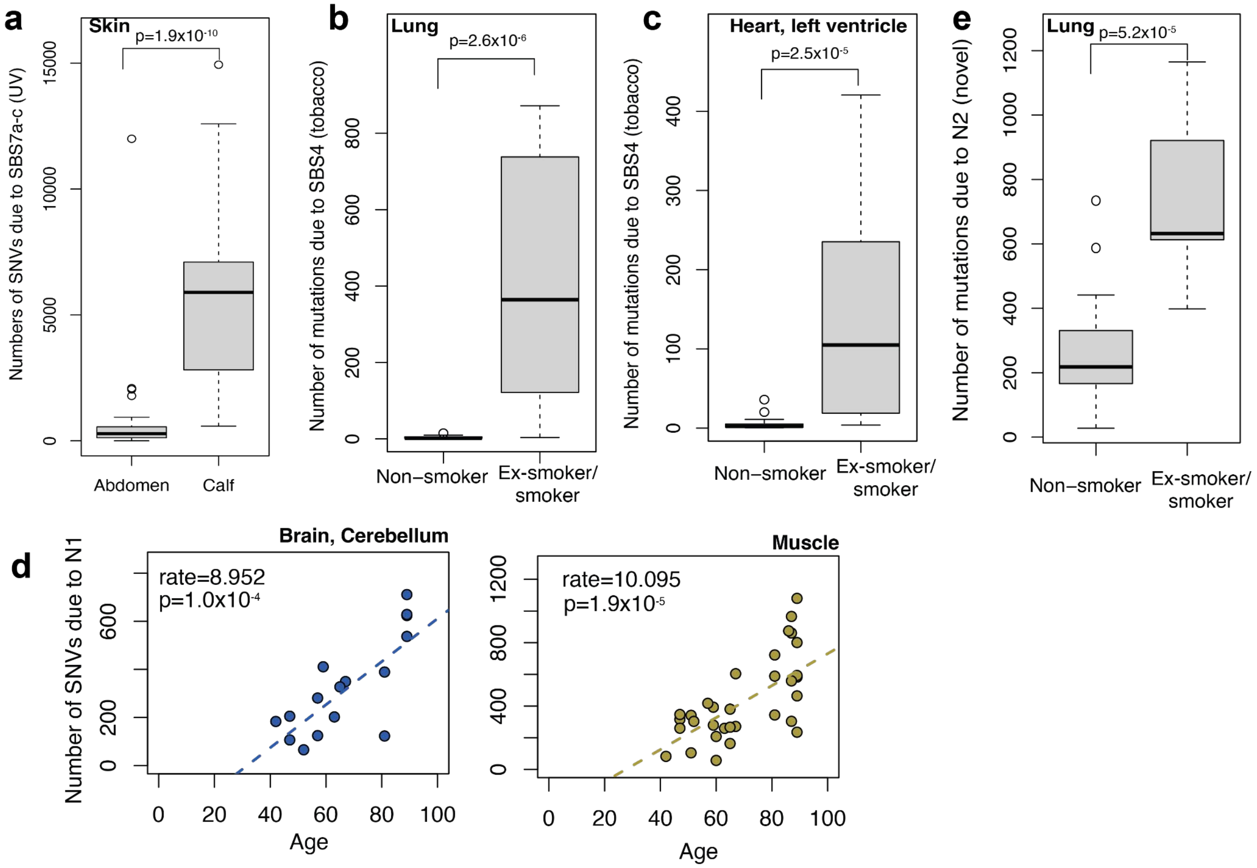


**Figure S7 | Exposure-relation between tissues and age-relation in novel signature, related to Figure 3e.**
Boxplots of sSNV genome-wide mutation burden inferred from duplex data due to SBS7a-d (UV-related mutagenesis) in skin samples from abdomen (17 donors) versus calf (18 donors) **(a)** and tobacco smoke in confirmed (ex-)smokers (9 donors) versus non-smokers (9 donors) in lung **(b)** and heart **(c).** P-values are calculated through a Wilcoxon rank sum test. **d,** Scatterplots of the inferred genome-wide mutation burden due to N1 across age in cerebellum (left) and muscle (right). Dashed lines indicate the slope and intercept as predicted by linear regression, with the sSNV rate per year and p-value (Pearson correlation test) indicated in the inset of the plots. Note that the maximum displayed age is 89. **e,** boxplot of mutation burden due to N2 in non-smokers versus confirmed (ex-)smokers. P-value is calculated through a Wilcoxon rank sum test.

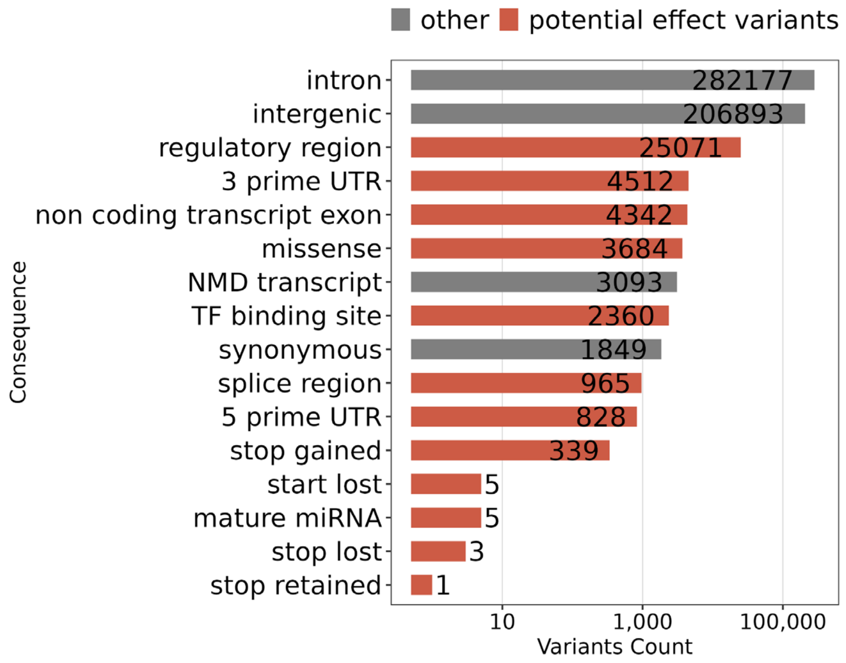


**Figure S8 | VEP consequence distribution among somatic sSNVs, related to Figure 3g.** Bars show sSNV counts for each Ensembl VEP consequence class on a $\log_{10}$-scaled x-axis. Related consequence annotations were collapsed for clarity (e.g., splice-related consequences into splice region; variants in NMD transcripts into NMD transcript). Colors indicate broad impact categories: grey, typically lower-impact/background consequences (intronic, intergenic, synonymous, NMD transcript); coral, consequences more likely to affect gene regulation or coding sequence (regulatory, UTR, missense, splice, stop-gained). sSNVs shared across tissues and/or donors were counted by the number of times they appeared.

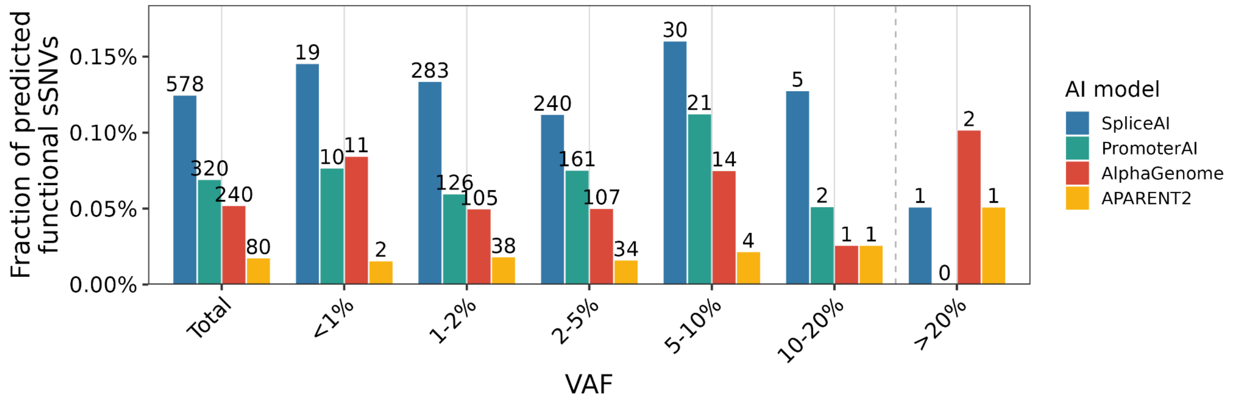


**Figure S9 | Fraction of sSNVs predicted to have a functional effect by each deep learning model across VAF, related to Figure 3g.**
Bars show the fraction of functional variants in that VAF bin (or across all VAFs in Total) predicted by SpliceAI, PromoterAI, AlphaGenome, or APARENT2. For each sSNV, the maximum VAF (from short-read sequencing) across donors/tissues was used to assign a VAF bin. Numbers above bars are absolute counts. Each variant is counted once per model.

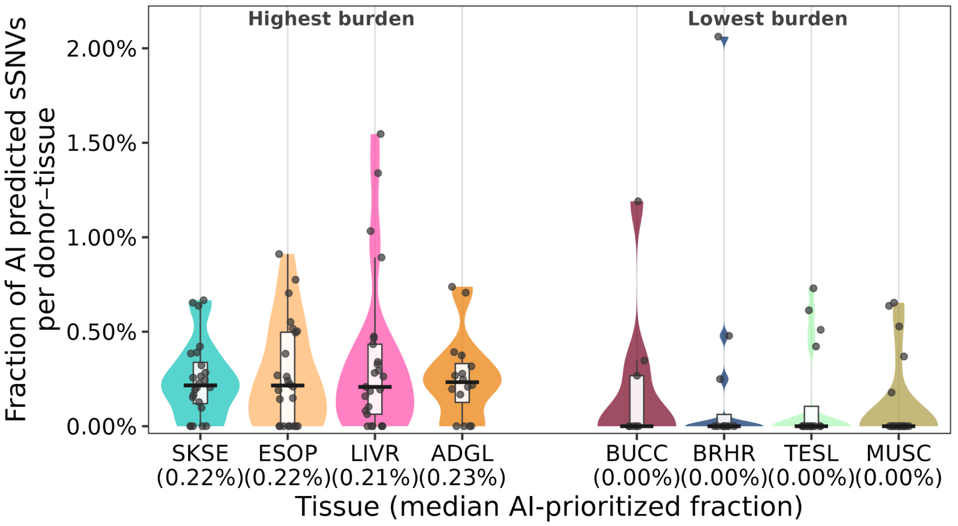


**Figure S10 | Tissue-level fraction of functional sSNVs prioritized by deep learning models in the highest- and lowest-burden tissues shown in Figure 3g, related to Figure 3g.** Violin plots and box plots summarize the across-donor distribution of the fractions of functional sSNVs prioritized by deep learning models per tissue; for each donor-tissue pair, the fraction of unique sSNVs prioritized by ≥1 model is shown. Points are individual donors; the thick horizontal line marks the median (also shown in parentheses on the x-axis).

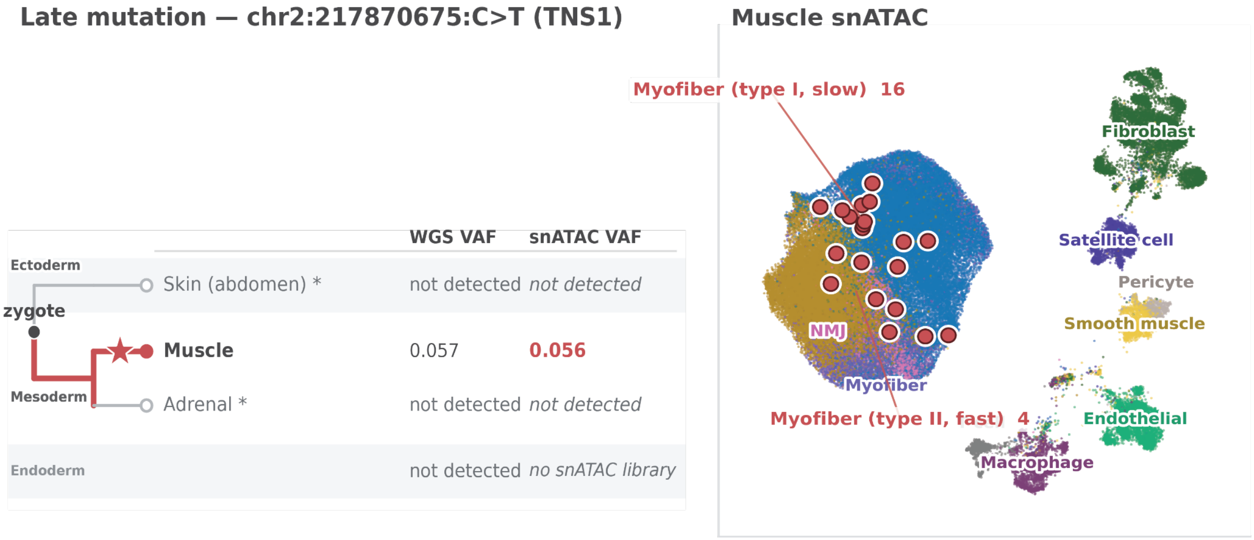


**Figure S11 | Late sSNV detected in snATAC-seq data, related to Figure 3i.**
UMAP of snATAC-seq data from muscle cells from donor SMHT006. Red dots show cells that harbour the identified late sSNV in *TNS1*. This site lies in a called ATAC peak in Muscle alone, so was not detected elsewhere. The muscle UMAP shows all muscle tissue samples from 3 donors with 20 cells carrying the identified sSNV from SMHT006 (Myofiber (type I, slow) 16; Myofiber (type II, fast) 4).

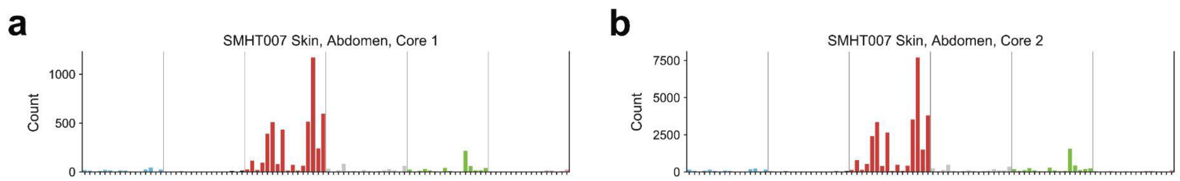


**Figure S12 | Mutational spectra of sSNVs in SMHT007 skin abdomen Core 1 and Core 2, related to Figure 4b.**
sSNV counts across the 96 SBS trinucleotide types in SMHT007 skin abdomen Core 1 (a) and Core 2 (b), with both spectra being dominated by the UV signature SBS7. Note different y-axes between panels a and b.

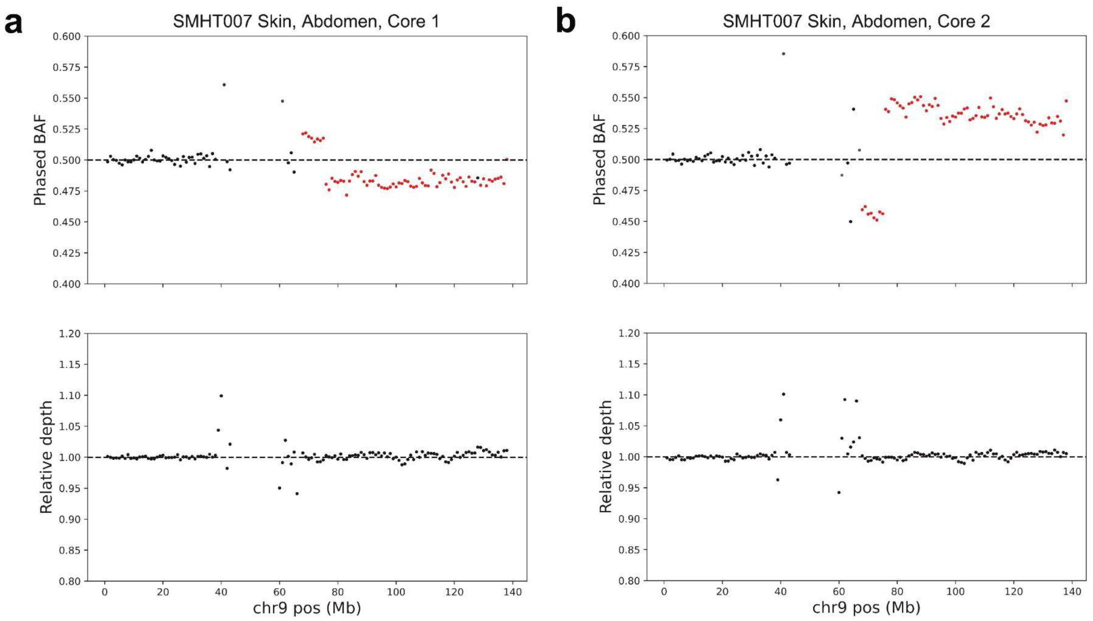


**Figure S13 | 9q CN-LOH in SMHT007 skin abdomen Core 1 and Core 2, related to Figure 4d.**
9q CN-LOH in SMHT007 skin abdomen Core 1 (a) and Core 2(b) showing that the 9q CN-LOH event is on opposite haplotype in each sample. Phased B-allele frequency (BAF, top) and relative depth (bottom) are shown.

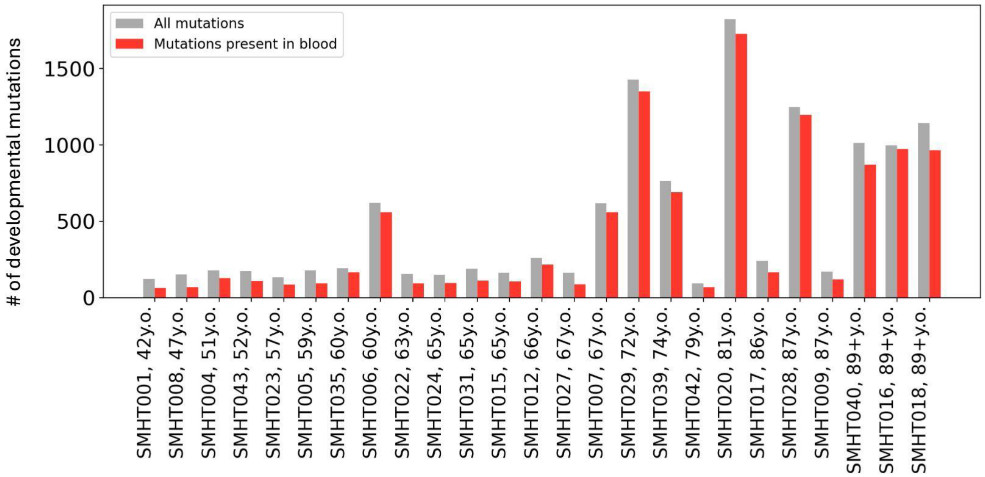


**Figure S14 | Number of early developmental sSNVs per donor, related to Figure 5a.**
Number of early developmental sSNVs per donor for all 25 donors (sorted by age), shown in total (grey) and as detected in either of two blood samples at 150X sequencing depth (red).

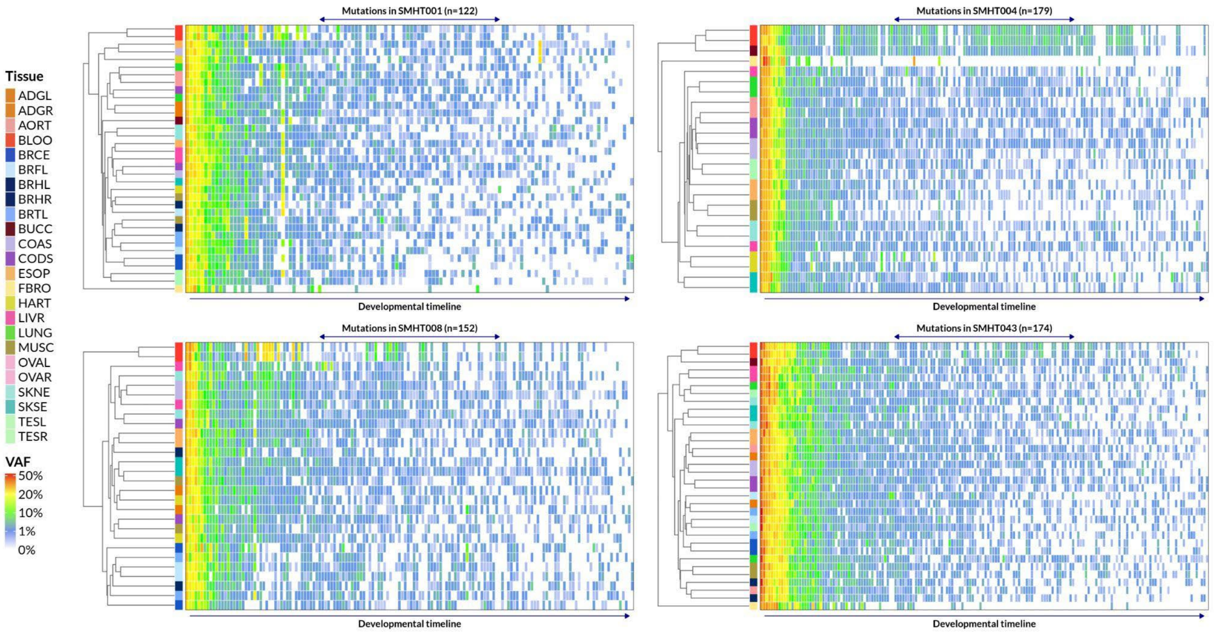


**Figure S15 | Presence of putative early developmental sSNVs across multiple tissues from the same donor, related to Figure 5a.**

Representative examples of mutation presence across tissues in four donors. Every mutation was considered to be present in a sample/tissue if it had at least two supporting reads. Samples/Tissues are clustered by mutations VAF using eJaccard distance and average linkage clustering approach. Mutations are sorted by their average VAF rank in samples/tissues, which is a proxy to developmental order or mutation occurrence. Early mutations are shared across all samples/tissues, while later mutations are shared across a few samples.

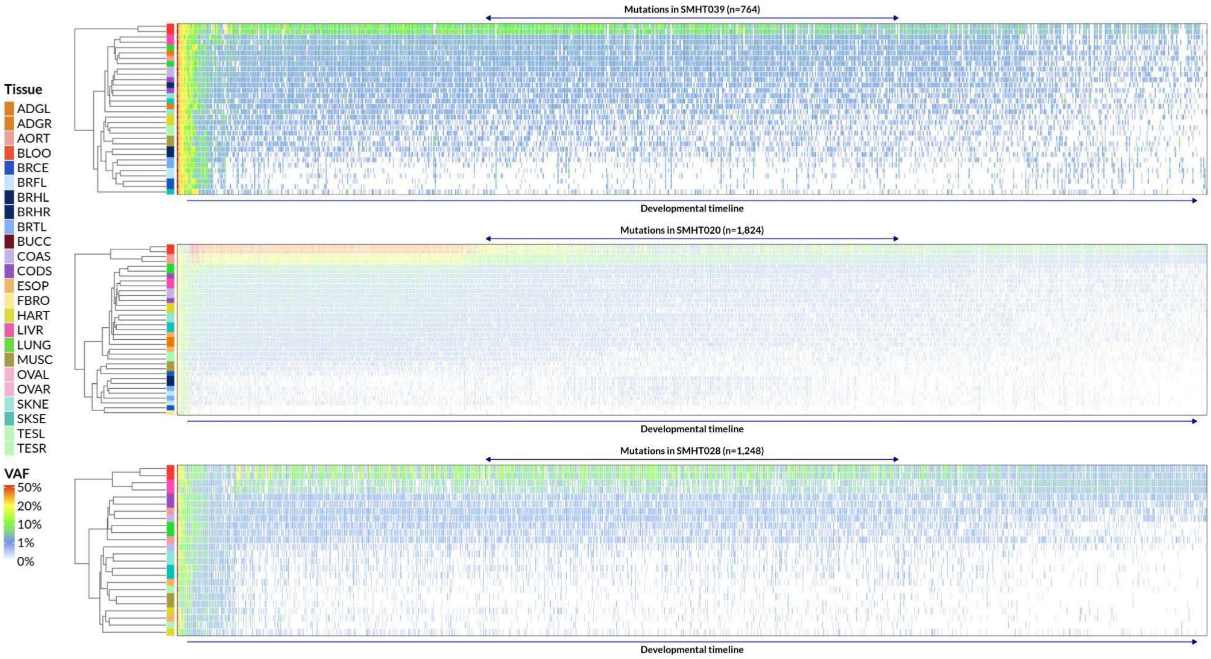


**Figure S16 | Donors with pervasive blood infiltration into solid tissues, as reflected in shared sSNVs having a higher VAF in blood, related to Figure 5a.**

Representative examples of mutation presence across tissues in three donors. Every mutation was considered to be present in a sample/tissue if it had at least two supporting reads. Samples/Tissues are clustered by mutations VAF using eJaccard distance and average linkage clustering approach. Mutations are sorted by their average VAF rank in samples/tissues, which is a proxy to developmental order or mutation occurrence. Most mutations have a hallmark of being originating in blood and infiltrating/contaminating into other tissues. Specifically, often their highest frequency was in blood. Also, brain samples had a minority of the mutations which is likely the results from limited infiltration due to blood-brain barrier.

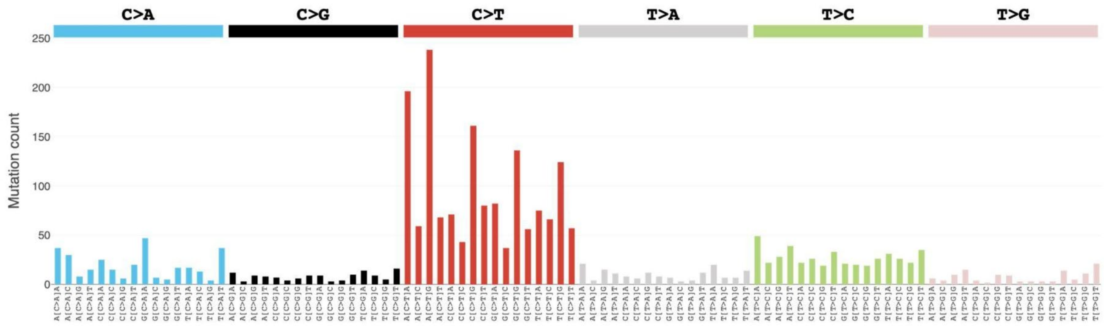


**Figure S17 | Trinucleotide mutational frequency of early developmental sSNVs, related to Figure 5a.**
Trinucleotide mutational frequency of early developmental sSNVs observed across the donors.

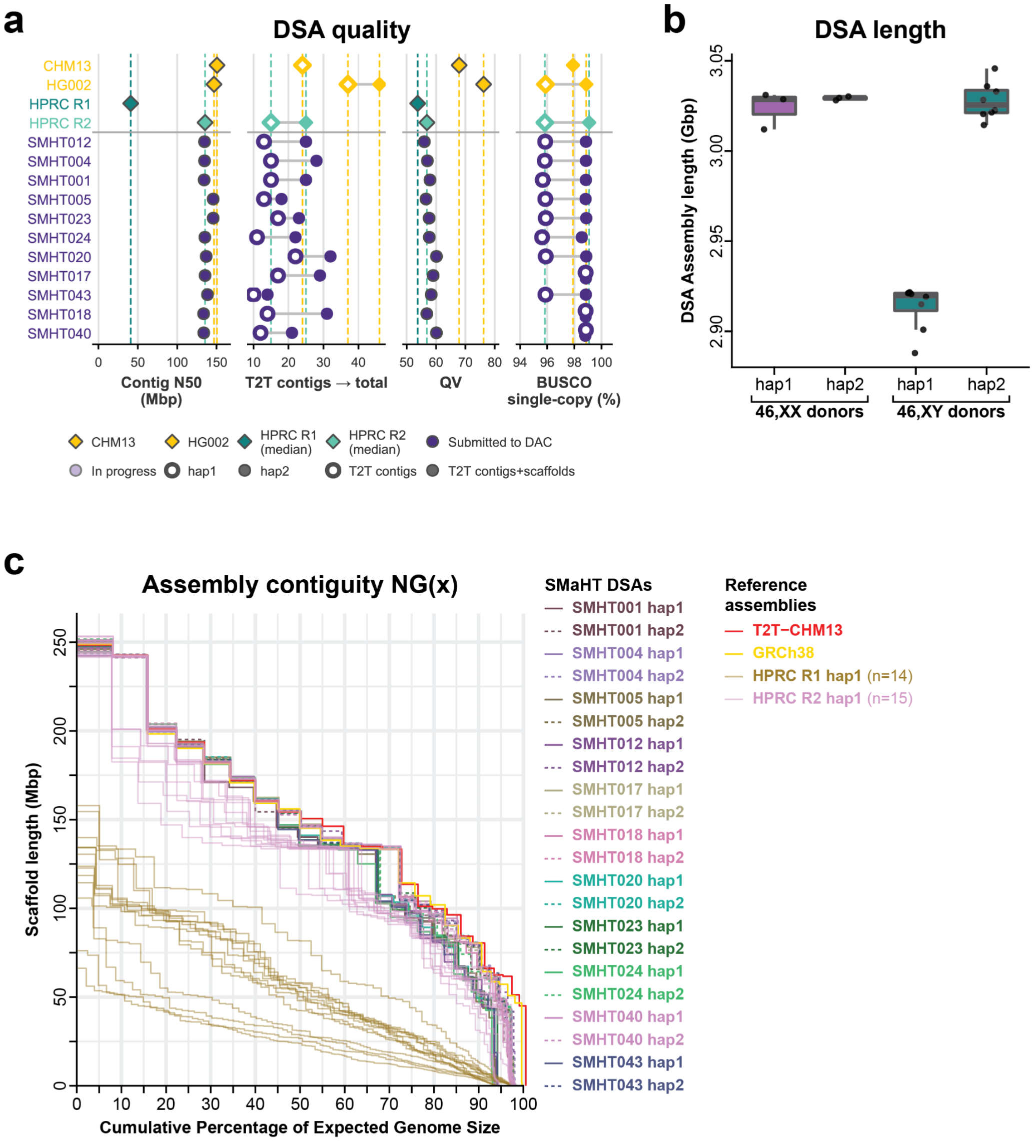


**Figure S18 | Validation of DSA quality, related to Figure 6a.**
**a,** (left) Genome assembly QV (Quality Value) scores (middle) contig N50, and (right) Benchmarking Universal Single-Copy Orthologs (BUSCO) single-copy gene percentage for CHM13, Human Pangenome Reference Consortium (HPRC) release 1 (HPRC R1) and release 2 (HPRC R2), as well as each of the of the DSAs. **b,** Box-and-whisker and swam plot showing the length of haplotype 1 and haplotype 2 assemblies in each of the 11 SMaHT DSAs. Each dot is a different donor. **c,** Assembly contiguity NG(x) plot for each of the SMaHT DSA haplotypes in relation to common reference genomes. NGx corresponds to the contig length such that using equal or longer length contigs produces x% of the length of the reference genome, rather than x% of the assembly length. The expected genome size is set to 3.1Gbp.

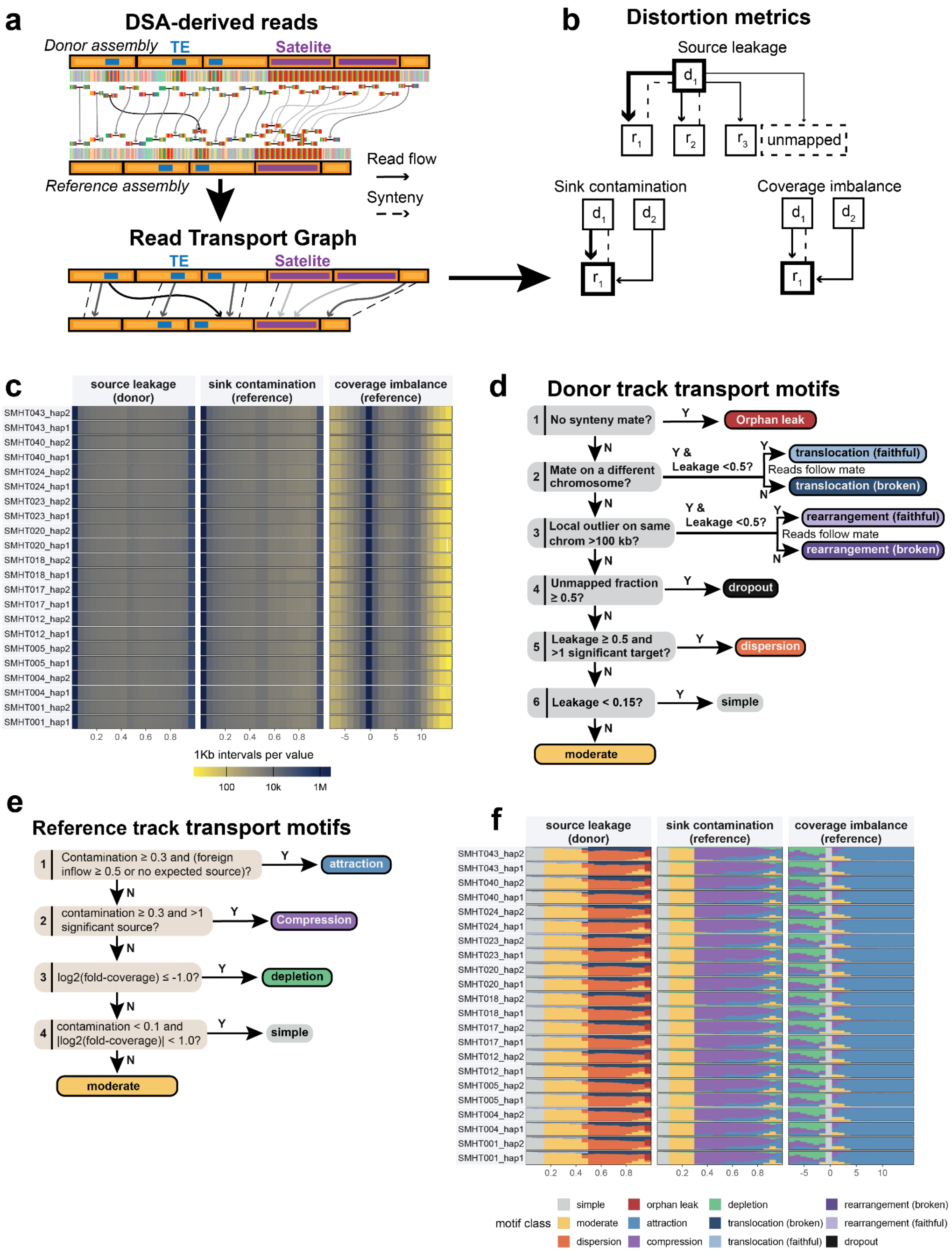


**Figure S19 | Identification of GRCh38 distortions using DSAs, related to Figure 6b.** MoRGANA measures how aligning short reads to a reference genome displaces them relative to their true origin, using each donor's own assembly as ground truth. Reads were simulated from the donor-specific assembly (DSA) of 11 SMaHT donors and aligned both to the reference and back to the donor's own assembly, for both haplotypes of each donor (22 haplotypes total). **a**, Construction of the read transport graph. Reads are simulated from the donor assembly (top track). Each read is then aligned to the reference assembly (bottom track). Solid arrows (read flow) trace where reads land; dashed arrows (synteny) mark where they belong, taken from an the assembly-to-assembly alignment from minimap2. Segmenting donor and reference into 1 kbp intervals yields the read transport graph, whose nodes are donor and reference intervals and whose edges carry the read-flow and synteny relationships between the two coordinate systems. **b**, The three distortion metrics, shown schematically over donor nodes (d) and reference nodes (r). Source leakage is a donor-interval metric, the fraction of an interval's reads that scatter away from its syntenic mate, including reads that fail to map (unmapped sink). Sink contamination and coverage imbalance are reference-interval metrics. Sink contamination is the foreign inflow reaching a reference interval from donor intervals other than its syntenic source(s), and it partitions by origin into DSA-unique (from donor intervals with no syntenic mate), intra-chromosomal, and inter-chromosomal. Coverage imbalance is the log2 ratio of observed to expected read mass at the interval. **c**, Genome-wide distribution of each metric per haplotype. Rows are the 22 haplotypes (donor ID, hap1 and hap2). Color encodes the number of 1 kbp intervals falling in each value bin, on the log density scale at bottom. Source leakage and sink contamination run from 0 to 1; coverage imbalance is log2 observed over expected. **d**, Decision tree assigning each donor interval a transport motif from its read-flow and synteny pattern. An interval with no syntenic mate is an orphan leak. An interval whose mate lies on a different chromosome is a translocation, faithful if leakage is below 0.5 and the reads follow the synteny mate, otherwise broken. A local outlier more than 100 kbp from its synteny mate on the same chromosome is a rearrangement, faithful or broken by the same rule. Of the remaining intervals, one with at least half its reads unmapped is a dropout, one with leakage at least 0.5 across more than one significant target is a dispersion, one with leakage below 0.15 is simple, and the rest are moderate. **e**, Decision tree assigning each reference interval a transport motif. An interval is an attraction if it is dominated by foreign, non-intrachromosomal inflow, taken as contamination at least 0.3 with foreign inflow at least 0.5 or no expected syntenic source. It is a compression if contamination is at least 0.3 across more than one significant source, and a depletion if log2 fold-coverage is at most -1.0. Of the rest, an interval with contamination below 0.1 and absolute log2 fold-coverage below 1.0 is simple, and the remainder are moderate. **f**, Transport motif composition per haplotype. As in (c), over the same 22 haplotypes and metric axes, but each 1 kbp interval is colored by its assigned transport motif rather than by density. The donor source-leakage track uses the panel (d) motifs; the reference tracks (sink contamination, coverage imbalance) use the panel (e) motifs. Colors denote motif class.

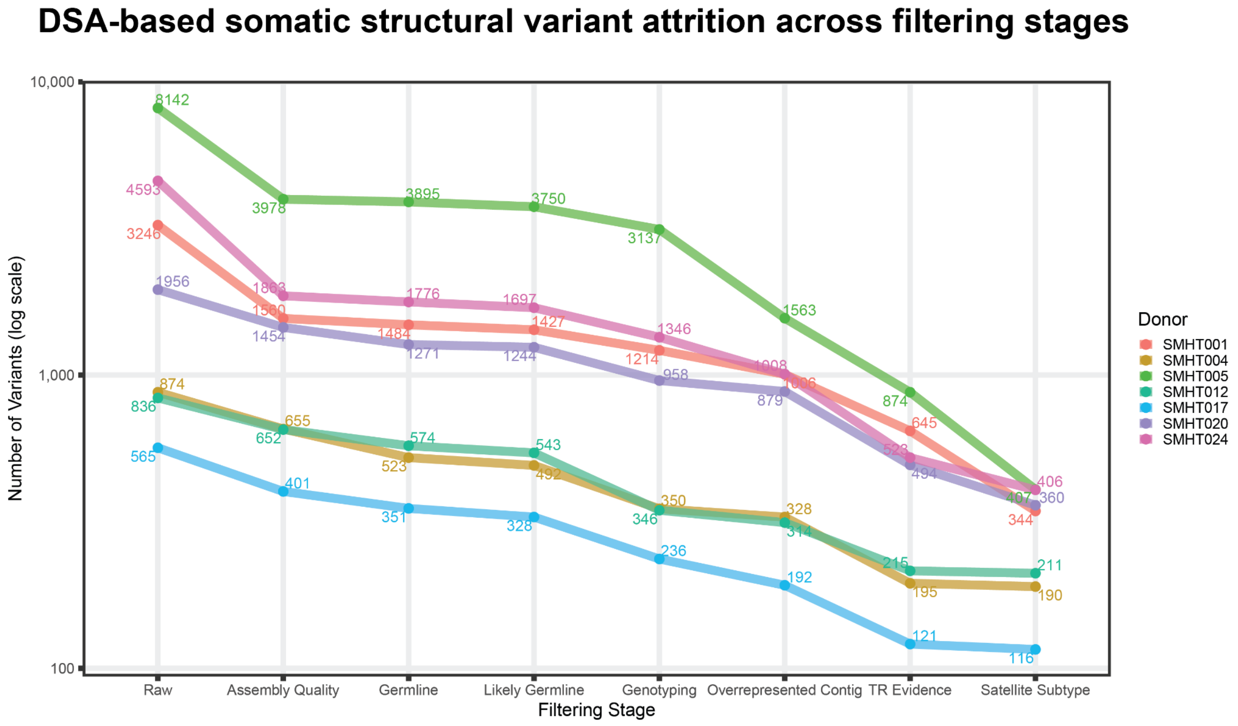


**Figure S20 | DSA-based putative sSV attrition across filtering stages, related to Figure 6c.** Number of multi-caller supported candidate sSVs remaining after each filtering step, per donor (log-scale y-axis), applied cumulatively in the order shown: Raw, putative sSVs supported by ≥2 of 4 callers (DELLY, Severus, Sniffles2, LongcallD); Assembly Quality, >10% overlap with a NucFlag/Flagger-flagged misassembled region; Germline, matches an inherent GRCh38-DSA germline difference; Likely Germline, present in >70% of tissues with hVAF ≥0.9 or 0.4–0.6; Genotyping, lacking kanpig support; Overrepresented Contig, on a contig with an outlier call count; TR Evidence, in a tandem repeat region lacking both Severus and LongcallD support; Satellite Subtype, RepeatMasker SAR or BSR_Beta annotation. Variants drop out at the earliest stage they fail and labeled points show the count remaining per donor at each stage.

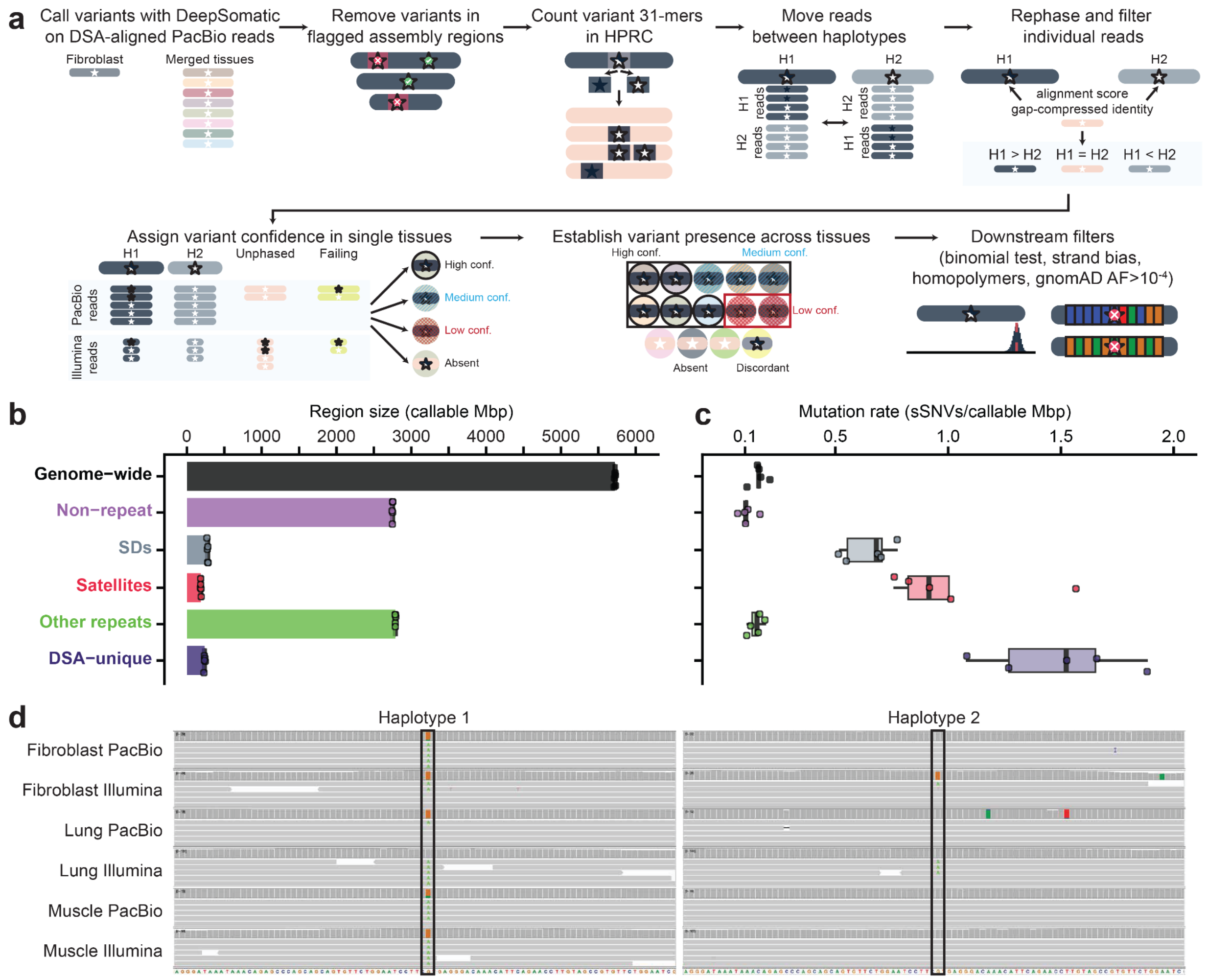


**Figure S21 | DSA-based identification and filtering for sSNVs, related to Figure 6c.**
**a,** Schematic outlining the key steps of the DSA-based sSNV calling and filtering pipeline. **b,** Amount of callable space across genomic regions for n=5 samples, with error bars representing 1 standard deviation. Callable space was defined as all assembled base pairs, excluding regions flagged by NucFlag or Morgana. **c,** Distributions of sSNV rate pooled across tissues for n=5 donors, excluding fibroblast-unique mutations, maintain significant enrichments in segmental duplications (SDs), satellites, and DSA-unique regions, as well as a significant depletion in non-repeat regions relative to the genome-wide rate. The center lines define the median mutation rate for each region, box limits represent the upper and lower quartiles, and the whiskers extend to the maximum and minimum points within 1.5× the interquartile range; any points beyond are outliers. *P-values* were calculated using paired t tests and adjusted for multiple testing using Benjamini-Hochberg: non-repeat 0.0013, SDs 0.0013, satellites 0.0048, other repeats 0.19, DSA-unique 0.0013. Single asterisk: $p < 0.05$. **d,** IGV screenshots of a G>A sSNV located in a satellite region of donor SMHT004. This event was identified in fibroblast, lung, and muscle tissues, and is shown in read data aligned to both haplotypes of the DSA.

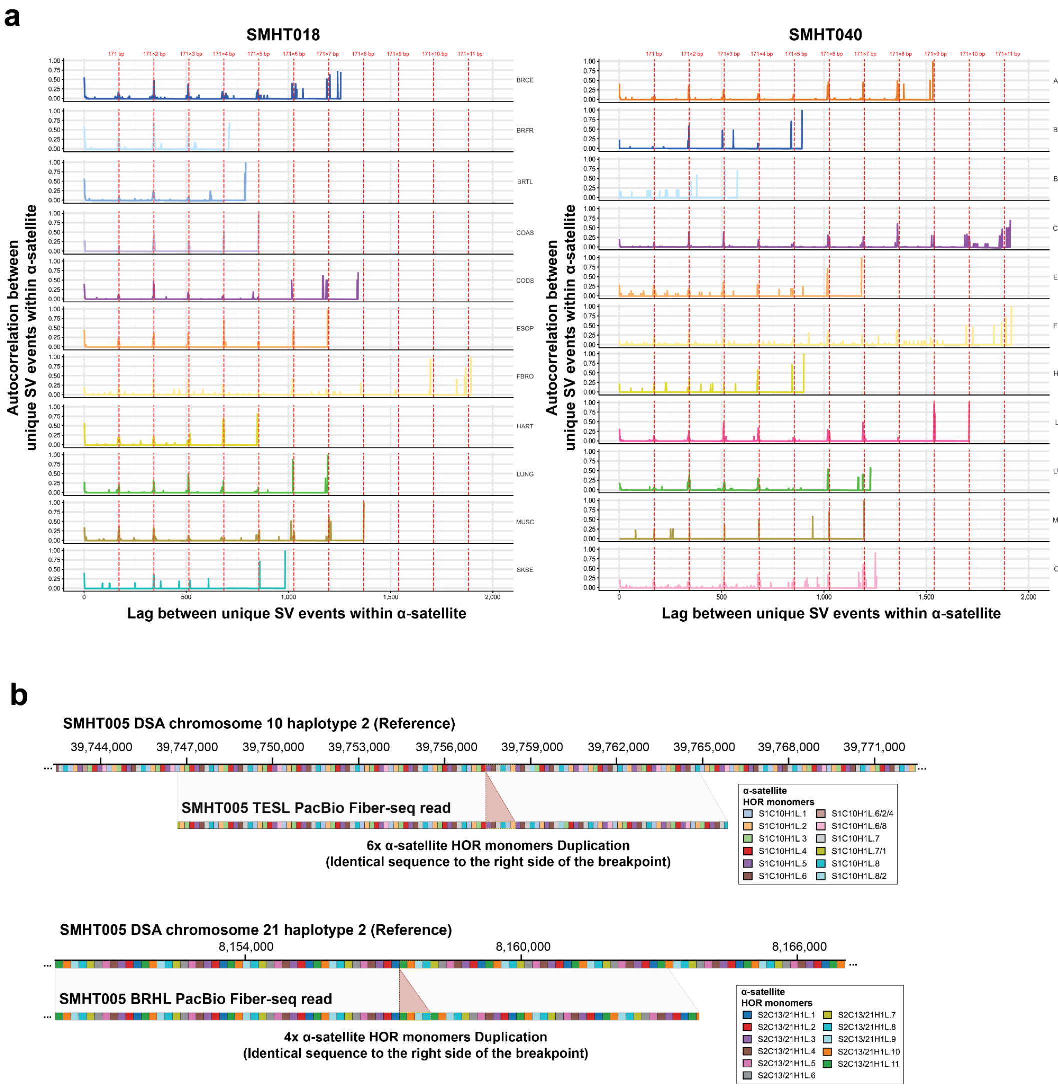


**Figure S22 | Identification of sSVs within centromere alpha-satellite arrays, related to Figure 6d.**

**a,** Autocorrelation of unique sSV events in α-satellite regions of centromeres across different tissues of two example donors (SMHT018 and SMHT040), identified using PacBio sequencing data, showing a periodic pattern at multiples of 171 bp, the size of an α-satellite monomer.

**b,** Genome browser view illustrating mosaic sSVs in TESL (top, chromosome 10) and BRHL (bottom, chromosome 21) from donor SMHT005, showing perfect duplications of 6x and 4x α-satellite HOR monomers, respectively. The inserted sequence is identical to the right of the breakpoint in both cases. Each colored box represents different α-satellite HOR monomer with mismatches between individual reads and the DSA indicated.

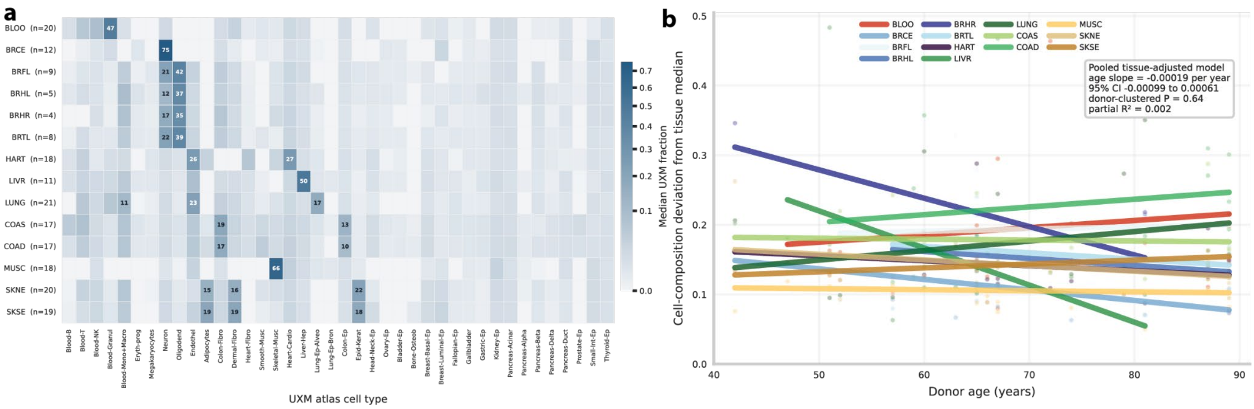


**Figure S23 | UXM-estimated cell-type composition and its relationship with donor age, related to Figure 7a.**

**a,** Median UXM-estimated cell-type fractions across the 16 tissues retained for composition analysis. Values ≥10% are labeled; **b,** Cell-composition deviation from the tissue-specific median across donor age. Points represent donor–tissue observations, and colored lines show tissue-specific linear fits. The pooled tissue-adjusted model showed no significant age association (donor-clustered P=0.51; $R^2$=0.004). Tissues with defining cell types not represented in the UXM atlas were excluded.

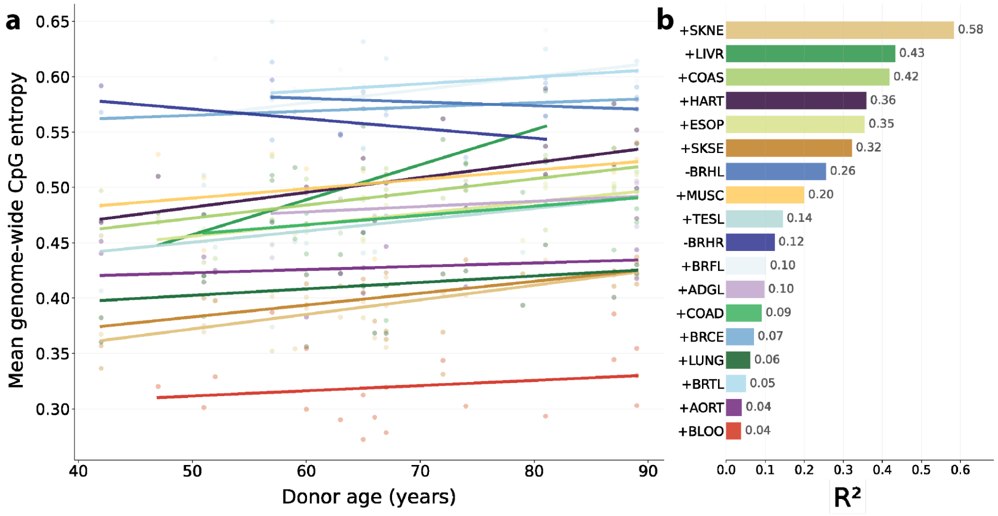


**Figure S24 | Tissue-specific associations between DNA methylation entropy and donor age, related to Figure 7a.**
Points represent individual samples, and colored lines show donor-level linear fits within each detailed tissue after averaging replicate samples from the same donor and tissue. The right panel shows Pearson $R^2$ for each tissue; "+" and "−" indicate positive and negative slopes, respectively. Only tissues represented by at least four donors were included.

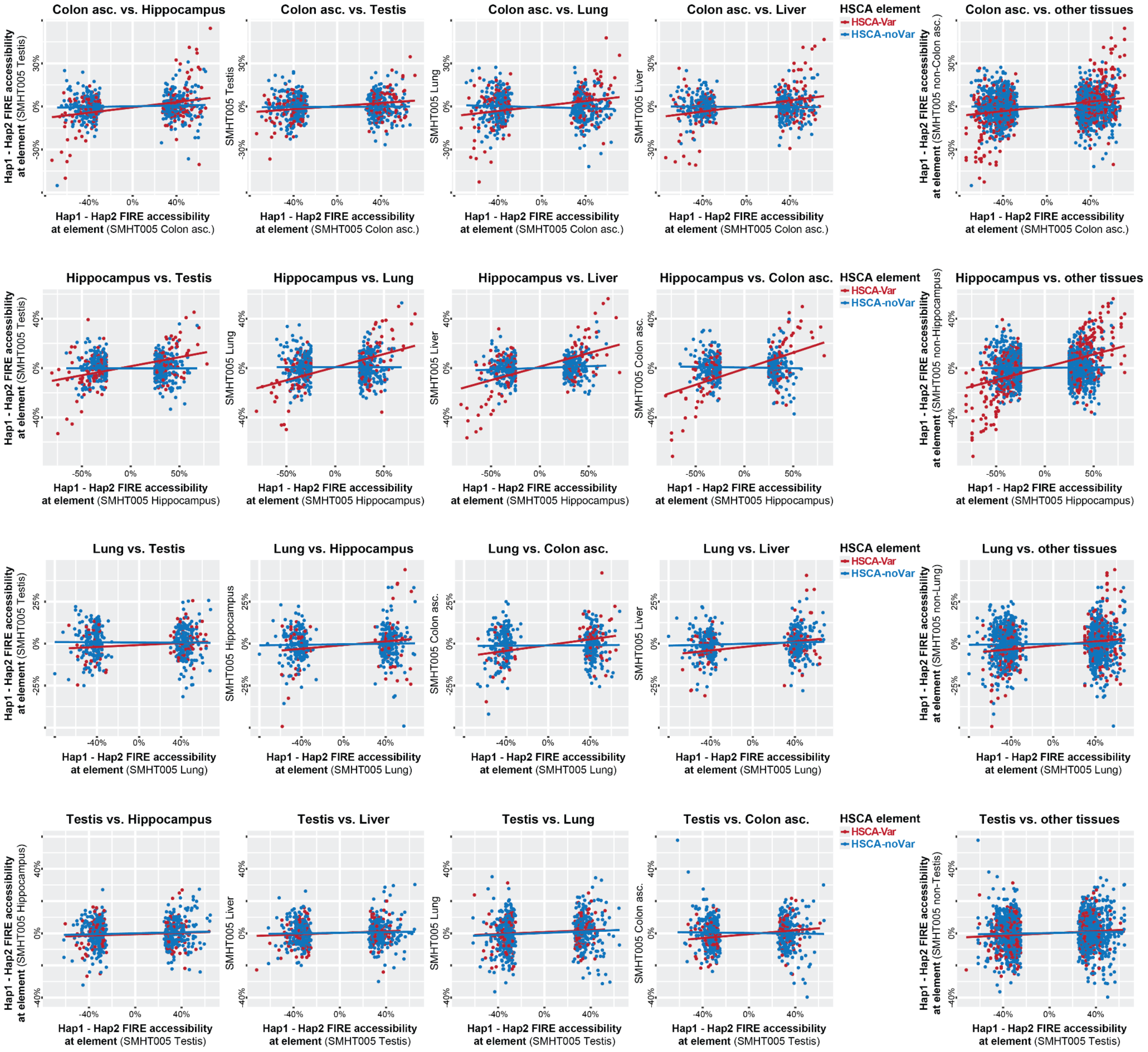


**Figure S25 | HSCA consistency across tissues from SMHT005, related to Figure 7c.** Scatter plots showing the difference in chromatin accessibility along haplotype 1 and haplotype 2 at elements that were identified as having haplotype-selective chromatin accessibility within (top) SMHT005 Colon ascending (second from top) SMHT005 Hippocampus (second from bottom) SMHT005 Lung (bottom) SMHT005 Testis using a Fisher exact test p-value cutoff of 0.005. Axis indicate the SMHT005 tissue chromatin data that is displayed. Elements are colored based on whether they were labeled as HSCA-Var or HSCA-noVar. Linear fits to the HSCA-Var or HSCA-noVar elements are shown for each plot. (right) For each of the rows, shown is a combination plot.

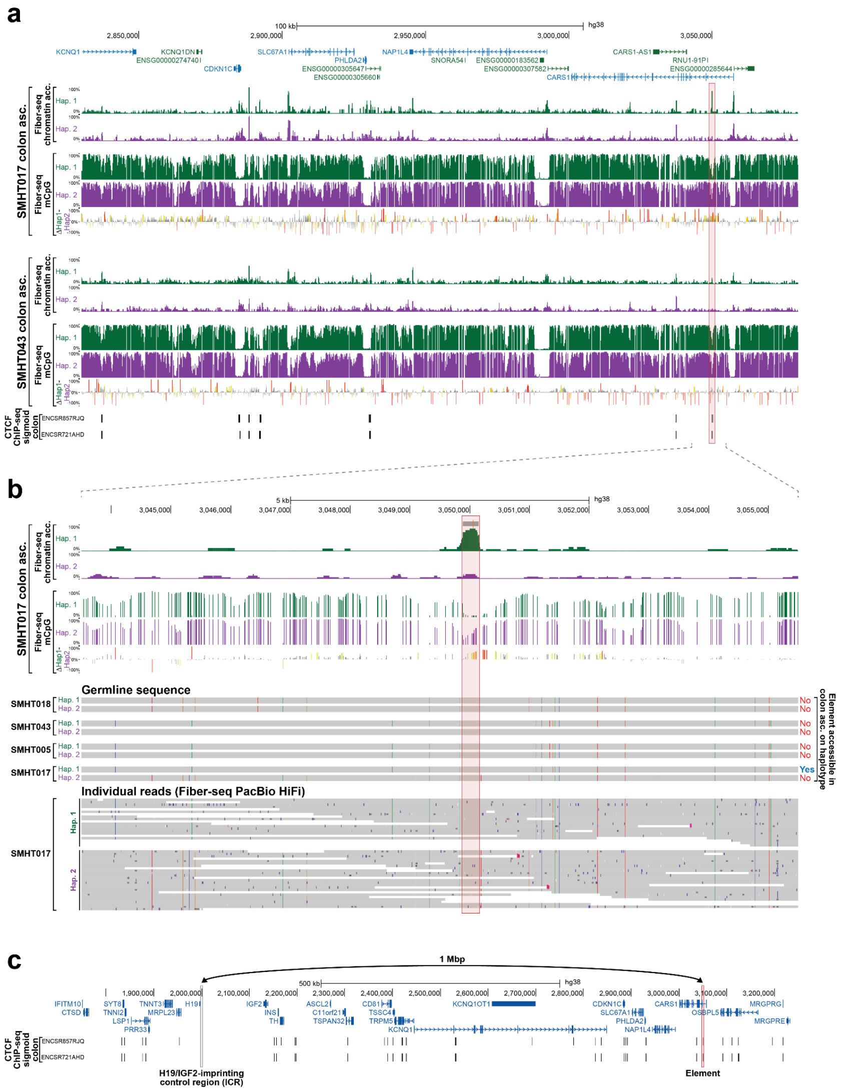


**Figure S26 | Stochastic epigenetic state along chromosome 11p, related to Figure 7d.**
**a,** Ascending colon Fiber-seq data from both SMHT017 and SMHT043 showing haplotype-resolved chromatin accessibility and 5mCpG methylation information within a 250 kbp window surrounding the element highlighted in Figure 7d. ENCODE CTCF peaks shown at bottom. **b,** (top) Ascending colon Fiber-seq data from SMHT017 showing haplotype-resolved chromatin accessibility and 5mCpG methylation information within a 12 kbp window surrounding the element highlighted in Figure 7d. (middle) Haplotype-resolved germline genetic information from from SMHT018, SMHT043, SMHT005, and SMHT017, as well as indication on right as to whether that haplotype shows chromatin accessibility in the ascending colon. Note that the haplotype selectively accessible in SMHT017 shows strong local sequence similarity to haplotypes in SMHT018, SMHT043, and SMHT005 that lack chromatin accessibility in ascending colon. (bottom) Individual Fiber-seq PacBio HiFi sequencing reads from SMHT017 ascending colon showing lack of a somatic genetic variant at the element of interest. **c,** Zoom-out of region encompassing the element highlighted in Figure 7d, showing that it is located 1 Mbp away from the H19/IGF2-imprinting control region (ICR). ENCODE CTCF peaks shown at bottom.

### Supplementary Table Legends

**Table S1 | Overview of the first 25 SMaHT Donor Production Data generated by Centers**
Detailed overview of the first 25 SMaHT production donor data generated by institutions in the SMaHT Network. Target sequence coverage and target read numbers are indicated for WGS and RNA-Seq assays, respectively. GCC = Genome Characterization Center; TTD = Technology & Tool Development Groups.

**Table S2 | Germline variants from 25 SMaHT donors**
Germline variants from the 25 donors were screened against the American College of Medical Genetics (ACMG) Secondary Findings Work Group list v3.2 and classified using ClinVar. Pathogenic or likely pathogenic heterozygous variants were identified in BRCA2 (one donor), MUTYH (three donors) and KCNQ1 (one donor). No donor carried biallelic pathogenic variants in a recessive gene on the ACMG list.

**Table S3 | Gene overlapping somatic TR instability sites**
sSV sites overlapping genes (AnnotSV) and tandem repeats (adotto TR catalog v2.1) were inspected (Supplementary Methods) to identify seven loci with somatic TR instability in at least two donors in predominantly one tissue, with SKI indicating both calf and abdominal skin sites.

**Table S4 | sSNVs annotated as stop gain variants**
The table lists sSNVs annotated by VEP as stop-gained variants as well as the pLI of the gene. It includes each sSNVs ID, sample, tissue, and affected gene.

**Table S5 | Predicted functional sSNVs by deep learning tools**
The table lists sSNVs predicted to have functional effects by SpliceAI, PromoterAI, AlphaGenome, and APARENT2. It includes sSNVs that passed the specified cutoffs, their predicted target genes, and the scores generated by each tool. The cutoffs were a SpliceAI delta score > 0.2 and scores above the 95th percentile for PromoterAI, AlphaGenome, and APARENT2.

**Table S6 | Detection of developmental sSNVs**
Table showing the median detection rates for developmental sSNVs, corresponding to each group in Figure 5C

**Table S7 | Quality control metrics for DSAs**
Quality control metrics for the DSAs.

### Supplementary Methods

#### Supplementary Methods Section 1 | Donors and tissue collection

##### S1.1 Full donor inclusion and exclusion criteria

*Eligibility criteria:* Donor eligibility was determined by the partner OPO utilizing screening SOPs developed for SMaHT. Donors over the age of 18 years were considered eligible for inclusion if the following criteria were met: no history of HIV, Hepatitis C, or Hepatitis B, no history of IV drug use in 5 years prior to death, no chemotherapy or radiation treatment in 24 months prior to death, no known chromosomal or genetic disorder, no active or history of metastatic cancer, no diagnosis of multisystem organ failure at time of death, not a recipient of an organ or allogeneic bone marrow transplant, no whole blood transfusion in 48 hours prior to death, and no positive blood cultures or sepsis at the time of death. Eligibility for brain donation also required that the donor did not have a cause of death related to brain injury or head trauma and was not declared brain dead or was ventilator dependent for more than 24 hours prior to death. Donors could be included in SMaHT for all tissues other than brain if the brain-specific exclusion criteria were not met. Tissue recovery had to be completed within 24 hours of cross-clamp or cardiac cessation for SMaHT eligibility[1].

##### S1.2 Per-donor tissue inventory and anatomical sampling schema

*Biospecimen collection:* Tissues for the SMaHT study were recovered from postmortem donors by the Tissue Procurement Center's (TPC) network of three organ procurement organizations (OPOs). SMaHT-specific standard operating procedures (SOPs) for donor screening, authorization for tissue donation, biospecimen procurement and preservation were initiated at each partner OPO. Briefly, samples of up to fifteen distinct tissue sites and whole brain were requested for recovery from each eligible donor[2]. The SMaHT tissue sampling design is categorized by tissue type: blood, buccal swab, whole organ (brain), solid organ (muscle, adrenal gland, lung, heart, liver, gonads), and mucosal tissue (aorta, esophagus, colon, and skin). Twelve cc of whole blood was recovered from each donor in two EDTA blood tubes and frozen in 1 cc aliquots on dry ice. Two buccal swab samples were recovered from each donor using 4N6FLOQSwabs (QIAGEN). Swabs were frozen in a -80°C freezer 48-72 hours after collection. Brain was recovered whole and placed on wet ice for shipment to University of Maryland Brain and Tissue Bank (UMBTB) for processing. Solid organs were collected as 1.5 cm thick sections of variable lengths depending on tissue size then divided into multiple aliquots: 3-8 frozen aliquots (each 1x1.5x1cm) and 1-6 fixed aliquots (each 1x1.5x0.5cm) per tissue site. Mucosal tissues were collected as 7.5x1.5cm x full thickness tissue sections then divided into multiple aliquots: 6 frozen aliquots (each 1x1.5cm x full thickness) and 3 fixed aliquots (each 0.5x1.5cm x full thickness) per tissue site. Spatial orientations of all tissue aliquots were maintained in tissue cassettes via recovery and labelling SOPs. All tissue samples were trimmed of excess fat and rinsed with saline prior to aliquoting and preservation. Frozen tissues were preserved using a thermal tray on a bed of dry ice. The tissue samples were placed into cryosettes and placed on the thermal tray for a minimum of ten minutes to allow adequate tissue freezing. Frozen tissues were then stored in a -80°C freezer. Fixed tissue samples were placed in tissue cassettes and placed in 10% formalin for histopathology.

*Brain sampling:* Four regions of brain (cerebellum, frontal lobe, temporal lobe, and hippocampus) were sampled by UMBTB upon receipt. Six frozen aliquots (each 1x1.5x1cm) and one fixed slice were sampled from each brain region. Frozen tissue aliquots were placed on a cold tray and submerged briefly in a bath of 2-methylbutane sitting over dry ice until completely frozen, then placed in cryosettes and stored in a -80°C freezer. Fixed tissue slices were placed in tissue cassettes in 10% formalin for neuropathology.

*Tissue aliquot subsampling for distribution:* Frozen tissue aliquots were photographed on a bed of dry ice prior to subsampling to document gross morphology and map the extraction of multiple 3.0mm tissue cores for distribution. Tissue core extraction was performed using the CXT 353 Bench-Top Frozen Sample Aliquotter (Basque Engineering, West Newbury, MA). The machine was pre-cooled to −80 °C with liquid nitrogen ($LN_2$) according to manufacturer-recommended parameters and $LN_2$ levels were maintained during extraction to prevent warming. Pre-labeled tubes and 3.0mm probes (NBS Scientific, Cannonsburg, PA) were chilled within the instrument prior to use. Extraction probe speed and depth were optimized for each tissue type. Guided by a laser positioning system and the predefined digital extraction map, 3.0mm tissue cores were extracted from the tissue aliquot cryosette and dispensed into corresponding pre-labeled cryovials. Each tissue aliquot was photographed again following tissue core extraction to document the location of each recovered tissue core for any future spatial analyses.

*Fibroblasts sample collection and generation:* A full-thickness skin sample was collected from the leg below the back of the knee on the lateral side from each donor and shipped on ice in media (MEM with HEPES supplemented with 10% bovine calf serum and 100ug/ml Gentamicin/Amphotericin B (all reagents from Gibco)) to UMBTB. Fibroblast primary cultures were expanded as previously described[3] using fibroblast culture media (M199:M106 medium (1:1; Invitrogen) supplemented with 15% fetal bovine serum (FBS; ATCC), 0.4ug/ml Hydrocortisone (Sigma), 10ng/ml EGF (Invitrogen), 0.25ug/ml Fungizone (Invitrogen), and 50ug/ml Gentamicin/PenStrep (Invitogen)). Cells were passed twice using 0.25% trypsin-EDTA (Invitrogen). One million cells per vial were cryopreserved with 5% DMSO.

**S1.3 Biospecimen QC metrics**

*Histopathology processing and review:* All fixed tissues were paraffin embedded for histopathology review as a quality control (QC) tissue measure and to further characterize tissue composition of recovered tissues. A hematoxylin-eosin-stained slide was generated from each formalin-fixed paraffin embedded (FFPE) block. All slides were digitized with whole-slide imaging (WSI) at 40x magnification on an Aperio slide scanner and evaluated by a pathologist at UMBTB. The TPC utilized a standardized approach to collect histopathological metadata to describe the cellular composition of tissue samples, described in[4]. Pathology metadata for each fixed tissue section includes autolysis score, percent target tissue present, percent non-target tissue present, and any unexpected pathological findings (i.e., necrosis, inflammation, etc). For neuropathology, a dementia and age-related immunohistochemical evaluation for amyloid-β, pTau, α-synuclein, and TDP-43 was performed for donors over the age of 50 years. An initial QC evaluation includes verification of the correct target tissue is present. If a mismatch between the expected target tissue type and the slide histology is identified during the initial pathology QC review and cannot be resolved, the tissue is removed from the SMaHT biorepository for downstream use. Histopathology review may also deem a tissue unacceptable if there is <10% of target tissue present in tissue section or diffuse tissue abnormalities indicative of a pathology process.

**S1.4 Donor metadata fields (age, sex, ancestry, environmental exposures)**

*Donor and tissue metadata collection:* Donor clinical data were collected by the OPO and include donor demographics (age, sex, height, weight, and body mass index (BMI)), medical history (cause of death and past medical history), social history (alcohol, tobacco, and illicit drug use), and serological testing results. Tissue sample data collected include the tissue type, ischemic time, and pathology detail. In addition, UBERON ontologies were utilized to standardize anatomical locations for tissue sites.

### Supplementary Methods Section 2 | Sample preparation and sequencing

#### S2.1 Nucleic acid and nuclei extractions

DNA and RNA were extracted from blood, tissue, fibroblast and buccal samples across different GCCs using validated workflows. RNA was extracted from frozen tissues using column-based purification methods with on-column DNase treatment to remove genomic DNA contamination. For single-nucleus RNA-seq and ATAC-seq applications, frozen tissue samples were mechanically dissociated in an extraction buffer followed by magnetic-bead-based enrichment.

**DNA extraction and nuclei isolation protocols - Short-read**

*[GCC-BCM] DNA Extraction:* Tissue cores were homogenized in lysis buffer with 1.4 mm Ceramic Beads (Omni International) or cut with a razor blade and incubated overnight with Proteinase K. DNA was extracted from the homogenized tissues and blood using the Chemagic Prime 8 robot and Chemagen's proprietary Magnetic Bead technology, with the Chemagic Prime DNA Blood kit and the manufacturer's protocol (Revvity).

*[GCC-BCM] RNA Extraction:* Tissue cores were homogenized in TRIzol with 1.4 mm Ceramic Beads (Omni International). RNA from tissue homogenate and blood was manually extracted according to the manufacturer's Trizol protocol. The isopropanol/aqueous phase was then transferred to a RNeasy spin column (Qiagen), and the manufacturer's RNeasy protocol was then followed.

*[GCC-UW-SCRI] DNA Extraction:* DNA was extracted from tissue and fibroblast samples using the Qiagen DNeasy Blood and Tissue kit (69506) following the manufacturer's instructions. Lysis time varied by tissue type and ranged from 4 hours to overnight. DNA from 1mL blood samples were extracted using the Qiagen MagAttract HMW DNA kit (67563) following the manufacturer's instructions. The QIAmp DNA Mini kit (51304) was used to purify DNA from buccal swabs, following the manufacturer's instructions. The integrity of the extracted DNA was confirmed using the Agilent Femto Pulse instrument and the DNA yield was confirmed using the Invitrogen Qubit kit and Qubit 4 fluorometer.

*[GCC-NYGC] DNA and RNA extraction:* DNA and RNA for short-read bulk work were extracted using Qiagen's AllPrep kit (catalog number 80204), following manufacturer instructions. Additional RNA was extracted if initial core extraction was low RIN, using Qiagen's RNeasy Mini kit (catalog number 74106).

*[GCC-Broad] DNA Extraction:* For tissue-derived samples, genomic DNA was extracted from fresh-frozen tissue (20–25 mg per sample) using the Qiagen AllPrep DNA/RNA Universal Kit; total RNA was co-extracted but was not used for downstream sequencing. Tissue was cut to mass under continuous cold-chain handling and placed into a pre-chilled 2 mL round-bottom tube containing a 5 mm stainless steel bead, then homogenized in Buffer RLT Plus (supplemented with 14.3 M β-mercaptoethanol) using a TissueLyser II operated at 25 Hz for 4 minutes. Lysate was clarified by centrifugation and passed through an AllPrep DNA spin column to bind genomic DNA under high-salt conditions. The column-bound DNA was washed sequentially with Buffer AW1 (including a 5-minute room-temperature proteinase K incubation) and Buffer AW2, then eluted in Buffer EB. All tissue handling was performed on dry ice to preserve nucleic acid integrity. Purified DNA was transferred to barcoded Matrix tubes for downstream quantification and processing.

For whole blood samples, genomic DNA was instead extracted using the Promega Maxwell HT 96 gDNA Blood Isolation System (proteinase K, Cell-Lysis Buffer, paramagnetic Resin, Binding Buffer, and Wash Buffer) on a Hamilton STARlet liquid handler, a bead-based method that uses magnetic particle capture in place of centrifugation or vacuum filtration. 800 μL of thawed whole blood was aliquoted for automated processing, with 350 µL of this volume carried into extraction. Samples were lysed with proteinase K and Cell-Lysis Buffer (20 minutes at 75°C with shaking at 1,200 rpm), and DNA was captured on the paramagnetic Resin in the presence of Binding Buffer across two rounds of binding, each followed by magnetic separation. Bound DNA was washed three times with Wash Buffer and 50% ethanol, with shaking at 1,200 rpm and magnetic separation between washes, dried briefly at 70°C, and eluted in 10 mM Tris–1 mM EDTA Buffer across two sequential elution steps at room temperature. The final eluate (~150 µL) was transferred from the processing plate to barcoded matrix tubes and stored at +4°C.

*[GCC-WashU-VAI] DNA Extraction:* DNA from blood: DNA was isolated on the Promega Maxwell RSC 48 using the Maxwell RSC Blood DNA Kit (Promega AS1400), following the manufacturer's protocol. High molecular weight DNA from tissue: DNA was isolated with the Qiagen MagAttract HMW DNA Kit (Qiagen 67563), following the manufacturer's protocol. DNA from tissue: DNA was isolated with the Qiagen DNeasy Blood and Tissue Kit (Qiagen 69504), following the manufacturer's protocol. Samples were homogenized in Buffer ATL using the Qiagen Tissuelyser II for 20 seconds at 15 Hz prior to the addition of Proteinase K. The optional RNaseA treatment step was included.

**DNA extraction and nuclei isolation protocols - Long-read PacBio**
*[GCC-UW-SCRI]:* Cultured fibroblasts were harvested and snap-frozen. Cell pellets were thawed and split to ~5M cells per reaction prior to HMW DNA extraction using the Monarch HMW DNA Extraction Kit (NEB, T3050L) following manufacturer's recommendations but using a lysis shaking speed of 1000 rpm.

*[GCC-Broad]:* High molecular weight (HMW) genomic DNA was extracted from frozen tissue using the PacBio Nanobind PanDNA Kit. Tissue was homogenized in Buffer CT using a TissueRuptor at maximum speed for 10 seconds, pelleted by centrifugation (6,000 × g, 4°C, 5 minutes), and washed with additional Buffer CT before digestion with proteinase K and Buffer CLE3 for 30 minutes at 55°C with agitation (900 rpm; extended up to 2 hours if lysis was incomplete), followed by RNase A treatment under the same conditions. Buffer SB was added to the digest, and high molecular weight (HMW) genomic DNA was precipitated with isopropanol in the presence of a Nanobind magnetic disk, with binding promoted by 15 minutes of end-over-end mixing (HulaMixer, 20 rpm) at room temperature. Disk-bound DNA was washed on a magnetic rack with two rounds each of Buffer CW1 and Buffer CW2 and eluted in Buffer LTE following a 10-minute room-temperature incubation, yielding high-purity, HMW DNA. The eluate was transferred to barcoded matrix tubes and stored at +4°C pending downstream quantification.

**DNA extraction and nuclei isolation protocols - Long-read ONT**
*[GCC-NYGC] HMW DNA extraction:* HMW DNA was extracted with the Monarch HMW Extraction kit from New England Biolabs (catalog numbers T3050L and T3060L).

**Tissue Fiber-seq reactions and DNA extractions**
*[GCC-UW-SCRI]:* One or two brain tissue cores from the same donor and brain region were mixed together with 1mL of fresh homogenization buffer (250mM Sucrose, 15 mM Tris-Cl, pH 8.0, 15 mM NaCl, 60 mM KCl, 1 mM EDTA, pH 8.0, 0.5 mM EGTA, pH 8.0, 0.5 mM Spermidine, 0.1 mM DTT, 0.1% Triton X-100 1x Protease inhibitor, 0.2U RNasein plus) within a 5mL Dounce homogenizer on ice and homogenized using 10 strokes of pestle A followed by 10 strokes of pestle B. Samples were then filtered through a 70 micron filter and transferred to a 15ml low bind tube. 15 µl of activated Concanavalin A (ConA) beads (Cell Signaling #93569) was added to the sample, and the sample was then centrifuged at 345 g for 5 minutes at 4°C, and the

supernatant was removed. The pellet, which contains the nuclei and ConA beads, was then resuspended in Buffer A (15 mM Tris-Cl, pH 8.0, 15 mM NaCl, 60 mM KCl, 1 mM EDTA, pH 8.0, 0.5 mM EGTA, pH 8.0, 0.5 mM Spermidine), with 0.8 mM S-adenosylmethionine (SAM) (New England Biolabs B9003S) and 1.5 units/µl of Hia5 enzyme[5]. Based on the size of the pellet, this reaction was performed in a total volume of either 60 µl, 120 µl, 180 µl, or 240 µl. The reaction was performed at 10 minutes at 25C and stopped by adding sodium dodecyl sulfate (SDS) to a final concentration of 1%. Non-brain tissues were processed in a similar manner with the exception that prior to dounce homogenization, the tissue cores were placed in a ceramic mortar containing liquid nitrogen and then pulverized with the pestle while still frozen. Additional liquid nitrogen was added to the mortar, and the submerged pulverized tissue was scrapped off the mortar and then poured into the dounce homogenizer while still suspended in liquid nitrogen. The sample was then briefly thawed in the dounce homogenizer prior to the addition of homogenization buffer as above. For testis samples specifically, dithiothreitol (DTT) was added to the sample after the SDS addition at a final concentration of 1.6 mM DTT.
DNA was then extracted using either the Promega Wizard HMW Extraction kit (Promega A2920) per the manufacturer protocol, or the QIAGEN MagAttract HMW DNA Kit (QIAGEN Cat no. 67563) per the manufacturer protocol with slight modifications as described below. Specifically, for the QIAGEN MagAttract HMW DNA elution step, beads were incubated in Buffer AE (10mM Tris, pH 9.0, 0.5 mM EDTA) at 4°C overnight or over the weekend. After 4°C incubation, they were incubated for 10 min at 56°C at 1000 RPM, gently mixed using a wide-bore tip 5-6 times, placed on magnet, and then the supernatant was transferred to a new 1.5 mL LoBind tube.

**RNA extraction protocols**
*[GCC-UW-SCRI]:* Tissue cores were incubated in RNAlater-ICE Frozen Tissue Transition Solution (InVitrogen, AM7030) overnight at -20°C. Tissues were extracted with ReliaPrep RNA Miniprep Systems (Promega, Z6014) following manufacturer protocol with slight modifications as described below. Tissues were removed from RNAlater-ICE, supernatant discarded, and disrupted in 500 µl LBA buffer using motorized tissue grinder in a 1.5ml tube. The homogenized tissue was pipetted up and down 7-10 times then loaded onto a QiaShredder column (QIAGEN, 79656) and centrifuged at 16,000 rcf for 3 minutes. 500 µl of RDB buffer was added to the flow through, capped, vortexed for 10 seconds, incubated for one minute at room temperature before centrifuging at 10,000 rcf for 3 minutes. The supernatant was transferred to a new 1.5ml tube. 340 µl of isopropanol was added and the sample was vortexed for 5 seconds. 700ul of the homogenate was then transferred at a time to a ReliaPrep™ Minicolumn, which was centrifuged at 12,000 rcf for 30 seconds, followed by discarding the flow through. The remaining homogenate was applied to the same column, which was centrifuged at 12,000 rcf for 30 seconds, followed by discarding the flow through. 500µl of RWA was then added to the column, which was centrifuged at 12,000 rcf for 30 seconds, followed by discarding the flow through. 30 µl of prepared DNase buffer (24µl Yellow Core Buffer, 3µl MnCl2, 0.09M, 3µl DNase enzyme) was then added to the column and incubated for 15 minutes at room temperature. 200µl of Column Wash Solution was then added to the column, which was then centrifuged at 12,000rcf for 15 seconds. 500 µl of RWA was then added to the column, which was then centrifuged at 12,000rcf for 15 seconds, followed by discarding the flow through. The column was then transferred to a new collection tube, 300µl of RWA was added, which was then centrifuged at 16,000 rcf for 2 minutes. The column was then transferred to an elution tube, 33µl of Nuclease-Free Water was added, which was then centrifuged at 12,000 rcf for 1 minute, and the flow through was saved.

*[GCC-Broad]:* Total RNA was extracted from frozen tissue (10–17 mg per sample) using the Promega ReliaPrep RNA Tissue Miniprep System. Tissue was homogenized in a guanidine

thiocyanate/1-thioglycerol lysis Buffer (LBA + TG Buffer) using a TissueLyser II operated at 25 Hz for 5 minutes, cleared by centrifugation (10,000 × g, room temperature, 3 minutes; repeated at 14,000 × g if the lysate remained turbid), and mixed with isopropanol before being bound to a ReliaPrep silica minicolumn by centrifugation (12,000–14,000 × g). Bound RNA was treated with a freshly prepared on-column DNase I mixture (DNase I enzyme with Yellow Core Buffer and MnCl2) for 15 minutes at room temperature to remove genomic DNA, washed sequentially with Column Wash Solution and RNA Wash Solution, and eluted in nuclease-free water. The eluate (50 µL) was transferred to a barcoded 0.75 mL matrix tube and stored at –80°C pending downstream quantification and processing.

*[GCC-WashU-VAI]:* RNA from tissue: RNA was isolated with the Promega ReliaPrep RNA Miniprep System (Promega Z6014), following the manufacturer's protocol for fibrous tissues. The samples were homogenized in the LBA + TG buffer using the Qiagen Tissuelyser II for 4 minutes at 20 Hz, prior to the addition of buffer RDB.

**S2.2 Library preparation details - Illumina**

*[GCC-BCM]:* Two independent methods (Picogreen assay and 1% E-gels) were used to determine the quantity and quality of the DNA before library construction. For library preparation, DNA (1 ug) was sheared into fragments of approximately 450-600 bp in a Covaris E220 system (Covaris, Inc. Woburn, MA) in batches of 96 samples at a time, followed by double SPRI bead clean up to select a narrow band of sheared DNA for library preparation. DNA was end-repaired, 3'-adenylated and ligated using a set of 96 8-bp adapters (Illumina TruSeq UD Indexes v2, # 20040870) for sample barcoding. The final library size estimation and quantification are completed using the Fragment Analyzer (Agilent AAT, Inc) electrophoresis system and QuantStudio™ 6 Flex Real-Time PCR System (Applied Biosystems) respectively to achieve an average final library size of ~530bp and must be greater than 470 bp. Libraries were sequenced on The NovaSeqX instrument to generate 150 bp, dual indexed and paired-end sequence reads in a format of multiplexed pools to generate 412× - 541× coverage.

Mapping and QC Pipeline for Internal Quality Assessment Before Data Submission to DAC WGS sequence data were aligned to the hg38 reference genome, followed by variant calling using Illumina's Dynamic Read Analysis for GENomics (DRAGEN) software, v4.3.6. Genome coverage was evaluated by calculating the mean coverage and the distribution of coverage across the genome, including the proportions of bases covered at ≥1×, ≥10×, and ≥20×. The proportions of bases meeting these thresholds and the total number of mapped bases at Q20 or higher were reported for internal QC tracking. Alignment data were assessed for contamination using VerifyBamID v1.1.3 and an orthogonal confirmation of sample identity was applied using the Error Rate In Sequencing (ERIS) software developed at the HGSC to rapidly compare sequence data to genotypes from SNP arrays via an "exact match" test.

*[GCC-UW-SCRI]:* Starting with a minimum of 750ng of DNA, samples are sheared in a 96-well format using a Covaris R230 focused ultrasonicator targeting 380bp inserts. This insert size improves overall library performance and allows the longer sequencing read lengths on Illumina sequencing platforms (150bp) to be efficiently used without producing a significant number of overlapping reads. The resulting sheared DNA is cleaned with Takara NucleoMag beads to remove sample impurities prior to library construction. Shearing is followed by size selection and sample prep is performed using the KAPA Hyper Prep kit (KR0961 v1.14). End-repair, A-tailing, and ligation are performed as directed. Two final NucleoMag cleanups are performed after ligation to remove excess adapter dimers from the library. All library construction steps are automated on the Revvity Janus platform. Library yield is quantified using Invitrogen Quant-IT

dsDNA High Sensitivity kit (Q33120). Libraries are validated in triplicate using the Biorad CFX384 Real-Time System and KAPA Library Quantification Kit (KK4824).

Barcoded libraries are pooled using liquid handling robotics prior to loading. Massively parallel sequencing-by-synthesis with fluorescently labeled, reversibly terminating nucleotides is carried out on the NovaSeq X Plus sequencer. Base calls are generated in real-time on the instrument (RTA 4.29.3) and then demultiplexed, fastq files are produced by bcl-convert v4.2.7.

*[GCC-NYGC]:* Whole-genome sequencing (WGS) libraries were prepared using the NEBNext Ultra II FS DNA PCR-free Library Preparation Kit (NEB E7430L) in accordance with the manufacturer's instructions. 500ng of DNA was sheared enzymatically and was subsequently end-repaired and adenylated. DNA fragments were ligated to Illumina sequencing adapters and the libraries underwent bead-based size selection. Final libraries were quantified using the QuantStudio5 Real-Time PCR System (Applied Biosystems) and Fragment Analyzer (Agilent). Short-read WGS libraries were sequenced on the Illumina NovaSeq X Plus, using 2 x 150 bp cycles.

*[GCC-Broad]:* PCR-free whole-genome sequencing libraries were prepared according to the KAPA HyperPrep PCR-free manufacturer's protocol. Genomic DNA was quantified by PicoGreen fluorometric assay and normalized to a standard input concentration. DNA was mechanically sheared using a Covaris LE220-Plus instrument (200 W peak incident power, 50 cycles per burst, 25% duty factor, 60-second duration) to a mean fragment size of approximately 450 bp, compatible with 2 × 151 bp paired-end sequencing. Sheared DNA underwent end repair and A-tailing (KAPA End Repair/A-Tailing Master Mix; 30 minutes at 20°C followed by 30 minutes at 65°C), followed by ligation of Illumina indexed adapters (15 minutes at 20°C). KAPA HyperPure SPRI bead cleanup was performed after each enzymatic step, including dual-sided SPRI/PEG–NaCl size selection to remove adapter dimers and residual small fragments. No PCR amplification was performed.

Final libraries were quantified by qPCR against Illumina adapter sequences using the KAPA Library Quantification Kit (SYBR-based standard-curve assay on a QuantStudio 7 instrument), requiring a standard-curve $R^2 > 0.98$ and amplification efficiency of 90–110% to confirm library concentration and exclude failed libraries (<0.9 nM). Libraries were normalized to 1.8 nM and pooled according to the target sequencing coverage (24-plex across four lanes for NovaSeq 6000 S4 flow cells or 47–48-plex across six lanes for NovaSeq X 25B flow cells). Pools were re-quantified by qPCR and renormalized as needed before sequencing on Illumina NovaSeq 6000 or NovaSeq X instruments using 2 × 151 bp paired-end reads. Two independent libraries were prepared from each DNA sample and each sequenced to approximately 80× coverage, yielding a combined target coverage of 160×. Sequencing runs were required to meet predefined quality-control thresholds before data release, including the minimum number of reads passing filter for the corresponding flow-cell configuration, ≥80% of bases with a quality score of Q30 or higher, and ≥70% of clusters passing filter and occupied. Data were delivered as de-multiplexed, aggregated CRAM files with accompanying CRAI index and MD5 checksum files.

*[GCC-WashU/VAI]* Genomic DNA samples were quantified using the Qubit Fluorometer. Genomic DNA (~600-1000ng) was fragmented on the Covaris LE220 instrument targeting ~375bp inserts. Fragmented DNA was size selected using 0.8X ratio of Ampure XP beads (Beckman Coulter) to remove fragments less than 300bp. Dual indexed libraries were constructed utilizing the KAPA Hyper PCR-free library prep kit (Roche Diagnostics, Cat # 7962371001). Full length custom adaptors were used during ligation (IDT, UDI/UMI configuration with 10bp UDIs and a 9bp UMI in the i7 position). Libraries were run with KAPA

Library Quantification kit (Roche Diagnostics) to measure molar concentration. Libraries are sequenced on NovaSeq X using paired end reads extending 150bp. For this application targeting 500X coverage, 4 libraries were constructed and utilized to generate the >500X coverage. For both Kapa Hyper PCR-free libraries as well as the Bulk RNA-seq libraries, the molarity of each library was accurately determined through qPCR utilizing the KAPA library Quantification Kit according to the manufacturer's protocol (KAPA Biosystems/Roche) to produce cluster density appropriate for the Illumina NovaSeq X Plus instrument. Normalized libraries were sequenced on a NovaSeq X Plus Flow Cell using the151x19x10x151 sequencing recipe according to manufacturer protocol to generate a >500X WGS coverage, and >100M read pairs for the RNAseq library.

**S2.3 Long-read sequencing platform allocation**

Long-read sequencing was distributed across GCCs by platform, depending on the sequencing platforms available at each center. PacBio HiFi data were generated at the Broad Institute, the University of Washington (UW) and Washington University (WashU), while ONT data were generated at UW, Baylor College of Medicine (BCM) and the New York Genome Center (NYGC). Target coverage also varied by center, with the Broad Institute performing PacBio sequencing at a target depth of 12×, while all other centers targeted 50×. ONT sequencing was performed on PromethION instruments using R10.4.1 flow cells, and PacBio HiFi sequencing on Revio instruments.

**Library preparation details - PacBio**

*[GCC-BCM]:* Genomic DNA was quantified using Qubit dsDNA quantification broad range assay (Thermo Fisher Scientific). DNA size was determined using the Femto Pulse System (Agilent). A total of 2 libraries were prepared for each sample to be sequenced across 3 SMRT Cells to achieve 90× coverage. DNA was sheared using Covaris g-tubes (Covaris 520079) to achieve an average size of 18-22 kbp. Sheared DNA was size-selected on the PippinHT instrument (Sage Science) using the 6-10 kbp or the 15-20 kbp High-Pass definition. The size selected DNA was used as input for Pacbio SMRTBell Prep Kit 3.0 for library preparation. DNA damage repair, A-tailing and adapter ligation were performed as per manufacturer's instructions. SMRTBell Adapter Index Plate 96A was used for barcoding each library. Adapter ligated DNA was nuclease treated following the manufacturer's guidelines and purified using 1X Pacbio SMRTBell Cleanup beads. Final libraries were eluted in 30 µL Pacbio elution buffer and quantified using the Qubit dsDNA quantification high-sensitivity assay (Thermo Fisher Scientific). Final library size was determined using the Agilent Femto Pulse.Sequencing primer annealing and polymerase binding were performed using the Revio Binding Kit 3.0. Libraries were loaded onto the PacBio Revio machine utilizing SMRTlink v13 for workflow setup and loaded at 325pM loading concentration with 30 hours of movie time.

*[GCC-UW-SCRI]:* Extracted DNA samples were checked for quantity using Qubit dsDNA HS (Thermo Fisher, Q32854) measured on DS-11 FX (Denovix) and size distribution using FEMTO Pulse (Agilent, M5330AA & FP-1002-0275). Depending on initial length distribution, samples were sheared to a target peak length of ~20 kbp. Some samples were left unsheared; others were sheared with Megaruptor 3 Hydropores (Diagenode, B06010003 and E07010003). Moderately degraded or low-concentration samples were subjected to a light shear at setting 28 or 29, while intact DNA samples were sheared by processing twice, at settings 28 or 29 and 30 or 31. Highly intact and nonhomogeneous DNA samples (e.g., derived from fibroblasts) were pre-sheared with Megaruptor 3 DNAFluid+ (Diagenode, E07020001) before final shearing. Sheared DNAs were subjected to PacBio HiFi library prep via the SMRTbell Prep Kit 3.0 (PacBio, 102-182-700) using barcoded adapters (PacBio, 102-009-200). When final library QC permitted (minimum of 500 ng of individual or pooled libraries,) size selection was performed

with Pippin HT using a high-pass cutoff of 8-15 kbp (Sage Science, HTP0001 & HPE7510). Low-mass libraries or those with short final size distributions (~5-10 kbp) were instead treated with a mild size-selection using diluted AMPure PB beads per the manufacturer's protocol. Libraries were sequenced on the Revio platform on SMRT Cells 25M with Revio Chemistry V1 (PacBio, 102-817-900) or SPRQ (PacBio, 103-520-200) with Adaptive Loading and 30-hour movies.

*[GCC-Broad]:* Long-read whole-genome sequencing libraries were prepared from high-molecular-weight (HMW) genomic DNA for sequencing on the PacBio Revio platform. HMW DNA was quantified using PicoGreen fluorometry and diluted to the target input mass (approximately 3 µg at 58 ng/µL for standard-input samples), with fragment size assessed by Agilent Femto Pulse capillary electrophoresis to confirm input quality. Short DNA fragments (<10 kbp) were selectively depleted using a Short Read Eliminator (SRE) kit (1 hour at 50°C followed by centrifugation at 3,220 × g and 29°C for 1 hour), after which the remaining HMW DNA was sheared to a target size of approximately 15 kbp by controlled pipette shearing on a Hamilton STARlet liquid handler. Fragment size was verified by Femto Pulse using a 165 kbp ladder to confirm a 10-20 kbp size distribution. Sheared DNA was concentrated using SMRTbell bead cleanup and subjected to DNA damage repair and A-tailing (30 minutes at 65°C followed by 5 minutes at 37°C), after which barcoded SMRTbell adapters were ligated using the PacBio HiFi Prep Kit (30 minutes at room temperature) to generate circular SMRTbell libraries. Unligated or incomplete constructs were removed by nuclease digestion (15 minutes at 37°C), and a final diluted AMPure bead size-selection step enriched libraries >5 kbp, which were eluted in 25 µL and stored at −20°C until sequencing.
Prior to sequencing, SMRTbell libraries underwent primer annealing (15 minutes at room temperature) and polymerase binding using the Revio SPRQ Polymerase Kit (15 minutes at room temperature), followed by SMRTbell bead cleanup to remove excess polymerase. Library concentration was measured by Qubit fluorometry, and PacBio SMRT Link Loading Calculator software was used together with the measured library concentration and Femto Pulse insert-size distribution to determine the final loading concentration (typically 200–300 pM for standard libraries). The final diluted library, combined with a serially diluted internal control complex, was loaded onto PacBio Revio SMRT Cell 8M trays for sequencing. Sequencing was performed to a target depth of 12×–20× per sample, producing HiFi reads with an average length of 14.8 kbp. Raw sequence data were processed using PacBio circular consensus sequencing (CCS) to generate highly accurate HiFi consensus reads.

*[GCC-WashU/VAI]:* PacBio HiFi SMRTbell libraries were prepared following PacBio protocol 'Procedure & Checklist – Preparing Whole Genome and Metagenome Libraries Using SMRTbell Prep Kit 3.0'. Genomic DNA was fragmented with a mode of ~20 kbp using the Diagenode Megaruptor 3 instrument. Genomic DNA was then sheared twice using Shearing kit (P/N E07010003) with speeds of 28 and 30. Sheared sample was assessed via fluorometry (Qubit High Sensitivity DNA Kit) and Agilent Femto Pulse (Genomic DNA 165kb Kit). Libraries were made according to PacBio protocol utilizing barcoded adapters from SMRTbell adapter index plate 96A (PacBio P/N 102-009-200) to allow for multiplexing of samples during sequencing. Libraries were size selected using Sage PippinHT instrument and the 0.75% Agarose High-Pass 75E kit (P/N HPE7510) with a start size of 10000bp-15000bp. Size selected libraries were prepared for sequencing following instructions generated in PacBio SMRT Link v25.1 Sample Setup and utilizing PacBio Revio SPRQ polymerase kit (P/N 103-520-100). Sequencing was performed on PacBio Revio sequencer with an 'On Plate Concentration' of 170pM-200pM. Two SMRTcells per sample were targeted for sequencing for a total of ~60X and ~186Gb total coverage.

**Library preparation details - ONT**

*[GCC-BCM]:* Genomic DNA was quantified using Qubit dsDNA quantification broad range assay (Thermo Fisher Scientific). DNA size was determined using the Femto Pulse System (Agilent). Libraries were prepared from ~9 µg genomic DNA to be sequenced on 2 flow cells with the aim of achieving 60× coverage. DNA was sheared using Covaris g-tubes (Covaris 520079) to achieve an average size of 15-20 kbp for DNA from tissues. Sheared DNA was size-selected on the PippinHT instrument (Sage Science) using the 6-10 kbp High-Pass definition. ONT libraries were prepared using the SQK-LSK114 kit and the NEBNext Companion Module (E7180) following the manufacturer's instructions. Final libraries were eluted in the ONT elution buffer and quantified using the Qubit dsDNA quantification broad range assay. R10.4.1 flow cells were loaded with 15 fmoles of the final library and sequenced for 72 hours. If needed, the flow cell was washed and reloaded with an additional 15 fmoles of the library. A low-input protocol was used for samples that lacked sufficient starting material for standard library preparation. Low-input libraries were prepared from 1.5 - 2 µg genomic DNA that was sheared using g-tubes to achieve an average size of 12-18 kbp. ONT libraries were prepared using the SQK-LSK114 kit and the NEBNext Companion Module v2 (E7672) following the manufacturer's instructions. Low-input libraries were loaded on R10.4.1 flowcells using a loading amount of 7-10 fmoles. Reloads were performed as needed during a 72-hour sequencing run.

*[GCC-NYGC]:* High molecular weight (HMW) DNA samples were first sheared using the Megaruptor 3 (Hologic Diagenode, catalog number B06010003) to a target fragment size of 45 kbp, following the manufacturer's recommendations. Fragmented DNA quality was assessed using NanoDrop (ND-2000), Qubit Broad Range Assay (Q32850), and the Genomic TapeStation (G2964AA), all according to manufacturer instructions.
To improve 260/230 ratios, a 3X buffer exchange clean-up was performed using AMPure XP Beads for DNA Cleanup (A63882). The cleaned, fragmented HMW DNA was then prepared using the ONT Ligation Sequencing Kit V14 (SQK-LSK114), following the manufacturer's protocol. Final libraries were quantified again using the Qubit Broad Range Assay (Q32850) and Genomic TapeStation (G2964AA). HMW libraries were sequenced on Promethion P24 (PRO-SEQ024) using R10.4.1 flow cells (FLO-PRO114M) according to manufacture instructions.

*[GCC-UW-SCRI]:* DNA was extracted from tissue samples using the NEB Monarch HMW DNA extraction kit for Tissues (#3060L). No more than 25mg of tissue was cut into small chunks and then placed into a 1.5mL tube and mashed with the pestle. The sample was lysed according to the protocol at a shaking speed of 1000rpm (liver at 2000rpm; colon at 2000rpm for 15min, no shaking 30min). After lysis and RNAse treatment, brain samples were put on ice for 3 min before protein precipitation and phase separation. 800uL of the upper phase is combined with isopropanol to precipitate the DNA. DNA washes and elution followed the manufacturer's protocol. DNA was quantified by Qubit and size distribution was measured on the Femto Pulse. In order to selectively remove smaller fragments of DNA, we performed a Short Read Eliminator (PacBio 102-208-400) and cleaned up the DNA with a bead wash.

Libraries were constructed using between 2-9ug of DNA and the Ligation Sequencing Kit from ONT (SQK-LSK114) with modifications to the manufacturer's protocol. End repair was incubated for 20 minutes and the adapter ligation was incubated for 1 hour. The final library was eluted in 30ul of EB and quantified by Qubit. 200-400ng of library was loaded onto a FLO-PRO114M R10.4.1 flow cell for sequencing on the PromethION, with two nuclease washes and reloads after 24 and 48 hours of sequencing.

**S2.4 Duplex sequencing**
CODEC libraries were sequenced on Illumina NovaSeq X Plus. Detailed protocols for CODEC, NanoSeq, CompDuplex-seq and META-VISTA-seq, including adapter, barcoding and transposase strategies, input requirements and per-method duplex yield, are given below.

**Duplex sequencing: Library preparation, sequencing and analysis details**
*[GCC-Broad]:* CODEC libraries were prepared as previously described[6], using genomic DNA obtained from the same AllPrep DNA/RNA Universal Kit extraction described above. Genomic DNA was enzymatically fragmented to an average size of ~150 bp, and 20 ng of fragmented DNA was dA-tailed with a mixture of dATP and ddNTPs (ddTTP, ddCTP, ddGTP) following the method of Abascal et al. The dA-tailed DNA was used for CODEC library construction with CODEC quadruplex adapters, PCR amplified, and sequenced on a NovaSeq X Plus 25B flow cell, targeting approximately 0.5-1x duplex coverage.
CODEC data were processed as previously described (Bae et al.), except that the hg38 reference genome without decoy sequences was used in this study, with samples analyzed from BCL files to variant calls using the standard CODECsuite (https://github.com/broadinstitute/CODECsuite).

*[GCC-BCM]:* CompDuplex libraries were prepared as previously described (https://www.protocols.io/view/compduplex-accurate-detection-of-somatic-mutations-kxygx3x4og8j/v1). For genomic DNA extracted from production donor tissues, 20 ng extracted genomic DNA was sealed with thiol-ddNTP in the following reaction mix: 1 uL of 10X ThermolPol Reaction Buffer (New England BioLabs, Cat. B9004S), 0.2 uL of 5mM each thoil-ddNTP (TriLink BioTechnologies, Cat. K1003), 0.15 uL of *Bst* DNA Polymerase, Large Fragment (New England BioLabs, Cat. M0275L), nucleus free water to 10 uL. The reaction was incubated at 65℃ for 10 min, quenched with 1 uL 0.5 M EDTA. Next, the sealed genomic DNA was purified with 0.5X Ampure XP beads for CompDuplex library preparation. For each sample, we aimed at the library complexity of ~90 million genomic DNA fragments, and the libraries were sequenced on the Illumina NovaSeq X 25B flow cell at 5~8 reads per fragment. Analysis details are described here[7].

*[GCC-BCM]:* NanoSeq libraries. NanoSeq (HpyCH4V and Mung Bean) libraries for the donors materials were prepared using 50-100ng of DNA as described by Abascal et al. with minor changes as described in Chao et al. https://www.protocols.io/view/optimized-mung-bean-nuclease-nanoseq-libraryprepa-
d4h38t8n.html). Libraries were pooled and sequenced on Illumina NovaSeq sequencing platforms for 30× coverage. Analysis details are described here[8].

META-VISTA-seq libraries were prepared from 250pg of bulk tissue genomic DNA as described previously https://www.protocols.io/view/tn5-duplex-sequencing-tn5-duplex-seq-for-low-input-6qpvr3nbzvmk/v1. The method, adapted from META-CS, uses engineered Tn5 transposase complexes pre-loaded with strand-specific adapters to simultaneously fragment and tag both strands of each DNA duplex[9]. Libraries were sequenced on the Illumina NovaSeq platform with paired-end 150bp reads.

**S2.5 PTA single-cell DNA sequencing**
Single-cell DNA sequencing was performed on 91 cells from the liver (LIVR), testis (TESL), heart left ventricle (HART), sun-exposed skin (SKSE) and brain (cerebellum, BRCE; hippocampus, BRHL; and frontal lobe, BRFL) of donor SMHT005. Single cells or nuclei were isolated from tissue cores using custom protocols for each tissue type.

**Nuclei isolation, Library prep, and sequencing - PTA**
*[TTD-Walsh]:* Single nuclei were isolated from SMHT005 NeuN-/NeuN+ cerebellum (BRCE), hippocampus (BRHL) and sun-exposed skin (SKSE) and amplified for DNA sequencing using the following protocols. 10 cerebellar nuclei (5 NeuN- and 5 NeuN+) and 10 hippocampal nuclei (5 NeuN- and 5 NeuN+) were collected using fluorescence-activated nuclei sorting (FANS) with nuclear staining for NeuN (Millipore, MAB377; clone A60, 1:1,500) and DAPI following a published protocol (https://www.protocols.io/view/ultra-deep-atac-seq-for-sorted-neurons-eq2lyw35rvx9/v1). 20 nuclei from sun-exposed skin were isolated using the same FANS protocol but excluded the NeuN staining step. All nuclei were sorted into a 96-well plate containing BioSkryb PTA v1 kit cell buffer. Sorted nuclei were subjected to genome amplification using the Primary Template-directed Amplification v1 kit (BioSkryb, 100136) following the manufacturer's protocol with the following adjustments. The initial lysis step was adjusted to plate mixing at room temperature for 1 minute then transferring the plate to incubate on ice (or PCR cooler) for 20 mins. The rest of the protocol was completed following the manufacturer's protocol. PTA product then underwent two QC tests: DNA concentration was quantified using Thermofisher Qubit Flex and DNA size, quantity, and integrity were tested using the HS D5000 TapeStation screen tape (Agilent, 5067-5593). A 4-locus PCR test was performed[10]. Sequencing libraries were constructed using the KAPA HyperPrep Kit (Roche, 07962363001) and the KAPA Unique Dual-Indexed Adapter Kit 15µM (Roche, 08861919702) following the manufacturer's protocol with a custom size selection step. In place of the recommended KAPA HyperPrep kit size selection, the size selection protocol from the Primary Template-directed Amplification kit (BioSkryb, 100136) was followed and repeated a second time. Five library pools containing 8 libraries each were sent to Broad for sequencing. Each library pool was sequenced on a single NVX 25B flowcell following the short-read Illumina protocol except that the library preparation step was omitted.

*[TTD-Choudhury]:* Approximately 100 mg of left-ventricular (HART) tissue from donor SMHT005 was finely minced and gently homogenized by Dounce homogenization and pipetting in 5 mL of ice-cold isolation buffer containing 0.32 M sucrose, 5 mM $CaCl_2$, 3 mM magnesium acetate, 2 mM EDTA, 0.5 mM EGTA, 10 mM Tris-HCl (pH 8.0), and 1 mM dithiothreitol. Cardiomyocyte cells were isolated by filtering the homogenate through 100 and 70 µM strainers, respectively (Pluriselect), and the resulting pellet was resuspended in a 0.1% PBS BSA. Cells were then incubated with cardiac troponin T conjugated with 488 fluorophores (dilution=1:100) for 20 minutes. After 20-minute incubations, cells were spun down at 750g for 10 minutes. The resulting cell pellet was resuspended in 1 mL of 0.1% BSA in PBS for sorting. For endothelial nuclei isolation, the homogenate was sequentially filtered through 100- and 70-µm strainers, and cardiac nuclei were purified by sucrose-density-gradient centrifugation. The nuclear pellet was resuspended in PBS containing 0.1% bovine serum albumin, passed through a 40-µm strainer, and counted. Purified nuclei were incubated for 20 min Alexa Fluor 647-conjugated anti-ERG antibody (Abcam, clone EPR3864) to identify the endothelial fraction. After staining, nuclei were centrifuged at 750 × g for 10 min, washed, and resuspended in 1 mL of PBS containing 0.1% bovine serum albumin. DAPI was added immediately before fluorescence-activated nuclei sorting. Following exclusion of debris, aggregates, and DAPI-negative events, single $DAPI^+$/cardiac troponin $T^+$ cardiomyocyte nuclei and $DAPI^+/ERG^+$ endothelial nuclei were collected separately.

Cardiomyocyte cells and endothelial nuclei were sorted into 96-well plates and subjected to whole-genome amplification by primary template-directed amplification (PTA). PTA was performed using the ResolveDNA Whole Genome Amplification Kit. Cells were sorted into 3 µL of pre-chilled Cell Buffer, lysed by adding 3 µL of MS Mix, and neutralized with 3 µL of SN1 buffer. Next, 3 µL of SDX reagent was added, and the samples were incubated at room temperature for 10 minutes. An 8 µL enzyme-containing reaction mixture was then added to obtain a final reaction volume of 20 µL. Amplification proceeded at 30°C for 10 hours, followed by enzyme inactivation at 65°C for 3 minutes. The amplified DNA was purified using AMPure beads, and DNA yield was quantified by PicoGreen binding with the Quant-iT dsDNA Assay Kit (Thermo Fisher Scientific). Amplified genomes were assessed by multiplex PCR and Bioanalyzer analysis. Samples that showed successful amplification at all four multiplex PCR loci were subsequently prepared for Illumina sequencing after library preparation. Libraries were prepared and sequenced at WashU/VAI following the short-read Illumina protocol except that libraries were constructed using the KAPA Hyper Prep Kit (Roche, Cat # 7962363001).

*[TTD-Urban]:* Single nuclei from the liver (LIVR), testes (TESL) and brain frontal lobe (BRFL) of donor SMHT005 were isolated and subjected to whole-genome amplification. 10 mg of tissue was chopped it into small pieces and resuspended in lysis buffer (319.6 mM sucrose, 5 mM $CaCl_2$, 3 mM $Mg(Ac)_2$, 0.1 mM EDTA, 10 mM Tris-Cl pH 7.4, 1 mM DTT, 0.1% Triton X-100). Samples were dounced in the the buffer and the suspension was passed the suspension through a 40 µm filter, and centrifuged at 500 rpm for 10 minutes at 4 °C. The pellet was resuspended in a sorting buffer (1% BSA, 1 mM EDTA, 10 mM HEPES in 1× PBS), stained with PI (10ug/ml) and kept on ice until sorting. Nuclei were sorted using a 100 µm nozzle and positive for PE-Texas red channel. FSC-A/SSC-A gating was used to assess nuclear integrity and FSC-A/FSC-H gating was used to exclude doublets, and dispensed into BioSkryb cell buffer 1 cell/well.

Whole-genome amplification followed the ResolveDNA™ PTA kit V1 (BioSkryb) protocol. All amplification steps were done in a DNA-free pre-PCR hood using either ResolveDNA™ PTA kit V1 (BioSkryb) per the manufacturer's instructions. Briefly, sequential addition of MS mix (1.5 µL SM2 + 1.5 µL 1× SS2), SN1 and SDX with mixing intermittently at 1,400 rpm was carried out to lysis nuclei and deproteinize DNA. For amplification, an 8 µL reaction mix (SB4, SEZ1, SEZ2, 1× SS2) was added and incubated at 30 °C for 10 hours, then 65 °C for 3 minutes to stop the reaction, followed by a 4 °C hold. Amplified DNA was purified with ResolveDNA magnetic beads and DNA concentration was measured using the Qubit™ dsDNA High Sensitivity Assay Kit (Thermo Fisher Scientific, Cat. Nos. Q32851/Q32854). A 4-locus PCR test was performed[11]. Library preparation and sequencing was performed at BCM following the short-read Illumina protocol.

**S2.6 Bulk and single-nucleus RNA-seq**

Two types of bulk RNA-seq data were generated from donor tissues: short-read whole-transcriptome sequencing (total RNA-seq) on Illumina using the Watchmaker RNA Library Prep Kit, and full-length transcript data on the PacBio platform using the Kinnex full-length RNA library protocol. Ribosomal and globin depletion was applied to the total RNA-seq libraries. Single-nucleus RNA-seq and ATAC-seq libraries were generated on the 10x Genomics platform and sequenced on Illumina.

**Library preparation details - Bulk tissue RNA-seq**

*[GCC-BCM]:* Whole transcriptome sequencing (total RNA-seq) data was generated using the Watchmaker RNA Library Prep Kit with PolarisR Depletion (7BK0002-096, Watchmaker Genomics). RNA quality and quantity was estimated using Agilent Bioanalyzer. To monitor sample and process consistency, 1 µl of the 1:50 diluted synthetic RNA designed by External RNA Controls Consortium (ERCC) (4456740, ThermoFisher) was added to 1 µg total RNA. In addition, as a process control, the Universal Human Reference RNA (UHR) (740000, Agilent Inc.), was processed in parallel with the RNA samples. Libraries were sequenced on the

NovaSeq 6000 instrument using the S4 reagent kit (300 cycles) to generate 2x150bp paired-end reads. In order to generate a minimum of 157-300M read-pairs per sample.
Mapping and QC Pipeline for Internal Quality Assessment Before Data Submission to DAC
The RNA-seq analysis pipeline cleans and processes raw RNA-seq data (FASTQs), providing robust QC metrics and has the flexibility to map the reads to either GRCh37 reference or GRCh38 (after excluding the alternate contigs). The pipeline aligns RNA-seq reads, removes duplicates, and generates QC metrics. It also quantifies gene expression using RSEM, generates QC metrics, and produces raw gene feature counts.

*[GCC-NYGC]:* Total RNA libraries were prepared with Watchmaker's RNA Library Prep kit with Polaris Depletion (catalog number 7BK0002), according to manufacturer recommendations. 200ng of total RNA were used for input for RNA purification and fragmentation. Purified RNA underwent first and second strand cDNA synthesis. cDNA was then adenylated, ligated to Illumina sequencing adapters, and amplified by PCR (using 10 cycles). cDNA libraries were quantified using Fragment Analyzer (Agilent) and Spectramax M2 (Molecular Devices). Bulk RNA libraries were sequenced on the Illumina NovaSeq X Plus, using 2 x 150 bp cycles.

*[GCC-Broad]:* Sequencing libraries were prepared from total RNA using the Watchmaker Total RNA Library Construction Kit with Polaris ribosomal/globin RNA depletion. Input RNA was quantified using a RiboGreen assay and normalized to approximately 500 ng total input (28 ng/μL). Ribosomal (cytoplasmic 28S/18S/5.8S/5S, mitochondrial 16S/12S, and 45S precursor) and globin (HBA1, HBA2, HBB, HBD, HBM, HBG1, HBG2, HBE1, HBQ1, and HBZ) transcripts were depleted by hybridization to sequence-specific probes (77°C denaturation followed by 65°C digestion for 15 minutes) and enzymatic digestion, followed by degradation of unbound probes (37°C for 10 minutes), an FFPE de-crosslinking step (70°C for 30 minutes), and AMPure XP bead cleanup. The depleted RNA was fragmented by heat and divalent cations (65°C for 1 minute), reverse transcribed into first-strand cDNA (25°C priming, 42°C reverse transcription, and 70°C enzyme inactivation), converted to double-stranded cDNA, and A-tailed (62°C for 10 minutes). Unique molecular identifier (UMI)-containing adapters were ligated to the A-tailed cDNA (20°C for 15 minutes), followed by SPRI bead cleanup. Libraries were PCR amplified using dual-indexed (P5/P7) barcoded primers (98°C initial denaturation; 10 cycles of 98°C, 55°C, and 72°C; followed by a final extension at 72°C) and purified by a second SPRI bead cleanup.

Completed libraries were quantified by PicoGreen fluorometry, normalized, and pooled. Prior to sequencing, pooled libraries were quantified by qPCR using a SYBR-based standard-curve assay on a QuantStudio 7 instrument, normalized to a target concentration of 2 nM, pooled at 94-plex, and re-quantified by qPCR. Pools falling outside the acceptable loading range (1–3 nM) were renormalized to 2.5 nM before sequencing on the Illumina NovaSeq X Plus 25B platform using 2 × 146 bp paired-end reads to a target depth of 200 million read pairs per sample. Sequencing runs were required to meet predefined quality-control metrics, including the minimum number of reads passing filter for the 25B flow cell configuration (up to 22 billion reads across eight lanes), ≥80% of bases with a quality score of Q30 or higher, and ≥70% of clusters passing filter and occupied. Reads were aligned to the GRCh38 reference genome using Illumina Dynamic Read Analysis for GENomics (DRAGEN; version 4.2.4), and data were delivered as de-multiplexed, aligned CRAM files with accompanying CRAI and MD5 checksum files.

*[GCC-WashU-VAI]:* Total RNA integrity was determined using Agilent Bioanalyzer or 4200 Tapestation. Library preparation was performed with 100ng to 500ng of total RNA. Libraries were generated with Watchmaker Library Prep Kit with Polaris Depletion (Watchmaker). Briefly,

ribosomal RNA was removed by an RNaseH method and purified with RNAClean beads (Beckman). mRNA was then fragmented in buffer by heating depending on RNA quality per protocol. mRNA was reverse transcribed to yield strand-specific cDNA. A second strand and A Tailing reaction was performed to yield fragments with an A base added to the 3' ends. Illumina sequencing adapters were ligated to the ends. Ligated fragments were then amplified per protocol using primers incorporating unique dual index tags. Fragments were sequenced on an Illumina NovaSeq X Plus using paired end reads extending 150 bases.

**PacBio Kinnex full-length transcript sequencing**
*[GCC-UW-SCRI]:* Extracted total RNA was quality checked using UV-Vis spectroscopy (Denovix DS-11 FX) and Agilent Bioanalyzer 2100 using the Total RNA Nano 6000 kit (Agilent, G2939A & 5067-1511.) Kinnex full-length RNA libraries were generated per manufacturer's recommendations (PacBio, 103-072-000). Samples were sequenced on the Revio platform on SMRT Cells 25M with Revio Chemistry V1 (PacBio, 102-817-900) or SPRQ (PacBio, 103-520-200) with Adaptive Loading and 30-hour movies. Data were postprocessed using SMRT Link v13.1 or 13.3 with the "Read Segmentation and Iso-Seq" pipeline to segment and classify reads.

**Library preparation details - snRNA and ATAC-seq**
*[GCC-BCM]:* Samples from 12 of the 25 donors were used for single-nucleus profiling, with 58 tissue samples processed for snRNA-seq and 42 tissue samples for snATAC-seq. Frozen tissue samples were minced on dry ice and lysed in Nuclei Extraction Buffer (Miltenyi Biotec) using a gentleMACS Dissociator (Miltenyi Biotec). Throughout the nuclei isolation procedure, all buffers were supplemented with RNase inhibitor to preserve RNA integrity. The resulting nuclei suspension was sequentially filtered, pelleted by centrifugation, gently triturated, and purified by Anti-Nucleus MicroBeads (Miltenyi Biotec) through positive magnetic selection to enrich intact nuclei and remove cellular debris. Purified nuclei were washed, resuspended, counted, and assessed for nuclear integrity and morphology prior to downstream library preparation. Purified nuclei were loaded onto a Chromium X instrument (10x Genomics) for GEM generation and library preparation according to the manufacturer's instructions. Single-nucleus RNA-seq libraries were prepared using the GEM-X Universal 5′ Gene Expression v3 kit (10x Genomics), whereas single-nucleus ATAC-seq libraries were prepared using the Chromium Next GEM Single Cell ATAC Reagent Kits v2 (10x Genomics). Libraries were sequenced on an Illumina NovaSeq 6000 platform (Illumina). Gene expression libraries were processed using Cell Ranger v9.0.1 (10x Genomics), and ATAC-seq libraries were processed using Cell Ranger ATAC v2.2.0 (10x Genomics) using the SMaHT reference package based on the GRCh38 human reference genome and GENCODE release v47 annotation.

**Single-cell Total RNA-seq (STORM-seq)**
*[GCC-WashU-VAI]:* Single cell suspensions from the benchmarking cell line mixtures were rapidly thawed in a 37C water bath and resuspended in increasing 1:1 dropwise volumes of warm (37C) flow buffer (HBSS with no divalent cations + 2% FBS + 25 mM HEPES) with gentle agitation. Resuspended cells were then pelleted at 300 x g at 4C for 5 minutes. Cells were washed in warm flow buffer and re-pelleted. Next, cells were counted and resuspended at 1 x 10^6 cells/mL in warm flow buffer containing 0.5 ug/mL DAPI for active viability surveillance during sorting. Single, live cells were index sorted into each well of a 384-well plate (Eppendorf) containing 2.17 uL Fragmentation Buffer (1.17 uL PBS pH 7.2 [Gibco 20012-027], 0.17 uL 10X Lysis Mix, 0.17 uL SMART scN6, 0.69 uL scRT buffer) using a BD FACSymphony S6 sorter running BD FACSDiva v9.1.3 and equipped with BD StepSort. The S6 was run with a 130 um nozzle at 14 psi. For cell deposition into 384-well plates, we used single cell sort mode. Single cell libraries were prepared using the STORM-seq kit (Takara Cat # 634751) and protocol (https://doi.org/10.5281/zenodo.15178455). Briefly, after sorting, plates were immediately

transferred to a pre-heated thermal cycler, heated to 85C for 3 minutes and snap cooled on ice for 2 minutes. Once cooled, 1.17 uL First Strand Master Mix (0.75 uL SMART scTSO mix, 0.08 uL RNase Inhibitor, 0.34 uL SMARTscribe RT) is added to each well. Immediately prior to dispensing, ERCC RNA Mix 1 (Thermo Fisher) is added to the First Strand Master Mix to create a 1:1000000 final dilution. First Strand Synthesis is performed at 42C for 180 minutes, followed by 70C for 10 minutes and 4C hold. After 1st Strand Synthesis, PCR1 (10 cycles of amplification: 94C 1 min, [98C 15s, 55C 15s, 68C 30s]x10, 68C 2 min, 4C hold) was performed with the addition of SMARTer RNA Unique Dual Index Sets A-D (SMARTer RNA Unique Dual Index Kits 96U sets A-D, Takara Cat # 634752, 634753, 634754, 634755). 4.67 uL of PCR1 Master Mix (0.33 uL nuclease free water, 4.16 uL SeqAmp CB PCR buffer, SeqAmp DNA Polymerase) is added to each well. Each well then receives 1 uL of a Unique Dual Index (UDI). UDIs were diluted 1:4 in 10mM Tris-HCl, pH 8.0 (Teknova) prior to addition. The plate was pooled and a bead-based cleanup (Beckman Coulter AMPure XP Beads) was performed. 162 uL of rRNA Depletion Master Mix (123.12ul nuclease free water, 16.2 uL 10X ZapR buffer, 11.02ul scZapR, 11.02ul sc-R Probes) is used to elute the depleted cDNA from the dried beads. The eluate is incubated at 37C for 60 min, 72C for 10 min, 4C hold. Finally, 12 cycles of amplification (94C 1 min, [98C 15s, 55C 15s, 68C 30s]x12, 4C hold) were performed for PCR2 (PCR2 Master Mix (208 uL nuclease free water, 400 uL SeqAmp CB PCR Buffer, 16ul PCR2 primers, 16 uL SeqAmp DNA polymerase) and the final library was eluted in 20 uL 10mM Tris-HCl pH 8.0. After QC, where necessary, an additional bead clean-up was performed to remove any remaining adapter-dimer. For the STORM-seq libraries, sizing was performed using the Agilent Bioanalyzer HS kit and concentration was determined using a Qubit fluorometer and dsDNA HS kit before sequencing using an Illumina NovaSeq 6000 S2 2x150 bp flow cell. Libraries were sequenced to generate an average of 1M reads per cell.

**S2.7 MEI capture method: Library preparation and sequencing**
*[TTD-Mills]:* Transposable Element Nanopore Cas9-Targeted Sequencing (TEnCATS) libraries were prepared from SMHT005 tissue cores as described by McDonald et al.[12] (https://dx.doi.org/10.17504/protocols.io.kqdg3q66ev25/v2) with the modifications detailed below. High molecular weight (HMW) genomic DNA (gDNA) was extracted using the Monarch HMW DNA Extraction Kit for Tissues (NEB, T3060L). For each library, 10–20 µg of genomic DNA was dephosphorylated with Quick CIP (NEB, M0525S). Cas9 ribonucleoprotein (RNP) complexes were assembled using Alt-R HiFi Sp. Cas9 (IDT, 1081060) and 850 ng of sgRNA and incubated at room temperature for 20 min. For BRCE, ESOP, COAS, CODS, LUNG, and TESL, separate RNP preparations targeted either L1Hs or AluYa5 and AluYb8 together. For SKSE, SKNE, BRTL, MUSC, HART, and LIVR, a single RNP preparation combined sgRNAs targeting L1Hs, *Alu*Ya5, and *Alu*Yb8. Dephosphorylated DNA was simultaneously dA-tailed with Taq DNA Polymerase (NEB, M0273S) and cleaved by the Cas9 RNPs. Thermolabile Proteinase K (5 µL; NEB, P8111S) was then added to degrade the Cas9 protein. The dA-tailed, TE-enriched DNA was used for library construction following the ONT ligation sequencing protocol with T4 DNA Ligase (NEB, M0202M) and the ligation sequencing kit (SQK-LSK114). Libraries were sequenced for 72h on a PromethION2 Solo using three PromethION flow cells per tissue sample. Reads were quality-controlled and filtered using samtools and Minimera, then aligned to GRCh38 using minimap2. The expected and reported sequence coverage is based on the median coverage across the on-target MEI loci of interest. sMEIs were identified using Nanopal, compared among tissues, and manually inspected.

**S2.8 DSA methods**
To generate DSA for donors, fibroblasts were cultured and DNA extracted. We used a combination of PacBio HiFi sequencing (or Fiber-seq) data, UL-ONT WGS data, and Hi-C to construct DSA. While PacBio HiFi methods are described earlier, others are discussed below.

*[GCC-UW-SCRI] Fibroblast culturing:* Cryopreserved fibroblasts cultures were thawed and expanded in a 1:1 mixture of M199 Medium (Invitrogen, 11150-059) and M106 Medium (Invitrogen, M-106-500) supplemented with 15% FBS (ATCC, 10010-023), 0.4ug/ml Hydrocortisone (Sigma, H0888-1G), 10ng/ml EGF (Invitrogen, PHG0311), 0.25ug/ml Fungizone (Invitrogen, 15290-018), 50ug/ml Gentamicin (Invitrogen, 15750-060), and 100U/mL Penicillin-Streptomycin (Invitrogen, 15140-122). Adherent cell cultures were split using 0.25% Trypsin-EDTA (Invitrogen Cat. No. 25200-114).

*[GCC-UW-SCRI] UL-ONT extraction and sequencing:* Ultra-high Molecular Weight DNA was extracted from fibroblast cell lines using a phenol chloroform extraction protocol (Logsdon, protocols.io, 2020). Briefly, 2-3 x $10^7$ cells were lysed in a buffer containing 10 mM Tris-Cl (pH 8.0), 0.1 M EDTA (pH 8.0), 0.5% w/v SDS, and 20mg/mL RNase A (Qiagen, 19101) for 1 hour at 37°C. 200 ug/mL Proteinase K (Qiagen, 19131) was added, and the solution was incubated at 50°C for 2 hours. DNA was purified via two rounds of 25:24:1 phenol-chloroform-isoamyl alcohol extraction followed by a chloroform back extraction. DNA was precipitated with ethanol and was solubilized in EEB (ONT) at 4°C for two days.

Alternatively, DNA was extracted from cell lines using the NEB Monarch HMW DNA extraction kit for Cells & Blood (#T3050L) following the manufacturers protocol with the following exceptions. 6 million cells were used for the starting input with a shaking speed of 600 rpm during the lysis step. DNA was precipitated with 300uL EEB (ONT) and solubilized at 4°C for two days.

Libraries were constructed using the Ultra-Long DNA Sequencing Kit V14 (SQK-ULK114) following the manufacturer's protocol. For monarch DNA, two extractions were combined for library prep and final elution volume was doubled from the protocol. For phenol chloroform extracted DNA, approximately 40ug of DNA was input into library prep and the final elution volume ranged from 2x-4x the protocol volume, depending on DNA visualized during the clean-up step. Final libraries were left at room temp over night to solubilize. 75 uL of library was loaded onto a primed FLO-PRO114M R10.4.1 flow cell for sequencing on the PromethION, with two nuclease washes and reloads after 24 and 48 hours of sequencing.

*[GCC-UW-SCRI] Fibroblast Hi-C reaction and sequencing:* Cultured fibroblasts were harvested and counted. 1-2M cells were washed and frozen dry at -80°C. Omni-C library preparation was conducted using either the Dovetail Omni-C Kit (PN 21005G Dovetail® Omni-C® Kit) for human cells v2.0 or Dovetail Omni-C Kit for human cells v2.1 (updated July 2025), or Dovetail Omni-C Kit for human cells v3 (updated December 2025), with minor modifications. Cell pellets were treated with 0.5-2uL of the Omni-C nuclease mix. Sample digests were reviewed and compared against the recommended range in the Omni-C protocol prior to proceeding with proximity ligation. 150-200ng of proximity-ligated sample was used for library preparation. In cases where <150ng was available, all samples were used for the library prep. The number of PCR cycles used for library preparation was dependent upon the amount of starting material from proximity ligation and ranged between 9-12 cycles. The final library was sequenced on an Illumina NovaSeqX platform and generated at least 1.2 billion 2×150 read pairs.

### Supplementary Methods Section 3 | Genome alignment, reference assemblies, and QC

#### S3.1 Reads alignment to the GRCh38 genome

*Illumina WGS:* Whole-genome sequencing reads were processed using the same alignment pipeline established for the SMaHT benchmarking study[13]. In brief, paired-end Illumina reads were first analyzed using fastp (v0.23.2) to remove reads containing artifactual poly-G runs from two-color Illumina NovaSeq X Plus chemistry, then aligned to the GRCh38 reference genome (GCA_000001405.15, excluding ALT contigs and hs38d1 decoy sequences) using BWA-MEM (v0.7.17) as implemented in Sentieon (v202308.01); this ALT/decoy-free build was used because ALT contigs can lower mapping quality and reduce variant-calling sensitivity. Aligned reads were then coordinate-sorted, assigned read-group information (based on sample, flow cell, lane, library), and had duplicates marked using Sentieon Dedup (equivalent to Picard MarkDuplicates). Following the GATK best practices, indel realignment and base quality score recalibration were performed using Sentieon Realigner and QualCal (equivalent to GATK IndelRealigner and BaseRecalibrator/ApplyBQSR), using dbSNP (v138) and the Mills/1KGP gold-standard indels as known sites, and original base qualities were retained in the OQ tag to allow reconstruction of original FASTQs submitted by the GCCs from the final BAM files.

*PacBio WGS & Fiber-seq:* PacBio whole-genome sequencing reads were processed using the same alignment pipeline established for the SMaHT benchmark study In brief, PacBio HiFi whole-genome sequencing reads were aligned to the GRCh38 reference genome (GCA_000001405.15), excluding ALT contigs and the hs38d1 decoy sequence, using pbmm2 (v1.13.0; https://github.com/PacificBiosciences/pbmm2), a PacBio-optimized wrapper around minimap2[14]. By default, all tags present in the unaligned BAM were carried through to the aligned output; the --strip flag was applied to remove nonessential kinetics tags (dq, dt, ip, iq, mq, pa, pc, pd, pe, pg, pm, pq, pt, pv, pw, px, sf, sq, st) and reduce file size, while retaining the methylation tags MM and ML and other assay-specific tags (for example, nucleosomes position in Fiber-seq data). Aligned reads were sorted by genomic coordinate, and read-group metadata specifying sample and library identity were assigned to each read.

*ONT WGS:* ONT whole-genome sequencing reads were processed using the same alignment pipeline established for the SMaHT benchmark study[15]. ONT long-read whole-genome sequencing reads were aligned to the GRCh38 reference genome (GCA_000001405.15), excluding ALT contigs and the hs38d1 decoy sequence, using minimap2 (version v2.26)[16]. To ensure consistency with the PacBio alignment pipeline, the following flags were applied: -Y to enable soft clipping for supplementary alignments; -L to produce long CIGAR strings (used in the CG tag); --eqx to represent matches and mismatches explicitly using = and X operators; and --secondary=no to suppress secondary alignments. Aligned reads were sorted by genomic coordinate, and read-group metadata specifying sample and library identity were assigned to each read. Methylation tags MM and ML were retained from the original unaligned reads and transferred onto the final alignments using Methylink (v0.6.0; https://github.com/projectoriented/methylink).

*RNA-seq:* Paired-end RNA-seq reads were aligned to the GRCh38 reference genome, excluding ALT, HLA, and decoy contigs, using the Sentieon (v202308.01) implementation of STAR (v2.7.10b)[17], in a pipeline adapted from the GTEx and TOPMed consortia. Alignment was performed against a STAR index built from GENCODE release 47 (GRCh38.p14) gene annotations[18], in two-pass mode (--twopassMode Basic, --twopass1readsN -1). The HLA region was also excluded from the reference genome to generate the STAR index files. Multi-mapping and splice-junction filtering were applied (--outFilterMultimapNmax 20, --alignSJoverhangMin 8), and chimeric-alignment detection was enabled (--chimSegmentMin 15 and associated chimeric-junction parameters). Alignment tags NH, HI, AS, nM, NM and ch were retained in the output (--outSAMattributes), and both genome- and transcriptome-aligned BAM files were generated (--quantMode TranscriptomeSAM GeneCounts) to support downstream transcript- and gene-level quantification. Duplicate reads were marked using Sentieon LocusCollector and Dedup, equivalent to MarkDuplicates in Picard (v2.9.0). Transcriptome aligned reads were used for quantification. To match the GTEx pipeline, quantification and QC were performed using two independent software packages. RSEM (v1.3.3) was used to generate gene- and transcript-level quantification, while RNA-SeQC (v2.4.2) was used to generate gene- and exon-level quantification as well as QC metrics. In GTEx, RNA-SeQC results are used for gene- and exon-level read counts and TPM values, and RSEM results are used for transcript quantification. Consistent with this approach, we release both sets of results, allowing users to select gene quantification from either RSEM or RNA-SeQC as needed.

*Kinnex:* For PacBio Kinnex long-read RNA-seq data, full-length non-chimeric (FLNC) reads were first clustered into high-quality consensus transcripts using the Iso-Seq cluster2 workflow. Both FLNC reads and consensus transcripts were then aligned to the GRCh38 reference genome (GCA_000001405.15), which excludes ALT contigs and hs38d1 decoy sequences, using pbmm2 (v1.13.0; wrapping minimap2 v2.26)[19] with the Iso-Seq preset (pbmm2 align --preset ISOSEQ --sort --strip --unmapped), which applies alignment parameters optimized for Iso-Seq/Kinnex data, sorts reads by genomic coordinate, removes nonessential kinetics tags, and retains unmapped reads. Following alignment, redundant transcripts were collapsed on the basis of shared exon–intron structure to define unique isoforms, isoform calls were filtered to remove likely artifacts and retain high-confidence transcripts, and the resulting isoform-level annotations were embedded directly into the aligned BAM files. Following alignment, redundant consensus transcripts mapping to the same genomic loci were merged into unique isoform models using IsoSeq collapse (v4.2.0; https://github.com/PacificBiosciences/IsoSeq), and the original FLNC reads were mapped back to isoforms to quantify full-length read support for each transcript, producing a read-to-isoform assignment file and transcript-support (FLNC count) statistics. Isoforms were then classified against the reference gene annotation using Pigeon (v1.3.0; pigeon classify; https://github.com/PacificBiosciences/pigeon), incorporating full-length read counts (--fl), CAGE-peak positions (--cage-peak), and polyA motif annotations (--poly-a) to assign each isoform to a reference gene and transcript and categorize it as known, novel, or a likely artifact. Isoforms were subsequently filtered (pigeon filter) to retain only high-confidence, non-artifactual transcripts together with their associated read-support counts, and the resulting isoform classifications and full-length read-support counts (recorded as the ct:i: tag) were embedded back into the individual aligned FLNC reads using an in-house script (FLNC_ImportTags.py), linking each high-confidence isoform to its supporting reads for downstream gene- and isoform-level expression analyses.

**S3.2 Quality control**

Per sample, alignment-based QC was performed using the same metrics and thresholds validated in the SMaHT benchmarking study[20]. All datasets released from the Network are of either “PASSED” or “FLAGGED” QC status, and datasets with “FAILED” QC status are not released. The QC thresholds were informed by the SMaHT benchmarking study, in which a threshold was determined to identify outliers in the QC metric distribution. Briefly, the mapping rate, duplication rate, and other alignment statistics (such as properly paired reads, average insert sizes) were calculated using Samtools (v1.17) stats/flagstat and Picard (v3.0.0; CollectAlignmentSummaryMetrics, CollectWgsMetrics, CollectInsertSizeMetrics), and the mean and per-base coverage were computed using mosdepth (v0.3.9)[21]. Sample identity was assessed with Somalier (v0.2.19)[22], flagging pairs with relatedness ≤90% as putative swaps or mislabels for manual review. Cross-individual contamination was evaluated with VerifyBamID2 (v2.0.1)[23] against 1KGP polymorphic sites. Assessments of the overall human sequence content and microbial contents were estimated using Kraken2 (v 2.1.3)[24]. Based on the benchmarking study, samples with contamination <1% were retained as suitable for somatic variant detection, while confirmed swaps or mislabeled samples were retracted. The thresholds for the following WGS QC metrics were determined based on the Benchmark WGS data: Aligned bases mismatch rate (0.08 for Illumina, 0.003 for PacBio, 0.01 for ONT), properly paired reads (92%), duplicate reads (15%), mapping rate (97%), mean insert size (250bp), and microbial contamination (0.1% for bacteria and 0.3% for virus, with overall human sequence content at 95% at the minimum). In addition for RNA-seq, the following QC metrics were examined: Median 3' bias (below 0.25 or greater than 0.75, for deviation from the expected 0.5), chimeric reads (1%), number of genes detected (25,000), estimated library complexity (50M), intergenic rate (0.1), exon/intron ratio (5), and rRNA rate (0.01). These thresholds were applied uniformly to determine inclusion of datasets in downstream analysis.

#### S3.3 DSA construction pipeline and validation

*[GCC-UW-SCRI] Fibroblast-derived PacBio HiFi+UL-ONT+Hi-C DSAs:* For the DSAs for 11 donors, we generated 60x PacBio HiFi, 30x Hi-C, and 60x ONT-UL data, with 20x of the latter from sequencing read lengths >100 kbp. Initial phased assemblies were generated via Verkko v2.2.1 and then filtered using NCBI FCS and BLAST to remove foreign contamination, mitochondrial sequence, Epstein-Barr virus, ribosomal DNA, and adapter sequences. The resulting high-quality phased assemblies have a median QV of 57.7 and a median contig N50 of 135.1 Mbp.

*[GCC-UW-SCRI] Evaluating quality of the DSAs:* We utilized multiple orthogonal assessments and different metrics to validate the quality of the assembly. We first assessed the contiguity of the DSA from N50 and NG50 metrics (based on 3.1Gbp genome size) using calN50 (https://github.com/lh3/calN50). Then, we calculated QV scores using the combination of Meryl v1.4.1 (meryl k=21 count ILLUMINA_READ.fastq.gz output READS_DB.meryl) and Merqury v1.3 (merqury.sh READS_DB.meryl HAP1_ASSEMBLY.fasta HAP2_ASSEMBLY.fasta MERQURY_OUTPUT) to estimate the sequence accuracy of the assemblies. QV values were derived as phred-scaled scores for each haplotype, based on error rates estimated by comparing the k-mer database generated from Illumina paired-end sequencing data from fibroblast with the assembly sequences. Next, we assessed the gene completeness of the assembled haplotypes using compleasm v0.2.6, based on the presence and integrity of 13,780 known single-copy orthologs included in the primates_odb10 database (compleasm run --mode busco -L DB_DIR -l primates_odb10 -o OUT_DIR -a HAPLOID_ASSEMBLY.fasta).

*[GCC-UW-SCRI] Donor-specific genome assembly graphs construction:* We constructed a donor-specific pangenome graph (DSG) for 11 samples for which a DSA was available using Minigraph-Cactus[25], without splitting by chromosome, clipping large unaligned regions or filtering rare variants. Each DSG contains four assemblies: the two DSA haplotypes, T2T-CHM13 and GRCh38. T2T-CHM13 is designated the primary reference scaffold for graph building. To identify assembly segments unique to the DSA (**Fig. 6b**) that are not present in GRCh38 or T2T-CHM13, we used halLiftover[26] to detect regions with no alignment to GRCh38/T2T-CHM13 in the DSG graph space.

*[GCC-UW-SCRI] Calling Fiber-seq peaks and evaluating haplotype-selective chromatin accessibility (HSCA):* Fiber-seq data was aligned to each DSA as described in section "Alignment of different sequencing data onto the DSA and post-processing", and Fiber-seq peaks were called using the FIRE pipeline (v0.1.2)[27]. Of the 11 samples with available DSAs, 8 had high quality Fiber-seq data in at least 2 tissues and were processed further. Tissues processed for each sample are listed below. (* = remained after applying the coverage filter described below)
SMHT004: LIVR*, LUNG*, MUSC*, TESL*
SMHT005: COAS*, BRHL*,LIVR*,LUNG*, TESL*
SMHT017: COAS*, CODS*, ESOP*, FBRO*, HART, LIVR*, LUNG*, MUSC*

SMHT018: BRCE, COAS, CODS*, ESOP, FBRO*, BRFL, HART*, LUNG*, MUSC, SKSE*, BRTL
SMHT020: BRCE*, FBRO*, HART*, TESL*
SMHT023: ADGL*, ESOP*, LIVR*, TESL*
SMHT040: ADGL*, BRCE*, CODS*, ESOP, BRFL*, HART*, LIVR, LUNG*, MUSC, OVAL*, BRTL*
SMHT043: AORT, BRCE*, COAS*, FBRO*, BRFL*, BRHL*, MUSC, SKNE, SKSE, BRTL*, TESL*

To identify haplotype-selective peaks and tissue-specific events in a single donor, we must compare peaks called in distinct genomic coordinate systems (the two haplotypes of the DSA), and across multiple tissue datasets. To do this, we performed consensus peak calling, described in detail in Lukas et *al*.[28]. All FIRE peaks called across tissues from a single donor serve as input into a graph-based framework, where peak calls are lifted between assembly coordinates to first (a) define a global set of 'consensus peak regions' -- assembly-agnostic genomic regions where a peak is called in at least one sample; and (b) pull raw chromatin actuation data at these consensus peak regions for each tissue and haplotype from pileups generated during the original, DSA-based FIRE runs. The final output table contains chromatin actuation information at each consensus peak region, for each tissue-haplotype combination, enabling us to directly compare chromatin actuation both between haplotypes and tissues. No comparisons between *donors* are being made at this step.

Several filters are applied to filter out low coverage tissues, define peaks, and define haplotype-selective chromatin accessibility. First, tissues with median coverage below 15 across all consensus peak regions are removed from downstream analysis. The remaining tissues are indicated with a * above. To define a Fiber-seq peak, we require haploid coverage ≥10 at that consensus peak region with at least 4 reads containing a FIRE element, and at least 25% chromatin actuation (reads with FIRE elements/total reads). To define a peak as haplotype selective (HSCA) in a tissue, both haplotypes must have coverage ≥10, with at least one called a peak (≥4 FIRE elements, ≥25% chromatin actuation), and a chromatin actuation difference of ≥25% between haplotypes. Additionally, a Fisher's exact nominal p-value, comparing the raw fraction of reads with FIRE elements in each haplotype, must be ≤0.005. We define a set of HSCA peaks for each tissue.

Next, we remove peaks associated with known imprinted regions[29], and split all HSCA peaks into those with identical sequence directly underlying the peak in both haplotypes (HSCA-noVar) and those that differ by at least one base (HSCA-Var). To ask if peaks which are HSCA in one tissue exhibit a tendency towards haplotype selectivity in others, and how this phenomenon compares for HSCA-Var and HSCA-noVar peaks, we plot the haplotype difference in accessibility in the tissue where the HSCA event was called against the haplotype difference at that peak in all other tissues, splitting by the Var/noVar groups. We then calculate the Pearson correlation across all HSCA events in both groups. High tissue-specificity of HSCA events is expected to show a low correlation, while more correlated chromatin accessibility differences between haplotypes across multiple tissues would lead to a higher correlation. We can calculate this correlation between two tissues (Figure S7c.1), between one tissue and all others (Figure S7c.1, right column), or combine all HSCA peaks across all tissues, for a single Pearson r value per donor (Figure 7c).

*[GCC-UW-SCRI] Obtaining various annotations of the DSA:* SD annotation was created using Sedef v1.1 (10.1093/bioinformatics/bty586). Repeat element annotation was created using Rhodonite (10.5281/zenodo.6036498), a snakemake workflow combination of RepeatMasker

v4.1.5 (http://www.repeatmasker.org), Tandem Repeats Finder (TRF) v4.09 and DupMasker, with a whole set of diploid DSA fasta as an input. RepeatMasker annotation was then used for soft-masking of the DSA. Subsequently we defined CpG islands regions in DSA based on the soft-masked fasta file using cpg_lh in UCSC Kent utility. In order to identify the genic region in the diploid DSA, we implemented LiftOff v1.6.3 to project GENCODE V47 annotation. We first separated out the diploid DSA into the different haplotypes and ran LiftOff (-copies -sc 0.95 -mm2_option="-a --end-bonus 5 --eqx -N 50 -p 0.5" -polish -cds -exclude_partial) on an individual haplotype, respectively. We merged the resulting projected annotation files together subsequently to obtain a full annotation for the diploid assembly.

*[GCC-UW-SCRI] Alignment of different sequencing data onto the DSA and post-processing:* Long-read sequencing data from each individual donor including PacBio HiFi, Fiber-seq, standard ONT and UL-ONT were aligned against diploid DSA of the same genetic origin using DSA-phasing-smaht pipeline (https://github.com/StergachisLab/DSA-phasing-smaht). Briefly, the pipeline performs (1) mapping to the diploid DSA using minimap2 (v2.29 or v2.31)[30] with --*preset lr:hqae*, (2) calculation of read depth using mosdepth (v0.3.14)[31] and (3) DNA methylation extraction from PacBio and ONT data using pb-CpG-tools (v3.0.0) and modkit (v0.6.3), respectively. After alignment, the pipeline additionally modifies the alignment tags, one of which is the mapping quality (MAPQ). Although the reads originated from a nearly identical genome, there are cases where the MAPQ reported by the aligner is 0, as reads derived from homozygous stretches longer than the read length are randomly assigned to one of the two haplotypes. In the case of alignment to the DSA, despite the ambiguity of such alignments, we reasoned that discarding such records would result in a loss of genomic information. We therefore reset the MAPQ of reads to 60 after performing the alignment. We also leveraged the PacBio data from fibroblasts which served as input for constructing each donor's DSA, to identify potentially misassembled regions of the DSA using NucFlag v1.0.0[32]. Additionally, we performed alignment of the Illumina paired-end short-read sequencing data to the DSA using bwa (v0.7.19; 10.48550/arXiv.1303.3997) mem and subsequently, duplicate reads were marked using SAMBLASTER (v0.1.26)[33].

*[GCC-WashU-VAI] Identifying distorted regions with read transport graphs built from DSAs:* In order to characterize where, and how often, read alignment is mishandled, we modeled the read transport process directly from high-quality DSAs with MoRGANA (https://github.com/juanfmacias/MoRGANA). MoRGANA works by first simulating reads from a DSA. It then aligns them using a specified aligner both to a reference assembly and back to the DSA. Because reads are simulated, the true origin of each is known. Aligning to an assembly yields read flow edges from source interval to aligned interval. Next it generated DSA to reference (GRCh38) alignments. This donor-to-reference genome alignment yields synteny edges. Each synteny edge links a donor interval to the reference interval where its reads should map to. Together these edges form a read transport graph, which describes how sequence from a donor gets handled by read alignment. From this graph, GOSSAMER (https://github.com/juanfmacias/GOSSAMER) derives two per-interval measures of fidelity. Source leakage is the fraction of an interval's reads that fail to reach their expected destination. Sink contamination is the fraction of the reads arriving at an interval that originated elsewhere. An interval is considered distorted when either exceeds 0.15.

We generated reads with the "deterministic lattice" read-simulation model implemented in MoRGANA. Rather than sampling fragment positions at random, the model tiles paired-end fragments along each DSA at evenly spaced start positions. The spacing between starts is the stride, and the offset of the first start within one stride is the phase. Reads were 150 bp, paired-end, and error-free, and the stride was 40 bp, which gives 7.5X coverage per phase (2L/S).

Fragment lengths were drawn from four presets, 255, 328, 384, and 457 bp, the quartile midpoints of a normal distribution with mean 356 bp and standard deviation 88 bp. Consecutive fragments cycle through these lengths and are interleaved, so the aligner sees the full spread of fragment lengths. This tiling gives uniform coverage and minimizes sampling noise, but a lattice at a single phase is biased toward its own start positions. We therefore simulated each DSA at eight phases and combined them into one graph, for 60X total. To choose the eight, we split the stride into eight equal windows and drew one phase at random from each, which keeps them spread across the stride rather than clustered. A fixed seed makes the choice reproducible, and no single phase was used on its own.

Reads were aligned with BWA-MEM (v0.7.17-r1188) to the GRCh38 primary assembly (hg38.primary25: chromosomes 1 to 22, X, Y, and M, with no alternate, decoy, or unplaced sequences) and, separately, to the donor's own assembly. Each DSA was aligned to GRCh38 with minimap2 (v2.28, -x asm10 -c --cs) to produce the genome alignment. Because a donor bin can align to more than one reference bin, synteny was weighted by aligned base pairs rather than assumed one-to-one. Each haplotype's transport graph was processed independently with GOSSAMER, at the same 1 kbp bin size the graph was built at. GOSSAMER computed the per-interval fidelity metrics and classified each interval into a transport motif. We exported the distorted intervals and merged adjacent ones into a per-donor set of distorted regions. These regions are in the DSA's scaffold coordinates. Because the somatic analysis runs in contig coordinates, we lifted them to contigs through each assembly's RagTag AGP.

**S3.4 Identifying somatic variants on the DSA space**

*[GCC-UW-SCRI] DSA-based sSNVs:* Variant calling was performed with DeepSomatic (v1.9.0) [34] using PacBio read data aligned to the DSA. For each donor, two DeepSomatic callsets were generated: one using only fibroblast read data, and one using PacBio read data from all other tissues merged with samtools merge (v1.23.1) [35]. These two callsets were intersected to generate the initial raw DSA-based sSNV callsets, ranging from approximately 50,000-150,000 sSNVs per donor before filtering. First, we removed all variants in regions assigned by NucFlag (v1.0.0 https://github.com/logsdon-lab/NucFlag) with any of these annotations: 'collapse', 'misjoin', 'low_quality', 'false_dup', 'het_or_mismap'. Next, we collapsed duplicate calls, where the same reference and alternate allele were identified at identical positions on homologous chromosomes or in paralogous regions, and removed any clustered variants, as defined by the presence of 3 or more sSNV calls in a sliding window of 1kbp. We then excluded any sSNVs in regions identified as distorted (see above “Identifying distorted regions with read transport graphs built from DSAs”). This initial round of filtering standardized callset sizes across donors to 30-40,000 sSNVs per donor.

To remove recurrent errors and germline artifacts, we compared our sSNV calls to the assemblies in the Human Pangenome Reference Consortium (HPRC) year 2 release [36]. We used jellyfish (v2.2.10) [37] to count 31-mers in agc-compressed [38] HPRC assemblies, as well as for each DSA. For each donor, we subset their DSA into satellites, SDs, or other repeats and generated 100,000 random 31-mers from each region to use as controls. We counted each control 31-mer in the HPRC samples and in the donor’s own DSA, calculating the log-fold change in the HPRC relative to the DSA. We then generated two sets of three 31-mers for each sSNV call, one set with the reference and one with the alternate allele, where the sSNV was at the first, middle, or last position of the 31-mer. We counted these variant 31-mers in the HPRC samples and in each donor’s DSA and compared them to our control distributions. For each set of reference or alternate 31-mers, we took the minimum observed count in the HPRC; if a 31-mer was not represented in the HPRC, the corresponding variant immediately passed to the next round of validation, and variants in unique space immediately failed if they were counted in

the HPRC even once. For 31-mers in repeat regions, we calculated their log-fold change and compared it to the control distribution by calculating the empirical p-value. For each region, we retained variants with empirical p-value <0.1, then applied multiple-testing correction with the Benjamini-Hochberg procedure. Any variant with p-value <0.05 after multiple-testing correction remained in our callset. Importantly, if a variant's 31-mers with the alternate allele were observed in at least half of HPRC haplotypes while 31-mers with the reference allele were observed in less than 10% of HPRC haplotypes, we flipped called variant's reference and alternate allele. This filtering step reduced the callset to 19-25,000 sSNVs per donor.
With this reduced callset, we used samtools to subset DSA-aligned PacBio and Illumina crams for every tissue from each donor, retaining only reads aligned to variant positions and their homologs, then used vg (v1.65) [39] to surject reads onto homologous chromosomes, resulting in an alignment file where each read is represented twice: once at the original aligned location, and once at the homologous position, provided that such a site exists. We retag these resulting crams with samtools calmd, then individually filter and rephase every read. For PacBio reads, we require base quality >20, read length >10kbp, gap-compressed identity to the reference >0.99, and aligned fraction >0.9. For Illumina reads, we require base quality >20, read length of 151bp, gap-compressed identity to the reference >0.98, aligned fraction of 1, and distance from read end >10bp. For reads that meet those thresholds, we compare both the alignment score and gap-compressed identity to both haplotypes. If a read has identical scores on both haplotypes, we consider it unphased.

For each variant site, we count the number of passing reads that support the reference and alternate allele on each haplotype, as well as those that cannot be unambiguously phased. Using those read counts, we determine if a variant is supported in each of a donor's sequenced tissues. We consider a variant to be high-confidence if it is supported by at least 1 PacBio read and 3 Illumina reads, provided both types of sequencing data are available. A variant is medium-confidence in a tissue if supported exclusively by PacBio (≥1 reads) or Illumina (≥5 reads), and low-confidence if below read thresholds. After evaluating a variant in each tissue individually, we combine data across tissues, using it to strengthen or weaken variant confidence. If a variant is high-confidence in at least one tissue or is medium-confidence in at least two tissues, all medium-confidence tissues are retained. If a variant only has low-confidence support across tissues, it is excluded from the callset. In addition, we remove variants that are phased to opposite haplotypes across tissues or sequencing platforms, as they are likely to be recurrent errors in sequencing or mapping. We find 8-10,000 sSNVs per donor are confidently supported across one or more tissues.

After confirming variant presence, we perform a binomial test, combining either PacBio or Illumina read data across all tissues with a variant. If the variant is phased, we only evaluate reads from the mutant haplotype, using $H_0=0.9$ for the binomial test (one-sided), compared to $H_0=0.5$ (two-sided) for unphased reads. After applying multiple testing correction with the Benjamini-Hochberg procedure, we retain all variants with p<0.01 for both types of read data. In addition, we use a two-sided Fisher's exact test, to evaluate strand-bias in Illumina data from tissues with the variant, retaining variants with p>0.01 after Benjamini-Hochberg correction. We then remove any sSNVs that are located at edges of homopolymers, either at the first or final position, or at the first adjacent position on either side, if the alternate allele matches the homopolymer subunit. Finally, for variants that can be represented in GRCh38, we evaluate their presence in gnomAD (v4) [40], excluding all variants with frequency $>10^{-4}$. This results in a final callset of 3-5,000 sSNVs per donor.

Finally, we determined the callable genome size for each donor by taking the full assembled genome size and subtracting regions annotated by NucFlag (see categories above) and distorted regions. For each genomic region, we used bedtools intersect (v2.31.1) [41] to define both the amount of callable base pairs, and which sSNVs are in the region. We calculated the mutation rate simply by dividing the total number of mutations in a region by the number of callable base pairs. Enrichment significance was calculated using a paired two-sided t test, comparing a given region's mutation rate to the genome-wide rate, and then corrected for multiple testing with the Benjamini-Hochberg procedure.

*[GCC-UW-SCRI] DSA-based sSVs:* To identify sSVs using the DSA, the aligned BAM file from each donor/tissue/platform combination was first realigned and haplotagged with LongcallD (v0.0.8)[42], which also produced an sSV callset. The resulting refined BAM was then used as input for the remaining three sSV callers, DELLY (v1.5.0) [43], Severus (v1.6) [44], and Sniffles2 (v2.7.0) [45], producing four initial sSV callsets per bam. Per sample, per caller VCFs were merged and normalized within each tissue (BCFtools, v1.23.1), and collapsed with Truvari (v4.3.1, -refdist 1000, -pctseq 0, -pctsize 0.7, type agnostic) [46] to remove near-duplicate site representations across callers. Within each tissue, sites supported by at least two of the four callers were retained. The union of all 2-of-4 sites across all tissues for a donor was used for joint genotyping across all of a donor's samples using kanpig (v2.1.0, mosaic mode) [47] to identify evidence for that sSV across all tissues for that donor. The result was a single, tissue and caller integrated VCF per donor with all sSVs with multi-caller support.

These sSVs were then further filtered to remove artifacts (**Fig. S20**). Using NucFlag (v1.0.0) and Flagger (v1.2.0)[48] annotations of each DSA, sSVs were removed if >10% of the variant overlapped a region that was collapsed or misassembled. Using PAV (v2.4.6.5) [49], we identified inherent germline differences between GRCh38 and each DSA, and sSVs matching those calls were also removed. Variants present in >70% of tissues that also had an hVAF of either ≥0.9 or between 0.4-0.6 were filtered out, as these patterns are also likely to reflect germline artifacts. Variants lacking kanpig genotyping support were removed, except those >1000bp or not of type INS/DEL, which were retained irrespective of genotyping support since kanpig is not optimized to genotype variants of that size or type. Variants on contigs with an unusually high number of calls relative to other contigs for that donor (Z-score >3) were also removed, as such enrichment likely reflects an assembly quality or alignment issue rather than genuine variation. sSVs located in repeat regions were annotated using RepeatMasker (v4.2.4, http://www.repeatmasker.org) based on 70% or more overlap with a repeat region. For variants in tandem repeat regions, support from both Severus and LongcallD was required, given their improved algorithm to detect sSVs in these regions [50]; satellite variants with a RepeatMasker annotation of SAR or BSR_Beta were also removed, as these subtypes occur on acrocentric short arms, where assembly reliability is reduced.

*[GCC-UW-SCRI] Identifying somatic structural variations in alpha-satellite regions of the centromeres:* In addition to DSA-based sSVs laid out above, sSV within the α-satellite of the centromeres were identified following the method from Sohn et al., 2025 [51]. Briefly, for each DSA we first determined the approximate location of the centromere by lifting the T2T-CHM13 centromere annotation onto the assembly with rustybam [52] (*rb liftover*), using the pairwise-alignment information between T2T-CHM13 and the DSA. The leftmost and rightmost coordinates of the lifted intervals in the DSA space were taken as the candidate centromeric region, and regions annotated by RepeatMasker as ALR_Alpha with ≥100 kbp in length were retained, which then used as inputs to HumAS-HMMER (https://github.com/fedorrik/HumAS-HMMER_for_AnVIL) to annotate α-satellite higher-order repeat (HOR) monomers.

Subsequently, sSV within α-satellite repeats were identified directly from PacBio long-read alignment to the DSA, by parsing out MD tags encoding mismatched and deleted reference bases as well as CIGAR strings using a custom Python script to extract insertion and deletion events of 50 bp or more. Each potential sSV was classified as α-satellite monomer-sized if its length fell within 5% of an integer multiple of 171 bp. We then further applied filtering to get the pruned sSV records leveraging read-to-DSA alignment quality metric; such as read length (≥10 kbp), gap-compressed sequence identity (≥0.998), the fraction of the query read aligned (≥0.995) and the relative position of each sSV within the read (inner 70%). We removed sSVs overlapping with regions flagged by Flagger and NucFlag, and calls shared between the fibroblast and tissue from the same individual, retaining a unique set of sSV within each tissue against the fibroblast. The interval of each sSVs was padded on both sides by its own sSV length to absorb breakpoint uncertainty that scales with event size, and the records with overlapping padded intervals were merged into positional clusters if they are in the identical unit size of α-satellite monomer (171 bp). For non-unit-sized sSVs, on the other hand, calls within 10% of each other in length were collapsed together.

To assess the distribution of these sSVs relative to the kinetochore binding domains, that is, CDRs, we first defined them using UL-ONT CpG methylation across α-satellite arrays called by modkit (pileup --combine-strands --cpg). Specifically, we merged α-satellite arrays within 10 kbp of one another and then computed the CpG methylation in 1 kbp bins. Per-α-satellite stretch z-scores calculated from these bins were used to determine CDR boundaries with a z score threshold of –1.25, so that hypomethylated bins are selected. To ensure that CDRs inferred from donor's fibroblast are consistent across different tissues within the same individual, we also determined per-tissue CDRs using the PacBio CpG methylation data generated with pb-CpG-tools, following the same per-α-satellite 1 kbp z-score approach. sSVs within α-satellite were then labeled as CDR-overlapping or non-CDR depending on their intersection with the CDR intervals obtained from each individual's fibroblast UL-ONT data. When computing the rate of sSVs within α-satellite HORs for each tissue, we normalized the number of sSV events by dividing them by the product of the total length of α-satellite in the DSA and the mean read coverage across α-satellite (only using reads with length ≥10 kbp) thereby accounting for the regional depth of coverage. Samples with coverage lower than 5 were discarded from the analysis and only samples with coverage of at least 8 were considered when calculating fold enrichment of α-satellite sSVs within CDRs relative to non-CDR regions. Autocorrelation was computed for each sample by tabulating the number of unique sSVs equal or greater than 100 bp at each integer base-pair length and calculating the autocorrelation function over this length distribution using pandas autocorr() method.

### Supplementary Methods Section 4 | Somatic variant detection

#### S4.1 Pipeline details - bulk sSNV calling

_SNV/indel germline variant annotations:_ Germline variant calling used for SNV filtering was done with DNAScope Hybrid [53], leveraging both short- and long-read data for each donor. Specifically, heart tissue was used for 19/25 donors with descending colon used for SMHT005, SMHT023, and SMHT029 and muscle used for SMHT015, SMHT039, and SMHT043. These tissues were selected based on an analysis of the most concordant tissues which also most often had matched short- and long-read data for the 25 donors.

_Types of Variant Evidence:_ Within each vcf, there are 4 "Cross" tags described in detail here which describe different lines of evidence that were evaluated for each sSNV call. **CrossTech**: there was sufficient variant read support in both short and long read technologies. As not every tissue had long read data available, for each donor we pooled all long-read PacBio data for every tissue to evaluate this tag as well as for use in the long-read phasing filter. **CrossTissue**: determining which variants are found in multiple tissues of a single donor. This tag was evaluated only for variants within easy regions and required a strict alt read support as it can easily be misled by recurrent short-read alignment errors. **CrossCore**: this tag is present if a variant was called by a variant caller in multiple distinct cores. **CrossCaller**: this tag is present if multiple variant callers agreed upon this variant.

#### S4.2 Pipeline details - Duplex sSNV calling

*CompDuplex-seq and NanoSeq:* Variant calling and filtering were performed using a unified CompDuplex–NanoSeq analysis pipeline, available at https://github.com/zonglab/CompDuplex. The workflow includes adapter trimming, alignment to GRCh38, consensus molecule generation, duplex validation, and stringent variant filtering based on base quality, mapping quality, and duplex support, ensuring high-confidence detection of ultra-rare mutations.

*META-VISTA-seq:* META-VISTA-seq data were processed, and sSNVs and INDELs were called, using the META-VISTA-seq pipeline v2.0.0. Read pairs were first preprocessed to identify the duplex barcode carried by each read against the reference set of 136 forward–reverse Tn5 barcode combinations, trim it from the read sequence, and write the forward–reverse barcode pair into the FASTQ comment field as a BC:Z: tag; overlapping mates were merged in the same step. Preprocessed reads were aligned to GRCh38 no-alt (GCA_000001405.15) with bwa-mem (v0.7.18; -C, primary) and minimap2 (v2.30-r1287; -ax sr -y, used for SNV cross-aligner confirmation), whose -C and -y options transfer this comment into the BAM BC tag. The unmerged read stream was aligned separately with bwa-mem for use by the merge filter. Pileups were generated per barcode from the BC tag, with the two duplex orientations of each barcode pair pooled into a single column, requiring mapping quality ≥ 30, base quality ≥ 30 (≥ 20 for the matched bulk column), soft-clipping ≤ 10 bp, and ≥ 4 reads per allele, and were restricted to the GIAB GRCh38_notinalldifficultregions mask. Calling was performed in pooled mode and restricted to autosomes. sSNVs required ≥ 4 ALT reads with ≥ 2 per strand, ALT allele balance ≥ 0.01, position ≥ 10 bp from the read end, ≤ 1 conflicting duplex read, mapping quality ≥ 50, concordant support from both aligners, ≤ 1 call per 100 bp window, and, in the donor-matched bulk WGS, depth ≥ 15 with zero ALT reads. INDELs used the same thresholds, were called from the bwa-mem alignments only, were restricted to biallelic sites overlapping the merged-fragment window, and were excluded within max(5, 2 × INDEL length) bp of a germline indel. Only calls supported on both strands were reported as somatic mutations; single-stranded and joint multi-cell calls were tracked separately. Three post-hoc filters were applied. First, variants were removed unless at least one supporting molecule placed them ≥ 12 bp from its nearest Tn5 insertion boundary. Second, positions matching gnomAD v4.1 sites at allele frequency ≥ 0.01 were removed. Third, each supporting barcode was re-piled up in the unmerged alignment, retaining double-stranded calls genotyped as homozygous ALT and single-stranded calls genotyped as heterozygous.

*CODEC:* CODEC samples were analyzed from BCL files through to variant calls using best-practice workflows from the standard CODECsuite (https://github.com/broadinstitute/CODECsuite). All CODEC data were processed through a cloud-based pipeline implemented in Terra using the Workflow Description Language (WDL). The pipeline is publicly available on Dockstore at https://dockstore.org/my-workflows/github.com/broadinstitute/TAG-public/SingleSampleCODEC.

#### S4.3 Pipeline details - sMEI

*sMEI callset generation:* We generated candidate sMEI callsets from long-read WGS data of 25 SMaHT donor tissue samples using PALMER (v2.3.3) [54] and longcallD (v0.0.8). For PALMER, candidate sMEIs were first identified using the default parameters (PALMER --input ${CRAM} --workdir ${WORKDIR} --ref_ver GRCh38 --output ${PREFIX} --type ${MEI} --mode raw --chr ${CHR} --ref_fa ${REF}). Only calls supported by at least one high-confidence target site duplication (TSD)-supporting read were retained. Calls located within low-confidence genomic regions were also excluded. For longcallD, the -s option was used to enable somatic variant detection, and the -T option was used for MEI calling (longcalld call -s -n ${PREFIX} -o ${PREFIX} -T ${AluY_L1_SVA_cons_noPA.fa} ${REF} ${CRAM}). Then the calls annotated as "Somatic" in the "INFO" field were retained for the following analyses.

*sMEI haplotype phasing:* To distinguish somatic from germline MEIs, we performed haplotype phasing using long-read WGS data. Single-nucleotide polymorphisms (SNPs) were first identified within a 40 kbp window centered on each insertion site using GATK4 HaplotypeCaller. Up to five high-confidence heterozygous SNPs (QUAL>500) closest to the insertion site were selected for phasing. Long reads spanning both a selected heterozygous SNP (hetSNP) and the insertion site were then used to determine the haplotype of the supporting reads carrying the insertion. Phasing was performed independently for each selected hetSNP, and the final classification was assigned according to the most frequently observed outcome across all phased SNPs. Insertions were classified as 'No hetSNPs' when no hetSNPs were identified within the 40 kbp window, and as 'No covering reads' when no long reads simultaneously spanned both the insertion breakpoint and a hetSNP. Insertions supported by both haplotypes were classified as 'Insertion in multiple haplotypes'. Insertions linked exclusively to a single haplotype, while the alternate haplotype was covered by reads lacking the insertion, were classified as 'Germline'. Finally, insertions linked to a single haplotype but supported by only a limited number of reads were classified as sMEIs, consistent with low-frequency somatic insertions. To avoid excluding true somatic events, only candidates confidently classified as germline ('Germline' or 'Insertion in multiple haplotypes') were removed, as these classifications provide strong evidence for germline origin. Additionally, to exclude the calls that arose from misalignment or misclassification during haplotype phasing, we cross-checked the haplotype phasing results across the tissue samples from the same donor and the calls classified as germline in at least one tissue were also classified as germline in other tissues.

sMEI target-primed reverse transcription (TPRT) and structural feature annotation

To identify target site duplication (TSD), synthetic paired-end reads were generated for each candidate insertion. Briefly, 500 bp of reference sequence upstream and downstream of the insertion site were extracted as the left and right pseudo-reads, respectively. The first 100 bp of the insertion sequence was appended to the left pseudo-read, whereas the last 100 bp was prepended to the right pseudo-read. The resulting pseudo-reads were aligned to the reference genome using BWA-MEM with default parameters, and the corresponding CIGAR strings were examined. A TSD was defined when the alignment match (CIGAR operation M) extended beyond the original 500 bp reference sequence, allowing the duplicated sequence and its length to be inferred. PolyA/T tails were identified by searching for homopolymeric tracts of ≥10 A (adenines) at the 3′ end or ≥10 T (thymines) at the 5′ end of the insertion sequence after excluding the annotated TSD. Up to two mismatches were permitted within the homopolymer tract. To annotate L1 structural features, insertion sequences were aligned against the corresponding mobile element consensus sequences obtained from RepeatBrowser[55] using BLAST. The degree of 5′ truncation was inferred from the 5′-most aligned position on the consensus sequence, and 5′ inversions were identified by the presence of reverse-orientation alignments at the 5′ end of the insertion.

*Raw-read level alignment signal rescue:* To recover sMEI candidates missed by individual MEI detection methods, we searched for supporting raw-read alignment signals across all available sequencing platforms (PacBio, ONT, and Illumina) and tissues from the same donor. For each candidate insertion, clipped sequences within ±30 bp of the breakpoint were extracted from aligned reads and compared with the insertion sequence identified in another platform or tissue.

Candidates with matching clipped-read evidence were considered supported at the raw-read level, even in the absence of a formal MEI call. Insertions supported by either an MEI call or raw-read level alignment evidence in two or more tissues from the same donor were classified as multi-tissue events.

Constructing the donor-specific FL-L1HS/L1PA2 catalogue: Because donor-specific assemblies (DSAs) were not available for all donors, we performed local assembly at candidate full-length L1HS/L1PA2 (FL-L1HS/L1PA2) loci to construct a donor-specific FL-L1HS/L1PA2 catalogue. To maximize the coverage of potential FL-L1HS/L1PA2 loci, we compiled candidate loci from three sources. First, we included reference L1HS/L1PA2 elements longer than 5,500 bp based on RepeatMasker annotations of the human reference genome (n=1,378). Second, to identify FL-L1HS/L1PA2 loci that may be absent or incompletely represented in the reference genome, we collected FL-L1HS/L1PA2 positions identified in the SMaHT DSAs and lifted these coordinates over to the reference genome using paftools (n=252). Third, we identified non-reference FL-L1HS/L1PA2 insertions using PALMER (v2.3.3) and LongcallD (v0.0.8). Putative somatic MEIs were excluded using an in-house phasing method for PALMER (v2.3.3) calls and the SOMATIC annotation provided by LongcallD (v0.0.8). We retained insertion loci supported by more than five insertion-supporting reads in PacBio or ONT data and generated a union set across all tissues from each donor. We further retained L1HS or L1PA2 insertions longer than 5,500 bp that were detected in more than three tissues. These non-reference insertion positions were then merged across donors (n=227). Candidate loci identified using these three approaches were merged to generate the final set of FL-L1HS/L1PA2 loci for local assembly. For each candidate locus, we extracted all PacBio reads mapped within ±10 kb of the locus with a mapping quality (MAPQ) of 60 by donors. The extracted reads were converted to FASTA format, and 100 reads were randomly selected per locus to reduce computational burden. The selected reads were then locally assembled using hifiasm (v0.25.0-r726) with the options -f0 -l0. RepeatMasker (v4.2.2) was subsequently used to annotate L1HS and L1PA2 elements within the assembled contigs. The resulting locally assembled FL-L1HS/L1PA2 sequences constituted the donor-specific catalogue used for subsequent source tracing of somatic L1 insertions.

Source tracing of sMEI using both transduction and L1 body sequence: Source tracing of sMEIs was performed as previously described, with minor modifications[56]. Briefly, the insertion sequence was analyzed using RepeatMasker (v4.2.2) to separate the L1 body sequence from the transduction sequence. For source tracing based on the transduction sequence, the transduction sequence was further examined to determine whether multiple transduction events had occurred by identifying poly(A/T) signals within the transduction sequence. In such cases, only the terminal transduction sequence of the insertion was extracted. These sequences were then mapped to the reference genome, and the best hit located within 10,000 bp of a FL-L1HS catalogue was considered the candidate source.
For source tracing based on the L1 body sequence, the body sequence was extracted using BLASTN (v2.14.1+) against the TE consensus sequence. Poly(A/T) tails were subsequently trimmed, allowing up to two mismatches within 5bp minimum poly(A/T) tail length. The resulting sequences were then searched against the donor-specific FL-L1HS catalog using BLASTN (v2.14.1+). For locus-level source assignment, BLASTN (v2.14.1+) results were first collapsed by locus, retaining the alignment with the highest bitscore for each locus. A source locus was assigned when the best-matching locus had a higher bitscore than the second-best distinct locus and its percentage identity was greater than or equal to that of the second-best locus.

CpG methylation profiling at the L1HS promoter: To profile CpG methylation at both reference and non-reference FL-L1HS promoters, we extracted reads aligned to donor-specific FL-L1HS loci with a mapping quality (MAPQ) greater than 60. These reads were then realigned to the donor-specific FL-L1HS sequences. CpG methylation was subsequently called using modkit (v0.6.1) for Oxford Nanopore Technologies (ONT) whole-genome sequencing data and pbcpgtools (v3.0.0) in reference mode for PacBio sequencing data.

*sMEI calling from TEnCATS for SMHT005:* To identify candidate somatic L1HS insertions from TEnCATS data, we used Nanopal as described in McDonald et al.[57] with modifications detailed below. The TEnCATS sequencing data for each tissue (3 PromethION flowcells per tissue) were pooled into a single dataset per tissue. For each tissue, reads were filtered for QC with Samtools (version 1.22.1 with parameters: --expr '[qs] >= 9') and Minimera (version v0.4.0 from https://github.com/Boyle-Lab/minimera/ with parameters: --monotonony-threshold 0.5), aligned to the hg38 reference genome using Minimap2 (version 2.28 with parameters: -aYy --eqx --secondary=no -x map-ont) then filtered to find reads containing non-reference MEIs using PALMER (version ed90190625 from https://github.com/WeichenZhou/PALMER/) and BLASTn (version 2.15.0). Individual reads were clustered by primary alignment location to call putative non-reference L1HS insertion sites and the number of supporting reads, producing one callset of insertions per tissue. We compared these callsets to identify possible somatic insertions by filtering for locations with at least three supporting reads in one tissue and no supporting reads in at least half of the tissues. The resulting candidates were then manually inspected and filtered in IGV to produce the final list of insertions.

**S4.4 Pipeline details - sSV**

*SV Discovery, Merging, Genotyping, and Annotation:* Tissues from individuals were sequenced with ONT or PacBio sequencing technologies across multiple sequencing centers. Some combination of tissue, core, technology, and center constitute a single sample. Each aligned BAM from samples was run through three SV discovery tools: DELLY [58] (v1.7.3, delly lr --outfile ${PREFIX}.delly.bcf --genome GCA_000001405.15_GRCh38_no_alt.fa.gz --technology ${TECH} ${BAM}); Sniffles [59] (v2.7.5, sniffles --input ${BAM} --vcf ${PREFIX}.snif.vcf.gz --threads 8 --minsvlen 40 --output-rnames --mosaic --mosaic-af-min 0.001 --mosaic-include-germline --minsupport 1 --cluster-binsize 50 --cluster-merge-len 0.10 --no-qc --tandem-repeats snf2-GRCh38.trf.bed --reference GCA_000001405.15_GRCh38_no_alt.fa.gz --sample-id ${PREFIX}); Severus [60] (v1.6, severus --target-bam ${BAM} --out-dir ./${PREFIX}.sevr.vcf.gz --threads 8 --min-sv-size 40 --vntr-bed severus-GRCh38.trf.bed.gz). With BAM and PREFIX referring to each samples' alignment and name, while TECH being either 'pb' for PacBio and 'ont' for ONT sequencing. The resultant VCFs from all samples processed for a donor were merged using BCFtools [61] v1.21 merge -F x -m none. These VCFs were then run through an initial round of genotyping with kanpig [62] v2.1.0 and parameters --maxpaths 500 --neighdist 500 --bandwidth 5 --maxclust 5 --maxcoverage 2000 --sizemin 40. sSVs with less than two reads of support across all samples were discarded. This initial genotyping step served as a filtering procedure to reduce the number of sSV representations given to the next step of collapsing putatively redundant sSVs using truvari [63] v5.4.1 collapse with parameters --sizemin 40 --dynthresh 5,30,50,1500 --keep maxqual. Here, the maxqual parameter chose the representative sSV from a cluster of matching sSVs as the one with highest alternate read support. The collapsed set of sSVs were then genotyped again using kanpig and the same parameters above in order to produce the final set of genotypes. Next, additional columns were added to the VCF with consolidated information across groups of samples. First, all reference and alternate allele coverage observed across all samples was summed and used to produce the individual-level genotype using kanpig's API. For convenience, coverage from samples comprising a single tissue were summed to produce tissue-level genotype fields.

Six variant level filters were applied to the VCF. First, sSVs below 50bp were marked as SMALL. Second, sSVs which were not discovered by at least two discovery tools or had cumulative coverage less than 6x were marked LOWDC. Third, sSVs with only 2x alternate read support were marked LOWDP. Fourth, sSVs with breakpoints within genome regions marked as ARD (described below) were filtered. Next, sSVs in neighborhoods (1,000bp) with at least 10 sSVs were marked as being in high density regions (HIDEN) and excluded due to the inherent complexities of accurate assignment and assessment of read support in these regions. Finally, sSVs with ≥90% sequence/size similarity of the same type were marked as either putatively redundant (REDUN) or split (SPLIT) representations.

The assembly-to-reference disagreement regions were built from aligning 231 haplotype resolved long-read assemblies from the Human Pangenome Reference Consortium (cite) to GRCh38 using minimap2[64] v2.28-r1209 -asm 5. Per-haplotype alignments for each sample were parsed and high-confidence regions of GRCh38 were classified as those covered by exactly one alignment for each haplotype with considerations for the sex of the sample and chrY / chrX pseudoautosomal regions. High-confidence regions across all haplotypes were consolidated with bedtools[65] and regions with fewer than 95% samples producing high-confidence regions were blacklisted by the ARD filter.

Additional annotations generated included the number of neighboring sSVs within 1,000 base-pairs using truvari anno numneigh; sSV class (e.g., Alu/TR) and gene overlap annotations were generated with default parameters on three tools: truvari anno trf using the adotto TR catalog[66]; SVAN[67]; AnnotSV[68].

_Refining somatic candidates:_ The process of discovery, merging, genotyping, and annotation (DMGA) produces a project-level VCF of SVs with genotypes indicating the germline (heterozygous/homozygous alternate) or low variant allele fraction (VAF; homozygous reference, with less than ~20% of read support on the alternate allele) for every sample, across each tissue, and cumulatively across all available donor sequence. The sSVs with a low-VAF at the donor-level which pass the variant-level filters described above and with at least one sample having ≥10x covering reads were selected as somatic candidates. These sSVs' start/end positions were buffered by 500bp and then merged with bedtools v2.30[69] to create candidate somatic loci. In order to minimize alignment ambiguities artificially amplifying the occurrence of low-VAF sSVs, we refined the alignments per-sample over the candidate somatic loci using LongCallD[70] v0.0.11 and parameters -s n ${sname} -T ${alu_file} -M 20 --refine-aln --region-file ${somatic_regions} -b ${sname}.bam ${is_ont} where 'sname' is the relevant sample name 'alu_file' is the longcalld provided AluY_L1_SVA_cons_noPA.fa, and 'is_ont' marked to true on ONT sequencing experiments. An additional run of sniffles SV discovery was performed on the LongCallD refined BAM with parameters -input ${sname}.bam --vcf --cluster-binsize 50 --mosaic-af-min 0.001 --mosaic --mapq 20 --minsvlen 40 --mosaic-include-germline --cluster-merge-len 0.10 --minsupport 2 --allow-overwrite. The resultant refined BAM and two discovery VCFs were then provided to the DMGA step described above. This second round of analysis on harmonized alignments reduces the amount of alignment ambiguity and serves to challenge the low-VAF signal observed in the initial round of analysis by removing poorly represented initial somatic candidates. Any sSVs surviving the second DMGA step becomes the updated list of somatic candidate sSVs.

_sSV Filtering:_ A total of 6 filters were applied to the candidate sSV in order to remove potential germline contamination and prioritize higher confidence sSVs. The set of somatic exclusion criteria were sSV which had no tissue producing ≥10x total coverage, presence in the 1KGP panel-of-normals, median supporting read mapping quality less than 60, evidence of batch effects across long-read sequencing platforms, and sSV which failed a phasing test and therefore was not likely supported by reads from a single haplotype.

For the 1KGP panel-of-normal, the long reads of 1,008 samples from two projects[71,72] were downloaded. Every 1KGP sample was genotyped against each donor's SV VCF using kanpig v2.1.0 and parameters gt --neighdist 500 --maxpaths 1000 --pileupmax 30 --seqsim 0.90 --sizesim 0.90 --ab 0.20. Every SMaHT donor sSV that was not private against the 1KGP samples (AF == 0) was excluded from the final set of sSVs.

To check for platform batch effects, reference/alternate read counts for each sSV were compared across long-read sequencing platforms (HiFi and ONT). For sSVs which were only present in a subset of tissues that did not have available technical replicates, no batch-test was performed. For sites where both platforms produced alternate read support, a two-sided Fisher's exact test was performed on the 2×2 contingency table of reference/alternate reads by platform. For sites where only one platform had alternate read support, a Beta-Binomial model was used, in which a Beta(alt+1, ref+1) posterior was derived from the platform with alternate read support (uniform Beta(1,1) prior), and the probability of observing zero alternate reads at the second platform's observed depth computed as the posterior predictive probability P(X=0). Sites with a significant result (alpha < 0.05) under either test were flagged as showing a potential batch effect.

For the phasing test, an sSV was considered to be likely from a single haplotype if it passes a one-sided binomial test of P(X ≥ k_minor | X ~ Binom(n, error_rate)) where k_minor is the number of reads from the less populated 1/2 haplotype, n is the total number of phased reads, and error_rate = 0.07. If ≥ 30% of sSV supporting reads were unphased, the site is deemed to have poor haplotagging quality and the sSV is assumed to be likely from a single haplotype.

For eight of the donors, their DSA was procured and two additional filters applied. These DSA subtraction filters aim to identify germline contamination (i.e., genotyping errors) in the discovered sSV. The first filter aligns each haplotype to GRCh38 with minimap2 v2.28-r1209 parameters -cx asm10 --secondary=no --cs and calls SVs with paftools parameters -q 60 -L10000. The resultant VCFs per-haplotype are consolidated to construct a set of germline SVs which are then subset to those within regions singly-covered per-haplotype and intersected with the donor's discovered SVs using truvari v5.4.1 parameters --pick multi. Any discovered sSV that matches a DSA derived SV is marked as MatchDSA. The second filter takes all reads which support a discovered sSV and aligns them to each DSA haplotype independently with minimap2 v2.28-r1209 and parameters -MD -a -x map-hifi|map-ont. Any read that aligns on either haplotype using ≥90% of its query sequence and contains no insertion/deletion ≥20bp is deemed as hitting the DSA. Any sSV with ≥50% of its supporting reads hitting the DSA is filtered as HitDSA.

*TR Calling:* Tandem repeats allele length were extracted based on the adotto v2.1 TR catalog, following the same strategy for both PacBio and ONT. We utilized an anchor sequence of 200 bp and stored pileups of aligned reads over each locus to generate a delta difference to the reported size in the catalog. These length deltas were used to compute TR allele-sizes, which were aggregated to perform unsupervised clustering of them. This resulted in 1...N clusters per-locus. Each locus was then classified based on the number and size of the clusters, focusing on loci showing instability (i.e., multiple clusters). Finally, per-locus the allele-sizes were distributed back to each sample, to then perform QC to detect platform bias (seen only in one sequencing platform) and low support (i.e single read clusters). Finally, loci classified as highly unstable were annotated (i.e., gene overlap) and manually inspected. For the sSV sites overlapping genes (AnnotSV) and tandem repeats (adotto TR catalog v2.1), long reads were classified into somatic/germline using stablevizer (https://github.com/ACEnglish/stablevizer) with tissue read minimum of two and -q 0. Manual inspection of sites with instability in at least two donors, unstable reads in both long-read platforms and no bias in by-platform depth (fisher exact test of ONT/HiFi stable and unstable read support $p < 0.05$) were selected. The prominent tissue annotation was selected as those with the greatest magnitude of median change relative to reference allele and somatic read length spread, with 'SKI' indicating both calf and abdominal skin sites.

**S4.5 Pipeline Details - Mitochondrial sSNVs**
*Variant calling:* An integrated pipeline was developed for the identification of mitochondrial variants within SMaHT donor samples. This workflow combines short-read (Illumina) and long-read (PacBio and ONT) sequencing data across four distinct callers, employing a cross-evidence validation framework to determine confidence tiers. Variant detection was executed at the resolution of individual tissues, donors, and technologies. Utilizing BWA-MEM and minimap2, both short and long reads were aligned to the GRCh38 reference genome. BAM files from diverse Genome Centers were merged to produce high-depth alignment files for downstream analysis. Variant calling was performed on these merged alignments. For short-read cohorts, Mutect2 (mitochondrial mode v1.0) and Mutserve (v2.0.3) were applied to Illumina CRAMs. Long-read processing involved the application of MitoScope (v0.3.0) and Himito (v1.1.2) to PacBio and ONT aligned BAM files.

*Intra-sample merging and normalization:* Per tissue and donor, we performed intra-sample merging by keeping GT and AF in the FORMAT fields. Contig names were standardized to chrM; indels were left-aligned/normalized, atomized, and multiallelic records split. Calls at a known blacklisted site (chrM:3107) were removed, and for short-read callers only PASS variants were retained. Each variant was further classified given three lines of evidence: CrossTech (≥2 technologies), CrossTissue (≥2 tissues from the same donor), and CrossCaller (≥2 callers). Variants were assigned to confidence tiers: HighConf (CrossTech, or CrossTissue and CrossCaller), LowConf (CrossTissue only, or CrossCaller only), or LikelyArtifact.

*Annotation:* Additional annotations included averaged per-technology heteroplasmy frequencies, variant origin (germline if present in all tissues, somatic if partially present), and heteroplasmy status (homoplasmic vs. heteroplasmic, 0.95 VAF threshold). To build a cohort-level callset across all tissue-specific samples from 25 donors, VCFs were merged into one multi-sample VCF with BCFtools, retaining per-sample annotations (Tier, ToolSupport, TechSupport, TissueSupport, AvailableTech, AvailableTissues, allele frequency for Illumina, PacBio and ONT respectively, HeteroplasmyStatus, VariantOrigin). Missing genotypes were filled with 0, and INFO was populated with AF, AN, and AC per unique variant.

**S4.6 Comparison of Somatic Mutation Burden in Tissues Across Multiple Donors**
*Sample Filtering:* To compare pairs of donor-tissue samples and identify individuals with a higher or lower burden of somatic single nucleotide variants (SNVs), we used the somatic SNV callset described above. Samples with more than 5,000 somatic SNVs (all SKSE samples and 2 LIVR samples) were first discarded.

*Outlier detection:* Pairwise comparisons were then carried out on the remaining 355 samples, tissue by tissue across donors. For each test every donor's D was scored as a robust modified z-score against that test's own median and median absolute deviation (MAD): $0.6745 \times (D - \text{median}) / \text{MAD}$. A donor-tissue pair was flagged an outlier at $|z| > 3.5$ and only tests with at least 5 reporting groups were treated as reliable. Three donors were flagged across many different tests (SMHT029 - SKNE, MUSC, BRTL, COAS, CODS, LUNG, HART; SMHT020 -

SKNE, COAS, CODS, AORT, HART, TESL; SMHT040 MUSC, BRFL, BRTL, BRHL, CODS, ADGL) while the majority of the remainder had only single isolated spikes.

### Supplementary Methods Section 5 | Data integration and downstream analyses

#### S5.1 Germline Variant Analyses

*American College of Medical Genetics (ACMG) Secondary Findings:* We used the 81 genes of the American College of Medical Genetics (ACMG) recommended secondary-findings list, version 3.2 [73]. Variant calls were generated from whole-blood samples from each donor using Illumina's Dynamic Read Analysis for GENomics (DRAGEN) software, v4.3.6. We filtered for only variants with an established clinical interpretation: calls were matched to ClinVar [74] and retained only when variants showed pathogenicity without a competing benign or conflicting information, so that variants of uncertain significance were not elevated.

*Germline Mitochondrial Variant Filtering:* Mitochondrial variants were called using four variant callers across three sequencing technologies — Mutect2 [75] and Mutserve [76] for Illumina short-read data, and Himito[77] and MitoScope[78] for PacBio and ONT long-read data. Variants were further classified as homoplasmic or heteroplasmic using a 95% variant allele fraction threshold, and as germline or somatic based on whether they were present across all or only a subset of a donor's available tissues. Mitochondrial variants were annotated based on MITOMAP [79], a comprehensive database of human mitochondrial DNA variation, where variant pathogenicity is classified according to the mitochondrial DNA-specific ACMG/AMP criteria [80].

#### S5.2 Mitochondrial sSNV Circos Plot (Fig. 2d)

Heteroplasmic variants for donor SMHT005 were visualized as a circular genome plot of the rCRS. Variants classified as HighConf with allele frequency >0.001 were extracted from the inter-sample merged VCF. Six tissues were displayed as concentric tracks ordered anatomically (cerebellum, hippocampus, esophagus, lung, fibroblast, and testis), color-coded by the tissue type. Each track shows a kernel density estimate of variant genomic positions with a fixed 200 bp Gaussian bandwidth and circular wrapping so that density is continuous across the origin, preserving the control-region hotspot that spans the D-loop (m.16024–576). Densities were normalized within each tissue to the lane maximum to emphasize positional enrichment rather than absolute call burden; the number of variants per tissue is reported below the plot. An outer ring annotates rCRS gene and control-region features, with protein-coding and rRNA genes labeled by strand, tRNA genes shown as unlabeled ticks, and the D-loop highlighted.

#### S5.3 Functional annotation of somatic mutations (Fig. 3g)

We use PromoterAI[81], SpliceAI[82], APARENT2[83] and AlphaGenome[84] to predict the molecular effects for sSNVs. The effects on gene expression of sSNVs located near promoter regions are predicted by PromoterAI. We downloaded pre-computed scores for all sSNVs within the promoters (±500 bp of the TSS) of all transcripts of protein-coding genes annotated in GENCODE v39 from the PromoterAI GitHub repository (https://github.com/Illumina/PromoterAI). These scores are based on the PromoterAI model, which was trained on strand-specific gene expression and other regulatory signals around the TSS and fine-tuned on rare promoter variants from GTEx across tissues, using paired RNA and genome sequencing data. The predicted scores range from −1 to 1 and reflect the change in expression relative to the mean level at that position, with negative values indicating under-expression and positive values indicating over-expression. We use the 95th percentile of all possible sSNVs in promoters as the cutoff (|PromoterAI_Score| >0.2028) to select the predicted functional sSNVs in our dataset. The splicing effects for sSNVs near canonical splice sites were predicted by SpliceAI. We downloaded pre-computed scores for all possible sSNVs within genes, computed with a maximum variant-to-splice-site distance of 50 bp, from Illumina Basespace (https://basespace.illumina.com/s/otSPW8hnhaZR). This model was trained on canonical junction annotations from GENCODE v24, supplemented with novel junctions observed in GTEx data. For each variant, SpliceAI outputs delta scores representing the change in probability of a site being gained or lost as a donor or acceptor. The overall delta score is the maximum of these four values and is interpreted as the probability that the variant is splice-altering. We used the authors' recommended high-recall threshold of 0.2 to select the predicted functional sSNVs in our dataset.

The polyadenylation effects for sSNVs near polyA-DB[85] canonical polyA sites were predicted by APARENT2. We downloaded the pre-computed APARENT2 scores from the APARENT2 GitHub repository (https://github.com/johli/aparent-resnet) which cover ±100 bp of the 3′-cleavage site. The model was trained on a re-processed HEK293 MPRA of alternative polyadenylation reporters and predicts the log-odds of proximal polyA site usage, with the delta log-odds representing the variant effect. We use the 95th percentile of the effects of all possible single-nucleotide variants on PAS usage as the cutoff (|APARENT2_Score| > 0.5053) to select the predicted functional sSNVs in our dataset.

The tissue-specific gene expression effects for sSNVs was predicted using a local installation of AlphaGenome (all-folds weights and inference code released by Google DeepMind), focusing on the RNA-seq tracks. For the sSNVs from each tissue, we used the RNA-seq tracks of the corresponding GTEx tissue. AlphaGenome scores each gene located within ±1 Mbp of the variant and reports the log2 fold change in predicted gene expression for the alternative allele relative to the reference. We evaluated the absolute raw predicted scores for common variants in each RNA-seq track, as their 95th percentile was approximately 0.2, we used an absolute raw-score cutoff of 0.2 to select the predicted functional sSNVs.
sSNVs that met the criteria for at least one of the four tools were selected as predicted functional sSNVs.

**S5.4 Donor exclusion from the developmental mutation analysis**
The number of candidate developmental mutations across donors showed a bimodal distribution, with nine donors having more than 500 candidates and the remaining donors having fewer than 260. The nine donors were enriched in age over 67 y.o. (p-value = 0.004 by Mann-Whitney U rank test), and the distribution of their candidates across samples revealed hallmarks of clonal hematopoiesis and blood infiltration into other tissues (Fig. S5a.1). Specifically, most candidate developmental mutations had their highest VAF in blood and were present in blood, lung, and multiple other tissues, but not in the brain, which is protected by the blood–brain barrier. In contrast, the distribution of mutations across tissues showed no obvious evidence of clonal hematopoiesis or blood infiltration into other tissues (Fig. S5a.1). Based on this evidence, we excluded the nine donors from the analysis of developmental mutations. Candidate mutations from the remaining sixteen donors represented the final set of developmental mutations.

**S5.5 Cell-type assignment of somatic tandem repeat expansion (Fig. 3b)**
The cell type assignment of somatic tandem repeat is based on the DNA methylation carried on single-molecule long-read. We utilized the SniffCell (v0.9.4) method (https://github.com/Fu-Yilei/SniffCell) to associate the DNA methylation carried on the same read with cell-type differentially methylated regions (ctDMRs). Based on the DNA methylation matching, we observed a recurrent neuron-specific somatic repeat expansion at *CHST8* (chr19:33757354–33757634, AAAG) among four SMaHT donors. The repeat expansion lies near three ctDMRs that effectively distinguish neurons from oligodendrocytes. At chr19:33,758,340–33,758,933, neuronal DNA was hypermethylated (0.82), whereas oligodendrocyte DNA was hypomethylated

(0.18); at chr19:33,763,425–33,764,301, methylation was 0.86 in neurons and 0.28 in oligodendrocytes; and at chr19:33,767,889–33,768,070, methylation was 0.89 in neurons and 0.28 in oligodendrocytes.

For each read and each ctDMR it covered, mean CpG methylation was compared with the neuron and oligodendrocyte reference means, and the read was assigned to the closer methylation state. Evidence from multiple ctDMRs was combined at the read level; when the evidence conflicted, a plurality assignment required at least 66% agreement. Together, these three ctDMRs assigned 83.2% (99/119) of the unique primary reads spanning the repeat in the SMHT020 and SMHT022 frontal-lobe ONT samples: 34 reads were assigned to neurons and 65 to oligodendrocytes. Assigned reads were separated into neuron and oligodendrocyte BAMs before tandem-repeat analysis.

High-confidence neuron-specific expansions were detected in SMHT020 and SMHT022. Inspection of the remaining SMaHT donors identified suggestive neuron-associated repeat instability in SMHT023 and SMHT030; however, each event was supported by fewer than three neuron-assigned reads and therefore did not meet SniffCell's confidence threshold. Overall, four donors showed evidence of neuronal repeat instability at this locus. Somatic reads were determined using stablevizer (https://github.com/ACEnglish/stablevizer) with parameters for minimum reads per-tissue at 2 and germline quantile at 0.1 showed instability in testis from SMHT001.

**S5.6 QC and cell-class annotation for snRNA-seq and snATAC-seq data (Fig. 3h and 3i)**
QC was conducted using an in-house developed computational pipeline. For snRNA-seq, we filtered cells with UMI counts below 500, gene counts below 300, and mitochondria fraction above 5%. We then used SoupX to purify the transcriptome measurement by subtracting the background transcripts. DoubletFinder further detected the potential doublets and retained the clean count feature matrices for singlets. For snATAC-seq, we filtered cells with TSS enrichment below 4 and unique fragment counts below 1000, and filtered doublets using ArchR. For snRNA-seq cell-class annotation, we processed each tissue independently. We used reference-free marker-based cell-class annotation with Harmony as the batch-integration engine. The top 2,000 highly variable genes were scaled and reduced by PCA. Harmony was run on the principal components for multi-donor tissues. We then clustered the data unsupervised at a resolution chosen by a bootstrap-stability sweep, and ranked per-cluster differentially expressed genes, then scored each cluster against a curated marker panel. For snATAC-seq data, we used reference-based label transfer from the matched snRNA-seq annotation. Iterative LSI on the tile matrix and Harmony batch correction were run over all snATAC libraries and then split into different tissues. Cell classes were transferred from the tissue-matched snRNA reference by unconstrained Seurat integration through ArchR, and we used Harmony dimensions with 10,000 variable genes for the final labels. A per-tissue UMAP was recomputed on the inherited Harmony dimensions for visualization.

**S5.7 CpG entropy analysis**
Methylation entropy was calculated from ONT-modified base BAM files using modkit entropy v0.6.1. CpG sites were analyzed with both strands combined. Entropy was calculated from methylation patterns spanning four CpG positions, with the four positions required to occur within a maximum genomic interval of 50 bp. Each CpG position was required to have at least 10 valid reads with each CpG sites' modkit's default confidence filtering was retained. Eight computational threads were used per sample. The representative command was:

```
modkit entropy \
  --in-bam SAMPLE.bam \
  --out-bed SAMPLE.entropy.bed \
```

```
--ref GCA_000001405.15_GRCh38_no_alt_analysis_set.fna \
--cpg \
--num-positions 4 \
--window-size 50 \
--min-coverage 10 \
--threads 8
```

The final entropy dataset contained one entropy BED file for each of 297 ONT samples. Sample identifiers were matched to donor age and detailed tissue annotations using the SMaHT production manifest and ONT metadata. After requiring non-missing age and entropy measurements, the genome-wide analysis included 294 samples from 24 donors, representing 21 detailed tissues and donor ages from 42 to 89 years.

For the genome-wide analysis, the entropy values from all evaluable windows were averaged within each sample to obtain a sample-level mean CpG methylation entropy. This mean value was used to assess age-associated changes within each detailed tissue. For the promoter-restricted analysis, entropy windows were intersected with the ENCODE cCRE v4 annotation using bedtools v2.31.1. Promoters were defined as cCREs annotated as promoter-like signatures (PLS). For each sample, the mean entropy was then calculated across all covered PLS cCREs.

The combined association between donor age and genome-wide methylation entropy was evaluated using a linear regression model that included detailed tissue identity as a categorical fixed effect. When multiple samples were available from the same donor and detailed tissue, their entropy values were averaged, yielding one observation per donor–tissue combination. The analysis included 263 donor–tissue observations from 24 donors and 21 detailed tissues. The model allowed each tissue to have a different baseline entropy level while estimating a common age-associated slope across tissues. Standard errors were clustered by donor to account for correlations among tissues collected from the same individual. The P-value was obtained from a two-sided test of whether the tissue-adjusted age coefficient was equal to zero. The contribution of age was summarized using partial $R^2$, calculated by comparing the full model containing age and tissue identity with a reduced model containing tissue identity alone: partial $R^2$ = (full-model $R^2$ − tissue-only $R^2$)/(1 − tissue-only $R^2$). The full-model $R^2$ was 0.871 and the tissue-only $R^2$ was 0.848, corresponding to a partial $R^2$ of 0.148 for age. After adjustment for detailed tissue identity, donor age was positively associated with genome-wide methylation entropy ($\beta$ = 0.00086 entropy units per year, donor-clustered $P = 1.45 \times 10^{-11}$; partial $R^2$ = 0.15).

**S5.8 SMHT005 blood fraction estimation from ONT methylation (Fig. 4f)**
Blood cell infiltration was estimated for ONT-modified base CRAM files using cell type deconvolution with reference to a DNA methylation atlas.
Read-level methylation counts were generated using wgbstools v0.3.0-7-g6f24bed, (https://github.com/nloyfer/wgbs_tools) using bam2pat, and cell type proportions were estimated for each sample using UXM deconv (https://github.com/nloyfer/UXM_deconv no version; commit 8d0bb45).

With the following representative commands:

```
wgbstools bam2pat \
    --nanopore \
    --genome hg38 \
    --out_dir DONOR \
    SAMPLE.cram
```

```
python UXM_deconv/uxm deconv DONOR/SAMPLE.pat.gz \
        --atlas UXM_deconv/supplemental/Atlas.U25.l4.hg38.tsv \
        --output SAMPLE.uxm_deconv.csv
```

For each sample, the blood infiltration fraction was calculated by summing the UXM-estimated contributions of the atlas cell types: Blood-B, Blood-Granul, Blood-Mono+Macro, Blood-NK, Blood-T.

**S5.9 Recovery correction for somatic mutation profiles from duplex sequencing**

Somatic mutations detected by duplex sequencing may be biased by the genomic territory each assay interrogates, and by context-specific biases in mutation calling. To quantify these biases jointly and to correct the somatic mutation profiles, we used germline heterozygous variants.

For each donor, we identified germline heterozygous sites from independent standard bulk sequencing of blood, retaining variants with a population allele frequency of at least 0.0001 in gnomAD version 4.1 (mutation set B). Because standard bulk sequencing provides near-uniform coverage of germline variants the genome, set B defines the genomic territory over which germline variants are callable as a whole, rather than the territory accessible to any particular duplex assay. Set B is a property of the donor and is expected in every tissue from that donor. With complete recovery across the callable genome, every site in B should therefore also be recovered from duplex reads in every duplex sample from that donor, whichever tissue it came from.

Duplex sequencing was applied to multiple tissues. By default, the analysis workflows for duplex assays are designed to remove germline variants, so we disabled the filtering steps that rely on the matched normal sample and on population databases. Variants were then called independently from the duplex data for each sample and restricted to the germline polymorphisms of that individual, giving mutation set D, a subset of B. Set D is an unbiased readout of mutations recovered the duplex assays, and it is subject to the same constraints that shape the somatic duplex call set from the same sample.

The contrast between sets B and D is the quantity of interest. Each duplex method may cover a different and non-random part of the genome. The ratio of D to B in each context therefore measures the combined loss from restricted territory, insufficient duplex depth, and context-specific calling behavior, expressed relative to what is callable genome-wide.

Sites in B and in D were classified into the six base substitution types in pyrimidine context and, using the immediately flanking bases, into 96 trinucleotide contexts (c). These are the same categories used for mutational signature analysis of the somatic calls.

For each of the 96 contexts, we calculated a sample-specific effective recovery rate:

$s_c = d_c / b_c$

where:

c indexes the mutation context,

$s_c$ is the effective recovery rate in context c,

$d_c$ is the number of germline heterozygous sites recovered by duplex sequencing in that sample (set D), and

$b_c$ is the number identified in blood (set B).

We refer to s as a sensitivity profile for brevity, but it reflects callable territory, duplex depth, and calling sensitivity together.

This yields a sensitivity profile s for every sample. Each context contains a large number of germline heterozygous sites, so the binomial confidence intervals on $s_c$ are narrow relative to the variation in s across contexts. The differences in recovery between contexts are therefore not attributable to sampling noise. We scaled each profile to unit mean,

$$s' = s / \mathrm{mean}(s),$$

and defined the correction factor for each context as

$$f_c = 1 / s'_c.$$

The somatic mutation profile was then corrected as

$$o'_c = o_c \times f_c,$$

where $o_c$ is the observed count of somatic mutations in context c. Because s is scaled to unit mean, the correction adjusts the shape of the profile and leaves the total mutation burden approximately unchanged.

The corrected profiles rescale each somatic spectrum toward the spectrum that would have been observed had the assay recovered mutations across the whole callable genome with equal efficiency in every context. Because every method is expressed on the same genome-wide reference, the corrected profiles are comparable across methods. The approach simultaneously captures the trinucleotide content of the reference bases, and the possible biases caused by the mutated bases.
